# Engineering P4 satellite phage particles with single and dual antimicrobial actions

**DOI:** 10.64898/2026.03.30.715334

**Authors:** Maria Ababi, Matthew Tridgett, Cindy Castado, Normand Blais, Sandra Giannini, Alfonso Jaramillo

## Abstract

Novel strategies for treating bacterial infections are needed to combat the growing threat of antibiotic resistance. Here we engineered P4-like particles for bacterial-culture proof-of-concept experiments in two antimicrobial modes: a single-action design intended to deliver model payloads into bacteria without programmed lysis, and a dual-action design intended to lyse target bacteria while releasing a secondary antigenic payload, exemplified here by model antigens. Using a P2 helper-free P4-like particle production platform, we designed, produced and tested P4-mediated single-and dual-action antimicrobial prototypes. After completing bacterial-culture proof-of-concept experiments, we optimized early-stage bioprocessing for future studies, leading to 10^11^ plaque forming units (PFU) per mL and 0.25 endotoxin units (EU) per 10^9^ PFU. We also challenged the P4 viral-vector packaging limit by deleting *sid* to favour packaging into P2-sized capsids (∼25.8 kb estimated cargo capacity). Importantly, repressing payload expression during particle production improved viral titers by about 2 logs, reduced detected cargo-sequence alterations from 12/20 to 0/20 sequenced post-transduction isolates and enabled higher-dose transduction with stronger detectable antigen signal. Altogether, this study supports P4-derived phage-like particles as antimicrobial prototypes that extend phage function beyond bacterial elimination and provides a basis for future validation in pathogen and animal models of infection.

## 1. Introduction

Antimicrobial resistance presents a growing threat to our global health, prompting research into antibiotic alternatives, including the use of phages—viruses that naturally kill bacteria [1]. Phage therapy has shown promise in several cases of personalized treatments over the past century for difficult-to-treat bacterial infections. In one retrospective series of 100 personalized cases, bacteria were eliminated in 61.3% of cases when phages were used with antibiotics, whereas antibiotics alone had negligible effect [2]. Despite aligned efforts, there are no phage-based products that have yet received marketing authorization through the conventional drug approval process. Major barriers include host-range specificity, pharmacokinetics, dosing, bacterial resistance, purity, manufacturing and regulatory complexity [3], [4], [5]. With advancements in synthetic biology, phages were engineered to follow a clearer path towards translational medicine by removing lysogeny genes, enhancing efficacy, engineering host range and supporting more controlled production [2], [6], [7]. This progress led to several phage-focused companies now advancing through preclinical and clinical trials. Engineered phages are also employed as tools for targeted CRISPR editing, biopanning, affinity-based diagnostics and purification methods [8], [9], [10].

In this study we evaluate engineering strategies that could support future antimicrobial and antigen-delivery applications of P4-like phage particles. We build on our previously reported helper-free production strain, C-5545Δcos-Marionette [11], and re-engineer P4-derived cargo vectors, hereafter P4 phasmids, to produce P4-like particles for the corresponding proof-of-concept experiments. Relative to other phage-delivery platforms, including broad-host-range P1 phagemids, lambda cosmids with larger DNA payloads (∼40-45 kb) and T4 capsids with very high packaging capacity (∼171 kb), the helper-free P2/P4 system was selected here for helper-phage-free, non-replicative bacterial delivery, while accepting smaller standard P4 cargo capacity (∼4 kb), host-range engineering and large-cargo packaging efficiency as design constraints [12], [13], [14]. The production strain supplies P2 structural proteins and prevents P2 genome packaging; the target strain used for the proof-of-concept experiments in bacterial cultures is the P2-nonlysogenic *E. coli* EMG2 (ATCC 23716) unless otherwise stated (Figure 1).

**Fig. 1.**
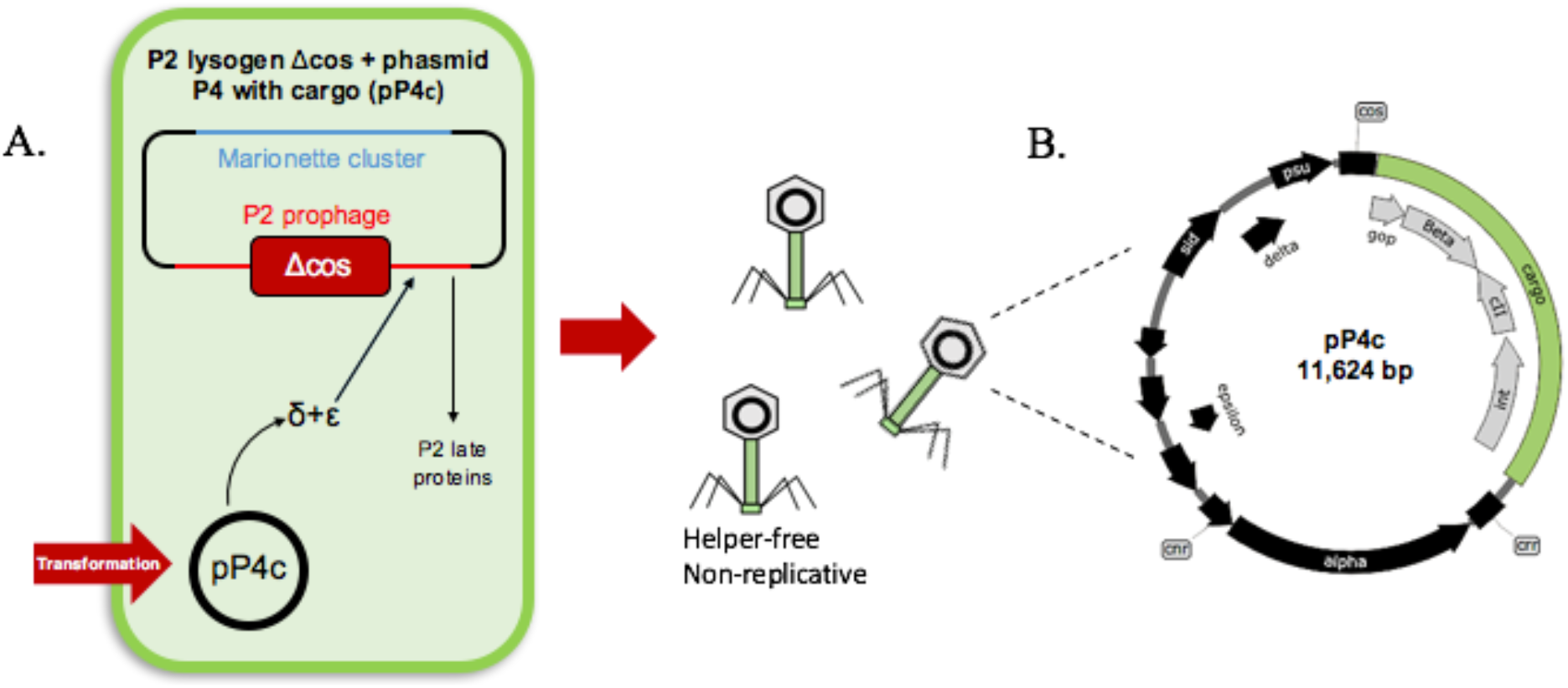
P4-like phage particle production. **A**, P4 production strategy: *Escherichia coli* C-5545 was modified to contain a P2 packaging signal knockout to avoid packaging of the P2 genome during the spontaneous awakening of the P2 prophage (helper phage) [11], and the Marionette cluster [15] to help regulate the heterologous expression. When transformed with P4 phasmids, P2 provides the structural genes for phage assembly [16], while P4 contributes with a series of regulatory elements (e.g. *ɛ* [17], *δ* [18], [19], *psu* [20] and *sid* [21], [22]) that induce the production of late helper proteins and change the morphology of the helper capsid to preferentially pack P4 genomes. **B,** P4 phasmid sequencing map (11,624 bp). Grey coding region = non-essential genes for lytic growth (*gop*, *beta*, *cII* and *int*), flanked by the cos and crr sites essential for packaging and replication of DNA, respectively; Green feature (nucleotide positions 247-4065) = Cargo coding region replacing the non-essential genes for lytic growth. By encoding custom cargo directly onto the P4 phasmid, the production of helper-free phage-like particles encapsulating heterologous cargo becomes replicative in the production strain and non-replicative when delivered to target P2-nonlysogenic strains.

The P2 helper-free P4-like particle platform used here was developed to produce phage-based particles that encapsulate custom payloads [11]. The particles are replicative in the production strain and non-replicative in the target strain. This separation supports controlled bacterial-culture proof-of-concept experiments, while any activity in animal models, regulatory use or applied therapeutic development would require separate validation.

We designed and evaluated two bacterial-culture prototypes: a single-action design for payload delivery without programmed host lysis and a dual-action design that combines host-cell lysis with release of a secondary antigenic product. We varied promoter strengths and ability to repress the prototype payload expression in the production strain (*i.e.* P_lacI_/P_lacI_^q^, P_LtetO1_/P_DiffJJ_, P_tac_, P_VanCC_) to also assess the P4-like particle platform performance. After these bacterial-culture proof-of-concept experiments, we evaluated packaging limits of the viral vectors and early-stage bioprocessing factors as engineering constraints for future targeted antimicrobial candidates.

## 2. Prototype design, optimization and testing in bacterial cultures

### 2.1. Phage-based single-action prototype

The phage-based single-action prototype was designed to test whether P4-like particles could deliver model payloads to *E. coli* EMG2 and yield extracellularly detectable products without programmed host lysis and associated endotoxin release. This bacterial-culture model does not test microbiome behaviour or therapeutic activity (Figure 2).

**Fig. 2.**
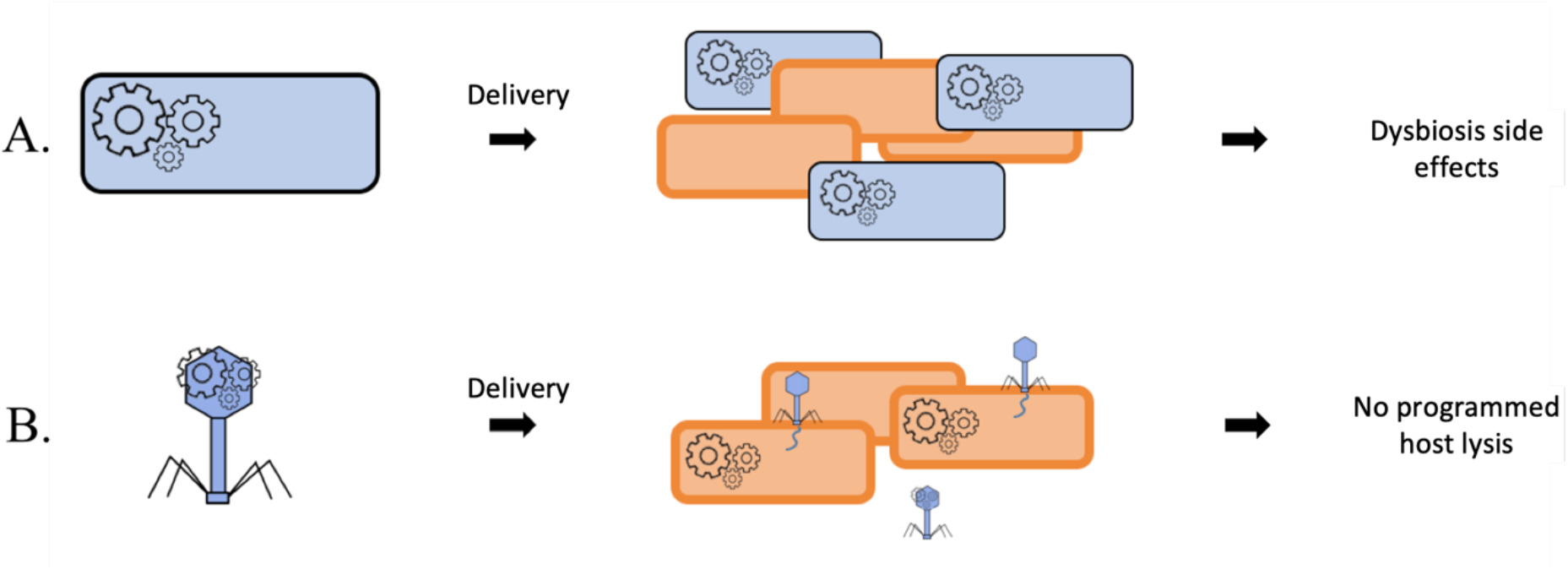
Single-action bacterial delivery compared with engineered probiotics. **A**, Engineered probiotics (blue) can persist alongside other bacteria (orange) after delivery and may alter the bacterial community. **B,** Non-replicative P4-like particles (blue) deliver payload to target bacteria (orange) without programmed host lysis. This schematic summarizes the delivery concept, and microbiome models were not tested.

To demonstrate the single-action antimicrobial concept, we engineered P4 phasmids to encode two Bordetella pertussis model antigens in parallel: a fragment of ADP-ribosyltransferase subunit S1 (p65) and a fragment of bifunctional hemolysin/adenylate cyclase (p72). Both proteins contribute to *B. pertussis* pathogenesis and were chosen as model antigens for bacterial-culture testing of the phage-based single-action prototype [23], [24].

#### 2.1.1. Design of single-action P4 phasmid constructs

P4 phasmid constructs were engineered as listed in Table 1 and as described in Figure 3. Two promoters, P_lacI_^q^ and P_LtetO-1_ with different activity during production, were included in the phasmid design to assess the impact of payload expression during particle production on the produced phage-like particles.

**Fig. 3.**
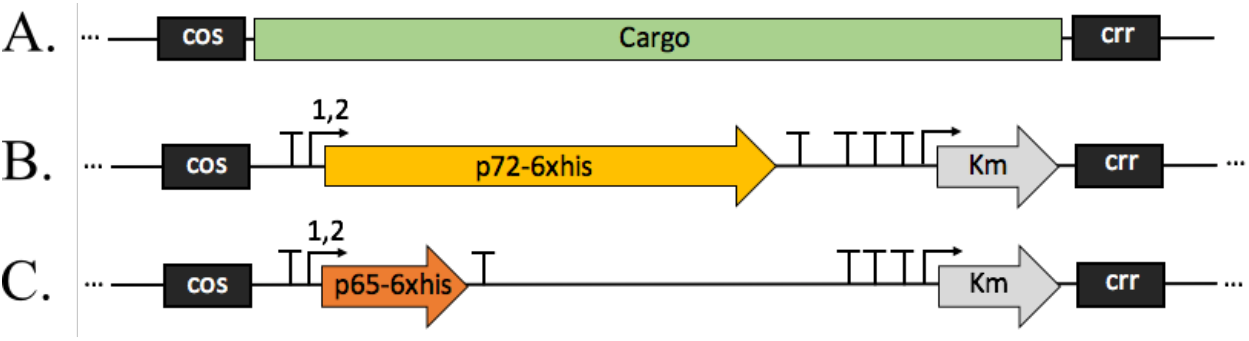
Design of single-action antimicrobials. **A**, pP4 engineered (11,624 bp) green feature = cargo region in nucleotide positions 247-4065. **B,** pP4 phasmid coding for cargo with a part of bifunctional hemolysin/adenylate cyclase, or ‘p72’; **C,** pP4 phasmid coding for cargo with a part of ADP-ribosyltransferase subunit S1, or ‘p65’; both transgenes are controlled by varying promoters illustrated as numbers 1-2. 1 = PlacI^q^; 2 = P_LtetO-1_; “,” = or; cos = P4 packaging signal; “T” = terminator; Km = kanamycin resistance gene; crr = cis required region for replication.

**Table 1.** Single-action antimicrobials phasmid constructs.

| Constructs tested | Model protein | Protein promoter | Protein expression in production strain | Protein expression in target strain | Molecular mass of model protein |
| --- | --- | --- | --- | --- | --- |
| pP4-LacIq-p72 | p72 | P<sub>lacI<sup>q</sup></sub> | constitutive | constitutive | 74 kDa |
| pP4-pLtetO1-p72 | p72 | P<sub>LtetO-1</sub> | repressed | constitutive | 74 kDa |
| pP4-LacIq-p65 | p65 | P<sub>lacI<sup>q</sup></sub> | constitutive | constitutive | 21 kDa |
| pP4-pLtetO1-p65 | p65 | P<sub>LtetO-1</sub> | repressed | constitutive | 21 kDa |

P_LtetO-1_ promoter leads to repressed expression in the production strain containing the Marionette cluster, C-5545Δcos-Marionette, while the P_lacI_^q^ remains constitutive. Both promoters lead to constitutive expression in the target strain, *E. coli* EMG2.

#### 2.1.2. Production of single-action P4-like particles

The designed single-action antimicrobial phasmids in Table 1 were transformed into C-5545Δcos-Marionette to produce in parallel helper-free P4-like particles packing the corresponding phasmid constructs (Figure 4A).

**Fig. 4.**
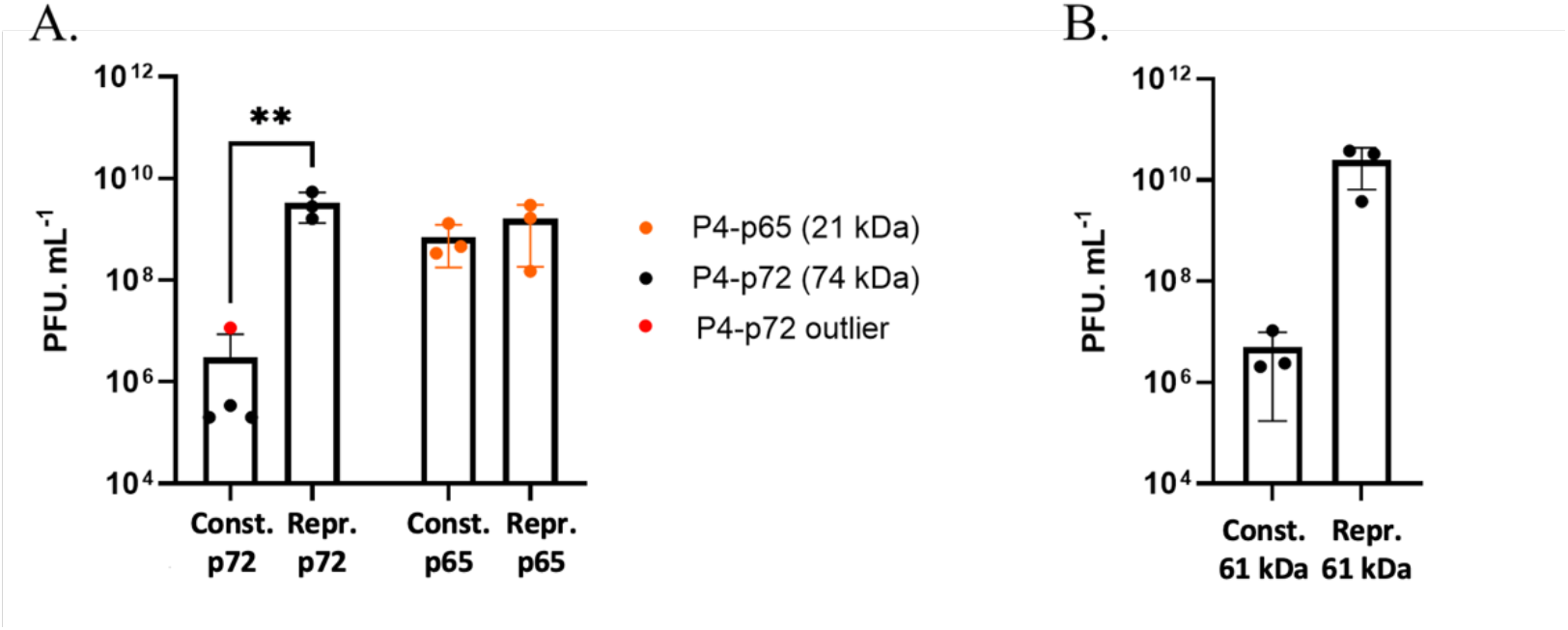
Small-scale production of P4-like particles encoding model proteins. **A**, Titers for particles produced from constitutive p72 phasmid pP4-LacIq-p72, production-repressed p72 phasmid pP4-pLtetO1-p72, constitutive p65 phasmid pP4-LacIq-p65 and production-repressed p65 phasmid pP4-pLtetO1-p65. **B,** Titers for particles produced from constitutive 61 kDa model-protein phasmid pP4-LacIq-Large and production-repressed 61 kDa model-protein phasmid pP4-pLtetO1-Large. Particle labels in the plots omit the leading “p” used for the corresponding phasmid construct names; for example, P4-LacIq-p72 denotes particles produced from phasmid pP4-LacIq-p72. Dots are individual measurements and bars are mean ± s.d. Red dot = ROUT outlier shown in panel A. **P ≤ 0.01. n ≥ 3.

This resulted in ∼10⁹ plaque-forming units (PFU)/mL when using the pP4-pLtetO1-p72 (repressed), pP4-pLtetO1-p65 (repressed) and pP4-LacIq-p65 (constitutive) phasmids. The pP4-LacIq-p72 phasmid coding for the 74 kDa protein expressed constitutively during production resulted in a significantly lower titer at about 10⁶ PFU/mL, with one outlier at 10⁷ PFU/mL.

To test whether the size of p72 and not its function had a negative impact on the production titer when constitutively expressed, two other P4 phasmids were engineered coding for a different 61 kDa protein regulated by the same promoters. The designed phasmids, pP4-LacIq-Large (constitutive) and pP4-pLtetO1-Large (repressed), were used to produce P4-like particles and exhibited a similar pattern as the earlier tested pP4-LacIq-p72 and pP4-pLtetO1-p72 constructs on the production titer: pP4-LacIq-Large construct resulted in about 3 logs lower titers compared to the repressed construct during production, pP4-pLtetO1-Large (Figure 4B). This supports the inhibitory effect of constitutively expressing a larger protein during the production of P4-like particles, likely due to exceeding a certain burden threshold on the production cells. A non-expressed length-matched control would be required to distinguish expression burden from DNA length.

#### 2.1.3. Bacterial-culture evaluation of non-lytic payload delivery

The P4-like particles were evaluated for post-transduction growth kinetics in *E. coli* EMG2 and for delivered protein expression in concentrated supernatants (Figure 5). The initial test for expression and extracellular detection was carried out from pET-24b plasmids transformed and induced in *E. coli* BL21-DE3 (Figure 5A). These showed the following SDS-PAGE migration patterns: between 22-25 kDa for p65; between 100-135 kDa for p72 (Figure 5A, blue and orange arrows, respectively). p72 migrates slower on the gel potentially due to its acidic isoelectric point at 3.9, as commonly observed for other acidic proteins [25].

**Fig. 5.**
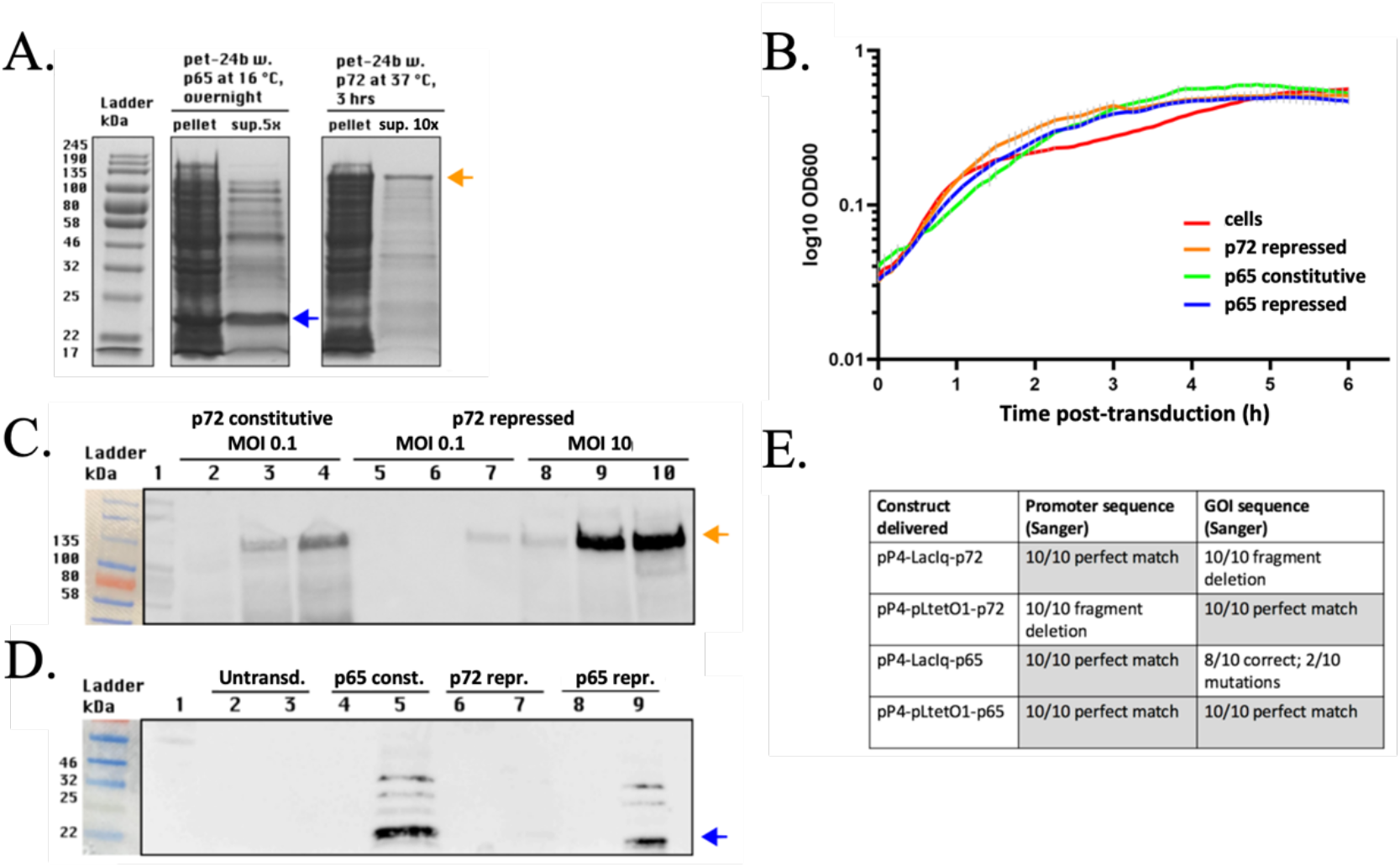
Single-action antimicrobial protein detection in. **E.** coli EMG2 culture supernatants. Blue arrows indicate p65; orange arrows indicate p72. **A,** SDS-PAGE expression controls from pET-24b vector in BL21-DE3 cells after 1 mM IPTG induction. **B,** EMG2 growth after P4-like particle delivery at MOI 10. **C,** p72 Western blot of 20× concentrated supernatant. Lane 1 = ladder. Lanes 2-4 = p72 constitutive (P4-LacIq-p72). Lanes 5-7 = p72 repressed (P4-pLtetO1-p72) at MOI 0.1. Lanes 8-10 = p72 repressed at MOI 10. Within each group, lanes show 3 h, 6 h and overnight. **D**, p65/p72 Western blot of 2× concentrated supernatant. Lane 1 = ladder. Lanes 2-3 = untransduced. Lanes 4-5 = p65 constitutive (P4-LacIq-p65). Lanes 6-7 = p72 repressed (P4-pLtetO1-p72). Lanes 8-9 = p65 repressed (P4-pLtetO1-p65). Paired lanes show 3 h and 6 h. **E,** Sanger sequencing summary for promoter and GOI regions from 10 post-transduction colonies.

The P4-like particles produced in Figure 4A were then used to deliver the respective packed phasmids with p72/p65 cargo into *E. coli* EMG2 (Figure 5B-D). Because the P4-LacIq-p72 stock was low titer, this construct was evaluated at MOI 0.1 for protein detection and is not included in the MOI 10 growth-kinetics panel. As expected, these particles showed no lytic activity at MOI 10 during 6 h post-transduction monitoring. Supernatants were analysed by Western blot with the anti-5xHis-HRP antibody to detect the expected migration patterns for p65 and p72 (blue and orange arrows, respectively). Extracellular p65 and p72 products were detected at 6 h post-transduction when regulated by either promoter: p65 at MOI 10 for PlacIq and PLtetO-1; p72 at MOI 0.1 for PlacIq and at MOI 0.1 or 10 for PLtetO-1. At the same protein and MOI, PlacIq generated a stronger detected signal than PLtetO-1 (p65 at MOI 10; p72 at MOI 0.1). Higher MOI increased the detected signal, but for p72 this comparison was only feasible with the PLtetO-1 construct because PlacIq-p72 production titer was low. These data show detection in concentrated supernatants and do not establish an active secretion mechanism.

Transduced constructs were also evaluated by Sanger sequencing of promoter and GOI regions (Figure 5E; Supplementary Table S5 and Supplementary Figures S2.2 and S2.3). The coding-region results from 10 isolated transduced phasmids per construct demonstrated the negative impact of transgene expression during production of P4-like particles on transgene coding-region sequence integrity. Both constructs that expressed antigen constitutively during the production of particles resulted in modifications in the transgene coding region post-transduction in the target strain: 10/10 deletions (frame shift) for pP4-LacIq-p72 and 2/10 mutations (mixed trace) for pP4-LacIq-p65. Importantly, no mutations/deletions were observed in the transgene coding region when using the P_LtetO-1_ promoter for which the antigen expression was repressed during production. The promoter-region fragment deletion detected for pP4-pLtetO1-p72 is reported separately in Figure 5E and Supplementary Figure S2.3. These observations are consistent with selection for deletion or rearrangement of burdensome cargos during production, although the specific molecular mechanism was not resolved.

Together, the pP4-pLtetO1-p65 and pP4-pLtetO1-p72 constructs supported a bacterial-culture proof-of-concept for the single-action prototype. Repressing payload expression during particle production improved particle yield and preserved the sequence integrity of the single-action payload coding region delivered into the target strain. Functional assays for p65 and p72 were not performed in this study and will be required before therapeutic activity can be inferred.

### 2.2. Phage-based dual-action prototype

The phage-based dual-action prototype was designed to test whether bacterial lysis could be coupled to release of a secondary antigenic product. This design could in future support combined bacterial killing and local antigen release, but unlike the non-lytic single-action design it is expected to release endotoxin when applied to Gram-negative targets (Figure 6).

**Fig. 6.**
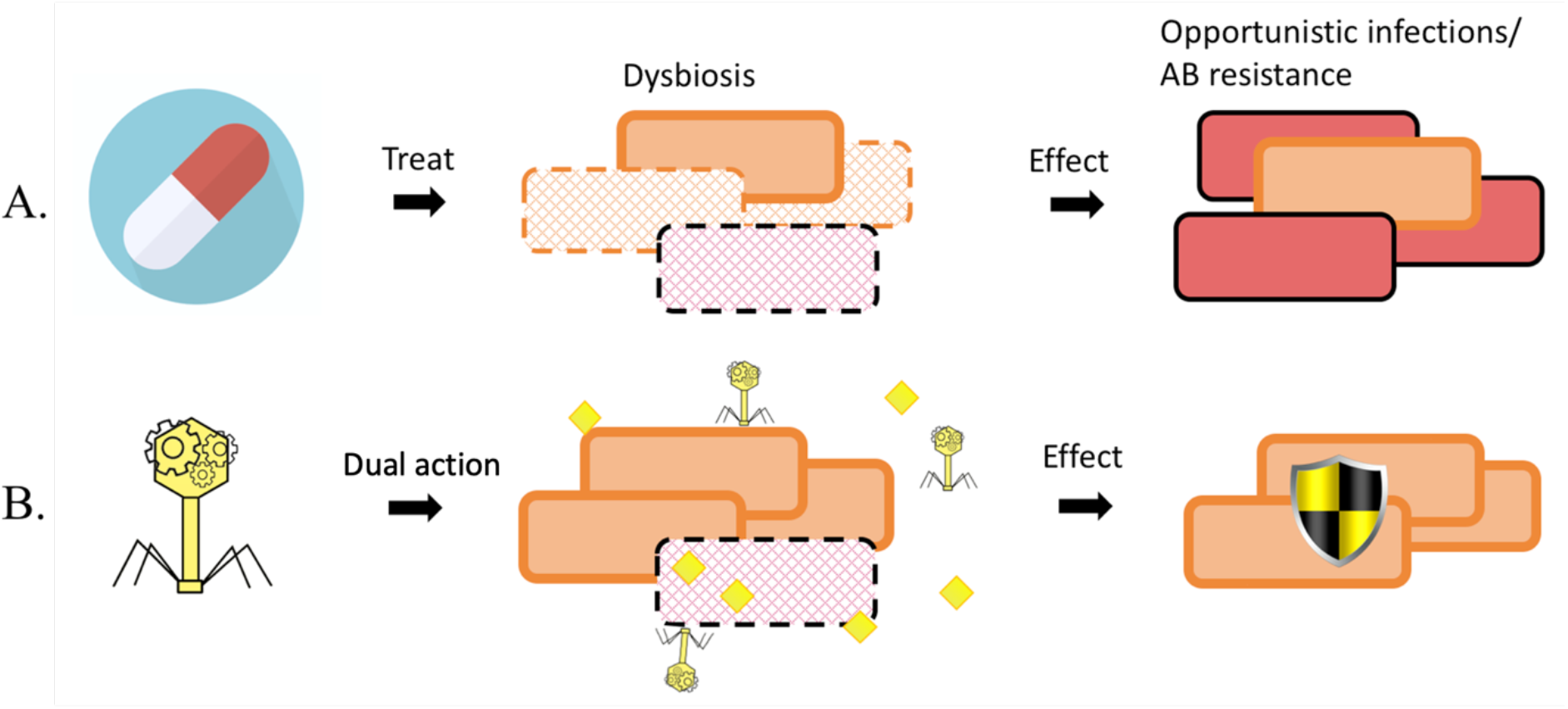
Conceptual lysis-coupled antigen release compared with antibiotic treatment. **A**, Antibiotic treatment can kill target pathogenic bacteria (red) but may also affect neighbouring non-target bacteria (orange). **B,** P4-like particles (yellow) conceptually deliver dual-action cargo to a target bacterium to combine bacterial killing with antigen release (yellow diamond). This schematic is conceptual, and pathogen, microbiome and animal-model validation remains future work. AB = antibiotics.

To demonstrate the dual-action antimicrobial concept, we engineered the P4 phasmid to differentially express the functional lytic cassette described previously [11], [26] and outer membrane protein E (PE) from a nontypeable *Haemophilus influenzae* (NTHi) strain [27], [28]. PE contributes to NTHi pathogenesis [29]. As a model antigen, recombinant wild-type PE is not secreted and was therefore used for bacterial-culture testing of the phage-based dual-action prototype.

#### 2.2.1. Design of dual-action P4 phasmid constructs

P4 phasmid constructs were engineered as listed in Table 2 and as described in Figure 7. Combinations of different promoter strengths for the lytic cassette (PLtetO-1 and its weaker JJ variant) and antigen expression (PlacI and its stronger variants, PlacIq and Ptac) were included in the phasmid designs to assess simultaneous expression kinetics of both the lytic cassette and antigen before cell lysis in the target strain, *E. coli* EMG2.

**Fig. 7.**
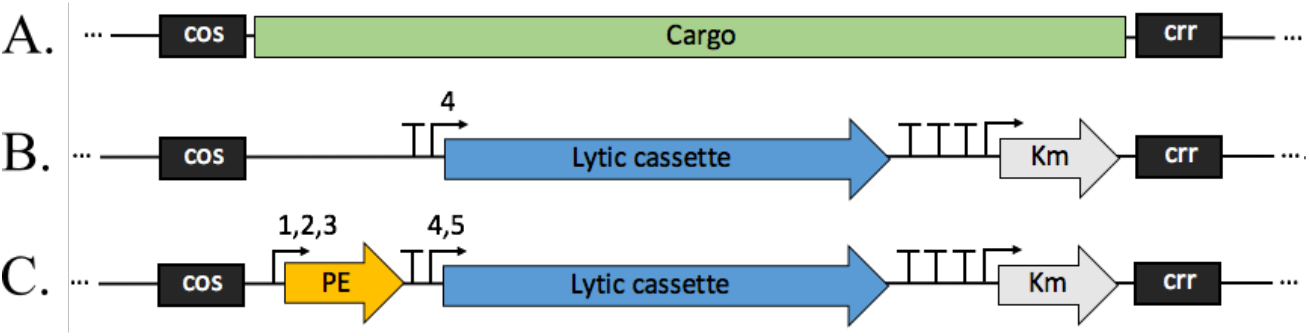
Design of dual-action antimicrobials. **A**, pP4 engineered (11,624 bp) green feature = cargo region in nucleotide positions 247-4065. **B,** pP4-DiffJJ-Test cargo region containing the kanamycin antibiotic resistance gene and a lytic cassette containing: *E. coli* phage MS2 gpL lysis protein, *E. coli* phage Phi X174 gpE lysis protein and Lambda LysS, LysR and Rz lysis proteins. **C,** pP4 phasmid with dual-action cargo encoding the PE antigen and the lytic cassette, controlled by varying promoters illustrated as numbers 1-5. 1 = P_lacI_^q^; 2 = P_lacI_, weaker variant of P_lacI_^q^; 3 = P_tac_; 4 = JJ weaker variant of P_LtetO-1_ (further referred to as P_DiffJJ_); 5 = P_LtetO-1_; “,” = or; cos = P4 packaging signal; “T” = terminator; Km = kanamycin resistance gene; crr = cis required region for replication.

**Table 2.** Dual-action antimicrobials phasmid constructs and lytic control. N/A = not applicable.

| Constructs tested | Lytic cassette | Protein of interest (POI) | Lytic cassette promoter | POI promoter | Lysins expression in production strain | POI expression in production strain | Lysins and or POI expression in target strain |
| --- | --- | --- | --- | --- | --- | --- | --- |
| pP4-DiffJJ-Test | yes | N/A | P<sub>LtetO-1</sub> JJ variant | N/A | repressed | N/A | constitutive |
| pP4-DiffJJ-LacI | yes | PE | P<sub>LtetO-1</sub> JJ variant | P<sub>lacI</sub> | repressed | constitutive | constitutive |
| pP4-pLtetO1-LacIq | yes | PE | P<sub>LtetO-1</sub> | P<sub>lacI</sub><sup>q</sup> | repressed | constitutive | constitutive |
| pP4-pLtetO1-tac | yes | PE | P<sub>LtetO-1</sub> | P<sub>tac</sub> | repressed | repressed | constitutive |

P_LtetO-1_, its weaker variant JJ and the P_tac_ promoters are repressed in the production strain containing the Marionette cluster, C-5545Δcos-Marionette, while P_lacI_ and P_lacI_^q^ promoters are constitutive. All promoters lead to constitutive expression in target strain, *E. coli* EMG2.

#### 2.2.2. Production of dual-action P4-like particles

The designed dual-action antimicrobial phasmids in Table 2 were transformed into C-5545Δcos-Marionette to produce in parallel helper-free P4-like particles packing the corresponding phasmid constructs (Figure 8).

**Fig. 8.**
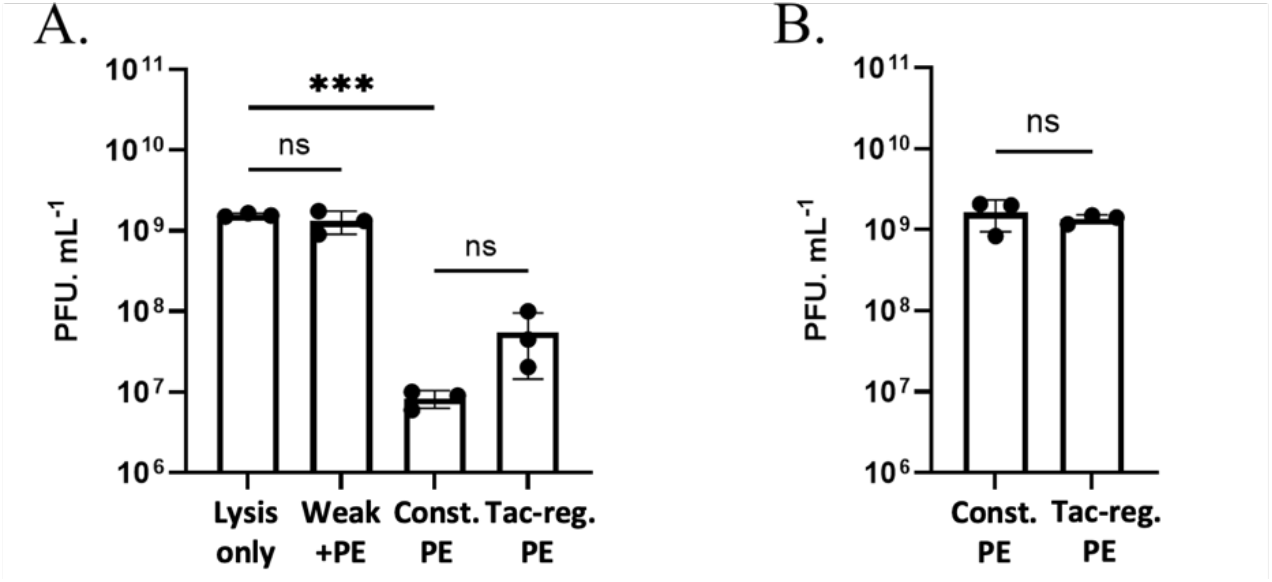
P4-like particle titer for dual-action cargo. **A**, First-round small-scale production from transformed C-5545Δcos-Marionette cells. **B,** Second-round 100 mL production from concentrated panel A particles. Labels show particles produced from the lysis-only control pP4-DiffJJ-Test, weaker-lysis plus PE construct pP4-DiffJJ-LacI, constitutive-PE construct pP4-pLtetO1-LacIq and tac-regulated-PE construct pP4-pLtetO1-tac. Dots are individual measurements and bars are mean ± s.d. Pns > 0.05. ***P ≤ 0.001. n = 3.

This resulted in about 10⁹ PFU/mL titers when using the pP4-DiffJJ-Test (lysis-only) lytic-only phasmid and the pP4-DiffJJ-LacI (dual weak/PE) dual-action phasmid with weaker promoters. A lower titer was observed for the two phasmid constructs carrying the PLtetO-1 lysis promoter, stronger than the DiffJJ variant in the design context (Figure 8A). To reach the needed titers for the next step, the latter two constructs with low titer were concentrated and used as seed stock for a second round of production (Figure 8B). The difference in the first round of production between the constructs could be due to the additional burden on the cells when expressing cargo regulated by stronger promoters leading to either constitutive (PlacIq) or leaky repression (Ptac) protein expression. Repressing PE expression during first-round production using Ptac improved titer compared with PlacIq, albeit non-significantly, likely due to its higher basal expression.

#### 2.2.3. Bacterial-culture evaluation of lysis-coupled antigen release

The P4-like particles were tested for differential expression of the lytic cassette and PE antigen after transduction into *E. coli* EMG2. In this target strain, the lytic cassette and PE are expressed constitutively; therefore, PE antigen accumulation before cell lysis is required for a dual-action readout.

First, the P4-like particles were evaluated for lytic activity by transducing into *E. coli* EMG2 at a range of MOIs from 1 to 10 (Figure 9).

**Fig. 9.**
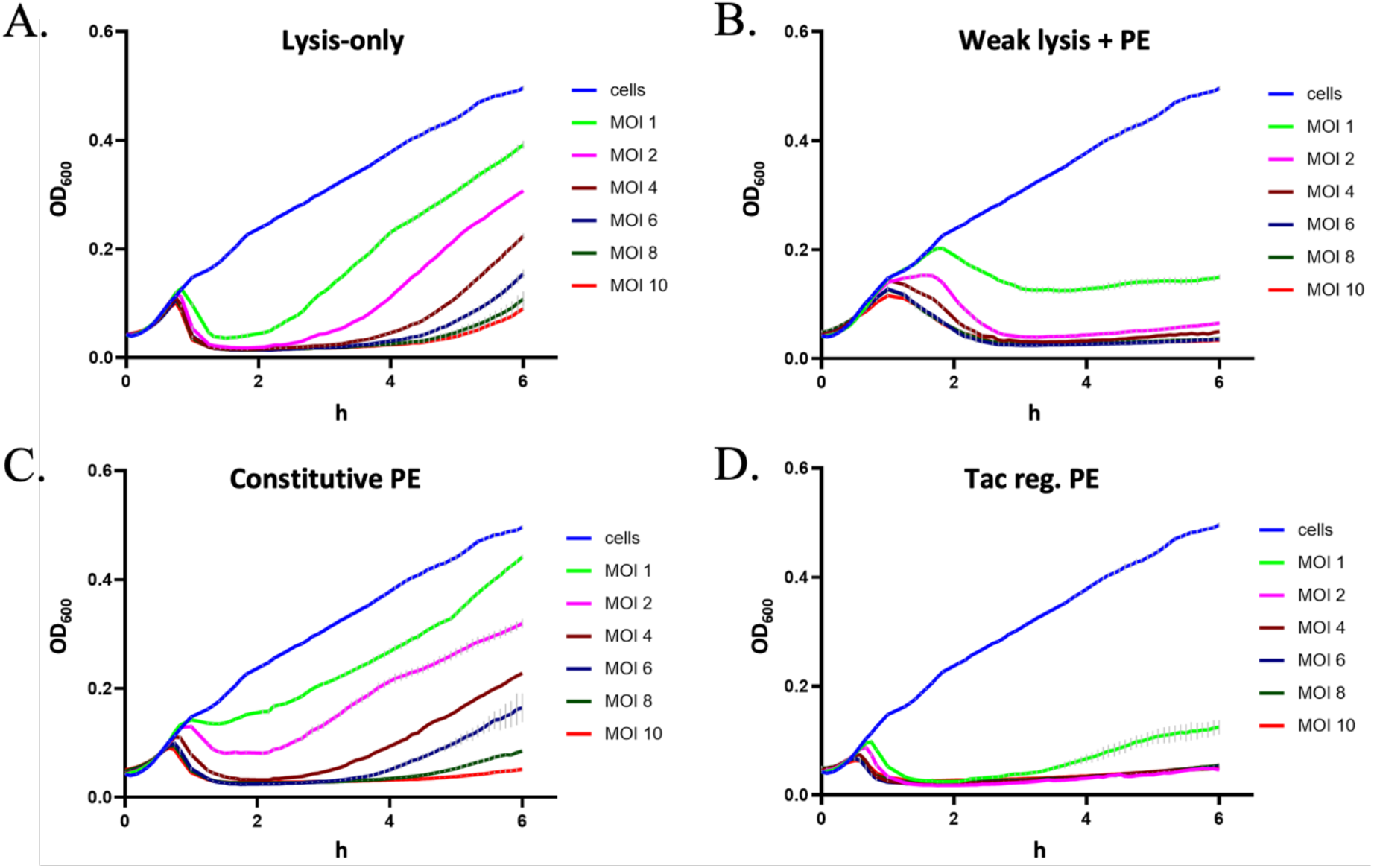
Post-transduction EMG2 lysis kinetics. EMG2 cells were mixed with P4-like particles at MOI 1-10 starting at OD600 = 0.2. **A,** Lysis-only control (P4-DiffJJ-Test). **B,** Weaker lysis plus PE (P4-DiffJJ-LacI). **C,** Constitutive PE with PLtetO-1 lysis (P4-pLtetO1-LacIq). **D,** Tac-regulated PE with PLtetO-1 lysis (P4-pLtetO1-tac). OD600 was normalized to dilution medium containing LB plus 80 mM MgCl2. n = 3.

Transduced constructs exhibited lytic activities at all MOIs and showed distinct lysis profiles (Supplementary Figure S3.2). Compared to the test lytic-only particles, P4-DiffJJ-Test (lysis-only), the P4-DiffJJ-LacI (dual weak/PE) resulted in about 1 hr delayed lysis at lower MOIs, consistent with an effect of additional constitutive expression of PE. Repressing or reducing PE expression during production was associated with faster lysis profiles for P4-pLtetO1-tac, in which PE is tac-regulated, and for P4-DiffJJ-LacI, in which the lysis cassette uses the weaker DiffJJ promoter, compared with P4-pLtetO1-LacIq, in which PE is constitutively expressed from PlacIq and the lysis cassette is controlled by PLtetO-1. Because lysis kinetics integrate promoter activity, particle titer, construct context, burden and lysis-cassette expression thresholds, these profiles should not be interpreted as a direct ranking of promoter strength alone.

The accumulation of PE before cell lysis was tested by evaluating the relative change in the supernatant of all constructs at 50% and 100% lysis. Different MOIs were selected for assessment due to their different lytic profiles (Figure 10).

**Fig. 10.**
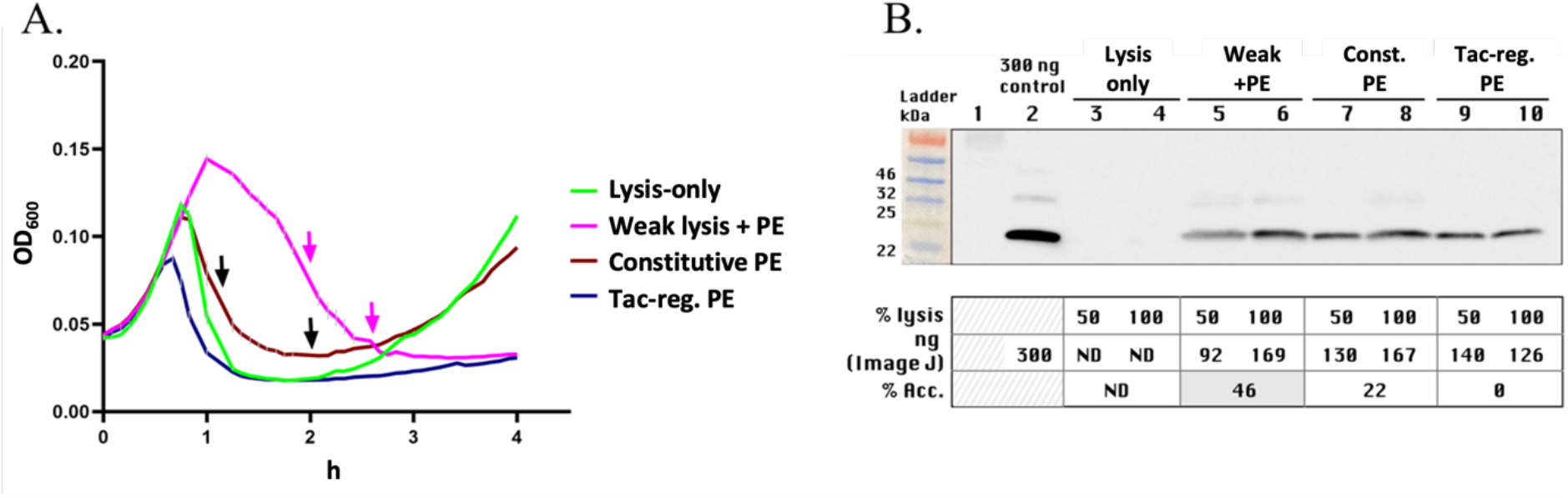
Differential expression in dual-action P4-like particles. **A**, EMG2 lysis kinetics for particles produced from lysis-only control pP4-DiffJJ-Test, weaker-lysis plus PE construct pP4-DiffJJ-LacI, constitutive-PE construct pP4-pLtetO1-LacIq and tac-regulated-PE construct pP4-pLtetO1-tac. Arrows mark Western blot sampling at perceived half-time and full-time lysis. **B,** PE Western blot from panel A supernatants. Lane 1 = ladder. Lane 2 = 300 ng PE control. Lanes 3-4, 5-6, 7-8 and 9-10 correspond to panel A in the same order. “% lysis” = perceived half-time (50%) and full-time (100%) lysis. “% Acc.” = ImageJ band-density estimate relative to the 300 ng PE control from the blot shown. ND = not detected. Original/source blots and source-analysis panels for panel B are presented in Supplementary Figures S4.7-S4.10.

PE antigen was detected in all post-transduced phasmids coding for PE at 50% and 100% lysis. For P4-DiffJJ-LacI, the proportional increase in antigen signal in the supernatant relative to lysis extent, from approximately 92 ng PE at half-time lysis to approximately 169 ng antigen at full-time lysis (semi-quantitative analysis by ImageJ; Supplementary Figure S3.3), was consistent with PE production before complete lysis and subsequent release into the supernatant. Hence, P4-DiffJJ-LacI supported the dual-action bacterial-culture proof-of-concept.

Because P4-pLtetO1-tac at MOI = 2 resulted in more antigen at half-time lysis than the other constructs, P4-pLtetO1-tac particles could be prioritized for follow-up antigen-release and lysis testing.

### 2.3. Expanding cargo size through *sid* deletion for large dual-action constructs

After removal of the non-essential genes for P4 replication (i.e. pP4-DiffJJ-Test), the P4 phasmid can accommodate about 4 kb of custom cargo in that region. Classic studies showed that P4 *sid* mutants fail to direct small P4 capsid assembly and instead produce P2-sized capsids containing multimeric P4 DNA, while later structural work established Sid as an external scaffold that redirects P2 capsid assembly from T=7 to T=4 geometry [21], [22], [30], [31]. Deleting *sid* could therefore increase P4 cargo capacity from approximately 4 kb to an estimated 25.8 kb, creating the possibility of more complex therapeutic approaches.

FliC is a flagellin protein encoded by the *fliC* gene from *Salmonella typhimurium* [32] and can be used as an adjuvant to improve the immunogenicity of vaccine antigens [33], [34], [35]. To accommodate larger cargo that includes the fused adjuvanted-antigen FliC-PE and the lytic cassette, we engineered the P4 phasmid with a *sid* knockout to bias packaging of the larger dual-action cargo into P2-sized capsids. Further, we tested the performance of the helper-free P4-like platform with *sid* knockout in bacterial cultures when challenged with larger cargo by assessing production titer and corresponding post-transduction readouts.

#### 2.3.1. Design of *sid*-deleted large-cargo P4 phasmid constructs

P4 phasmid constructs were engineered with *sid* knockout as listed in Table 3 and as described in Figure 11. The initial screening for fused FliC-PE variants was carried out from pET-24b plasmids transformed and induced in *E. coli* BL21-DE3 (Supplementary Figure S4.1, Supplementary Table S6 and Supplementary Figure S4.2). Note that all FliC-PE variants tested were mostly insoluble. FliCΔ188-346-PE expressed the most in the host cells and was thus chosen for further engineering in pP4 and bacterial-culture testing.

**Fig. 11.**
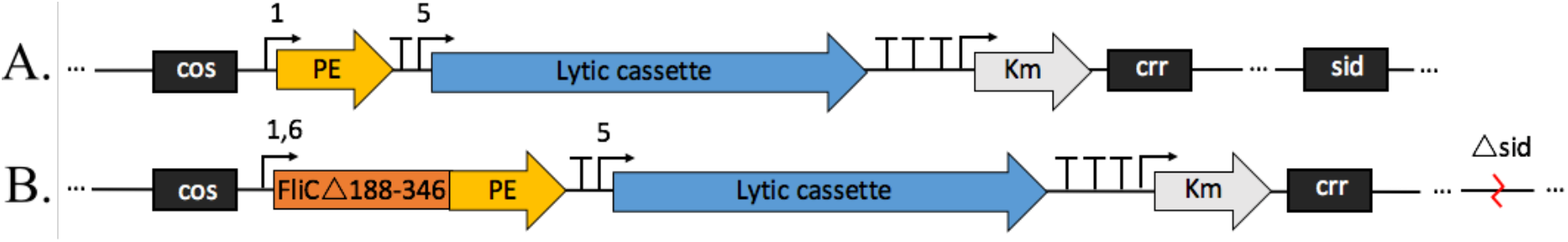
Design of large dual-action antimicrobials. **A**, P4 packaging size control with dual-action cargo encoding the PE antigen and the lytic cassette; **B,** Large dual-action test construct coding for P4 phasmid with *sid* gene deletion (except the 4 bp at the *sid* 3’ end) together with large cargo encoding the fused adjuvanted-antigen, ‘FliCΔ188-346-PE’, and the lytic cassette under the control of varying promoters numbered 1, 5, 6. 1 = P_lacI_^q^; 5 = P_LtetO-1_; 6 = P_VanCC_; “,” = or; cos = P4 packaging signal; “T” = terminator; Km = kanamycin resistance gene; crr = cis required region for replication.

**Table 3.** Large cargo dual-action antimicrobial constructs.

| Constructs tested | Lytic cassette | Protein of interest (POI) | Lytic cassette promoter | POI promoter | Lysins expression in production strain | POI expression in production strain | Lysins and or POI expression in target strain |
| --- | --- | --- | --- | --- | --- | --- | --- |
| pP4Δ*sid*-pLtetO1-LacIq | yes | FliCΔ188-346-PE | P<sub>LtetO-1</sub> | P<sub>lacI</sub><sup>q</sup> | repressed | constitutive | constitutive |
| pP4Δ*sid*-pLtetO1-pVanCC | yes | FliCΔ188-346-PE | P<sub>LtetO-1</sub> | P<sub>VanCC</sub> | repressed | repressed | constitutive |

The constructs with *sid* knockout were designed to include two different strong promoters to regulate the fused adjuvanted-antigen and assess the impact of *sid* deletion on P4-like platform performance: PlacIq and PVanCC. PVanCC leads to repressed expression during particle production in the C-5545Δcos-Marionette strain, while PlacIq is constitutive. All promoters lead to constitutive expression in the target strain, *E. coli* EMG2.

#### 2.3.2. Production of *sid*-deleted large-cargo P4-like particles

The designed dual-action antimicrobial phasmids with large cargo in Table 3 were transformed into C-5545Δcos-Marionette to produce in parallel helper-free P4-like particles packing the corresponding phasmid constructs (Figure 12A).

**Fig. 12.**
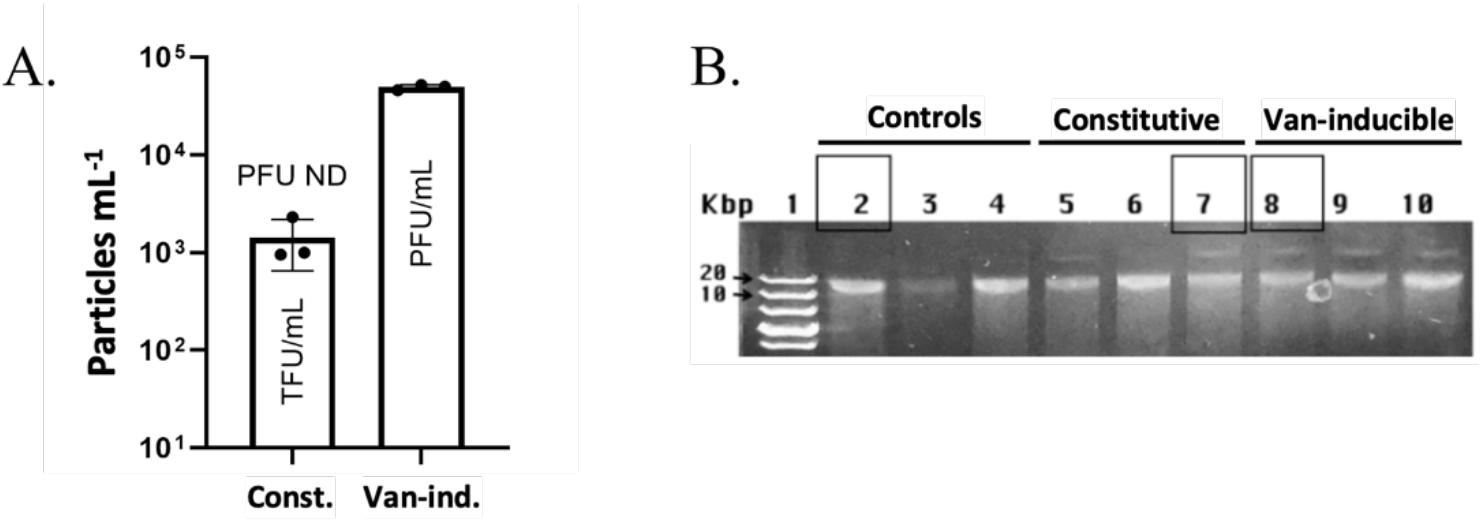
Production and isolation of sid-deleted large-cargo P4-like particles. **A**, Output for particles produced from Δ*sid* constitutive large-cargo phasmid pP4Δsid-pLtetO1-LacIq and Δ*sid* Van-inducible large-cargo phasmid pP4Δsid-pLtetO1-pVanCC, both carrying the lytic cassette and FliCΔ188-346-PE. **B,** Agarose gel of kanamycin-selected phasmids in *E. coli* Marionette. Lane 1 = ladder. Lanes 2-4 = standard control (pP4-pLtetO1-LacIq). Lanes 5-7 = constitutive. Lanes 8-10 = Van-inducible. Black squares = selected lanes. PFU = plaque-forming particles. TFU = antibiotic-resistant transductants. ND = not detected. n = 3.

This resulted in ∼1 × 10³ TFU/mL when using the pP4Δ*sid*-pLtetO1-LacIq (Δ*sid* constitutive) construct and ∼5 × 10⁴ PFU/mL when using the pP4Δsid-pLtetO1-pVanCC (Δ*sid* Van-inducible) construct. The lower titer relative to pP4 without *sid* knockout could be due to inefficient packaging into helper-free P2-sized capsids and would require further optimization. The terms PFU and TFU are used to report different operational readouts: PFU measures particles that form plaques on the production strain, whereas TFU measures antibiotic-resistant target cells that successfully received the phasmid.

Due to the low yield for direct transduction experiments in *E. coli* EMG2, cargo functionality was further investigated indirectly by transduction into *E. coli* Marionette strain. The initial screen for successful transfer of *sid*-deleted constructs was based on enlarged phasmid size as a sign of concatemerization (here dimers) attempting to fill the larger capsid. Upon transduction into the host cell, the packed concatenated dimer is likely to return to its monomeric state, as observed for other multimeric plasmids [36]. This could explain the two bands observed on the agarose gel for the transduced larger cargo constructs (Figure 12B). The *sid*-intact phasmid carrying PE and the lytic cassette, pP4-pLtetO1-LacIq (standard-size), was used as a negative control for concatemerization and a positive control for the following indirect differential-expression studies. The gel pattern is consistent with multimeric phasmid forms, but it does not directly determine capsid size, packaged DNA length, copy number or particle homogeneity.

#### 2.3.3. Induced lysis and payload-expression testing of *sid*-deleted constructs

Transduced *E. coli* Marionette isolates containing separately the pP4-pLtetO1-LacIq (standard-size), pP4Δ*sid*-pLtetO1-LacIq (Δ*sid* constitutive) and pP4Δ*sid*-pLtetO1-pVanCC (Δ*sid* Van-inducible) were tested for induced lysis and constitutive or induced payload expression. The lytic cassette regulated by PLtetO-1 was induced with anhydrotetracycline (aTc), while the adjuvanted antigen regulated by the PVanCC promoter was induced with vanillic acid (Van). Supplementary Figure S4.4 confirmed that single and combined inducers were not toxic under these conditions and did not interfere with differential expression.

First, the transduced phasmids were evaluated for growth kinetics and lytic activity when induced with aTc (Figure 13).

**Fig. 13.**
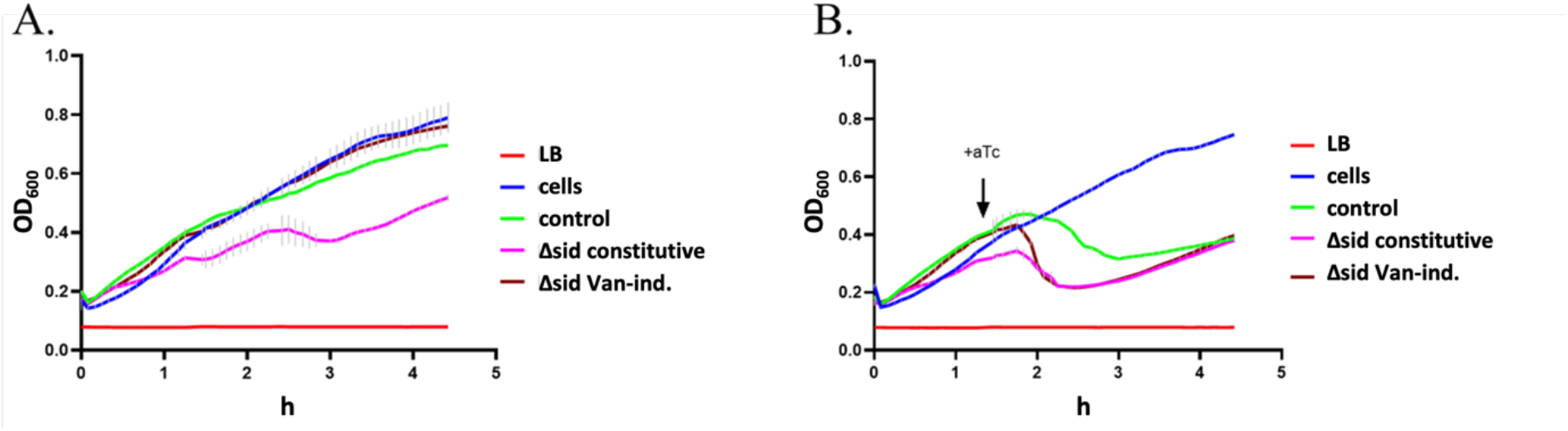
Induced lysis kinetics for large dual-action antimicrobials. **A**, Non-induced control. **B,** aTc-induced lysis. Curves show LB, E. coli Marionette, the standard-size PE control pP4-pLtetO1-LacIq, the Δ*sid* constitutive FliCΔ188-346-PE construct pP4Δsid-pLtetO1-LacIq and the Δ*sid* Van-inducible FliCΔ188-346-PE construct pP4Δsid-pLtetO1-pVanCC. n = 3.

The Δ*sid* constitutive FliCΔ188-346-PE construct (pP4Δsid-pLtetO1-LacIq) showed host cell growth inhibition (Figure 13A). Nonetheless, all constructs exhibited lytic activity when induced with aTc confirming lysis-cassette functionality. Interestingly, the same lysis cassette showed faster lysis in the transduced pP4Δ*sid*-pLtetO1-LacIq or pP4Δ*sid*-pLtetO1-pVanCC constructs compared to the transduced pP4-pLtetO1-LacIq construct (Figure 13B). This is consistent with concatemeric cargo carrying additional lytic-cassette copies after transfer into Marionette cells.

Further, the differential expression was examined by inducing lysis and protein expression with both aTc and Van, respectively (Figure 14A).

**Fig. 14.**
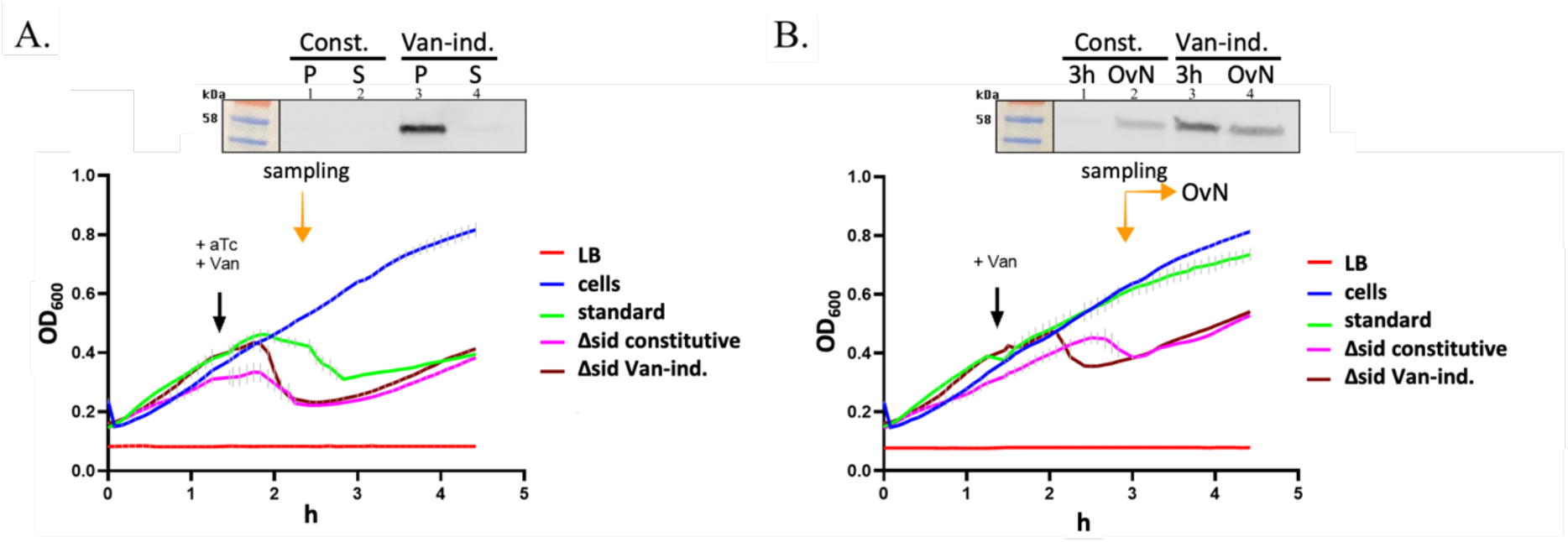
Induced lysis and FliC-PE expression in large-cargo constructs. **A**, aTc plus Van induction. Inset PE blot shows Δ*sid* constitutive (P4Δsid-pLtetO1-LacIq, lanes 1-2, P and S) and Van-inducible (P4Δsid-pLtetO1-pVanCC, lanes 3-4, P and S). **B,** Van-only induction. Inset PE blot shows the same constructs at 3 h and overnight (OvN). Curves show LB, E. coli Marionette, the standard-size PE control pP4-pLtetO1-LacIq, the Δ*sid* constitutive FliCΔ188-346-PE construct pP4Δsid-pLtetO1-LacIq and the Δ*sid* Van-inducible FliCΔ188-346-PE construct pP4Δsid-pLtetO1-pVanCC. Orange/yellow = sampling. Black = induction. OvN = overnight. n = 3. Original/source blots for panels A and B are presented in Supplementary Figures S4.11 and S4.12.

Fused adjuvanted-antigen was detected at 50% lysis in the pellet of Marionette cells transduced with pP4Δsid-pLtetO1-pVanCC (Δsid Van-inducible) only. FliCΔ188-346-PE from transduced pP4Δsid-pLtetO1-pVanCC was not detected in the supernatant as expected due to its insoluble nature (Supplementary Figure S4.2B). When allowed to express overnight, with Van only (Figure 14B), FliCΔ188-346-PE was confirmed to express also from transduced pP4Δsid-pLtetO1-LacIq (Δsid constitutive), albeit in smaller quantities compared to pP4Δsid-pLtetO1-pVanCC. The modest growth reduction observed without aTc induction in Figure 14B is consistent with basal lysis-cassette leakiness and/or stress from large-cargo expression in Marionette cells; therefore, the induced assays were interpreted qualitatively.

This supports detectable expression from the large-cargo constructs generated by *sid* deletion after transduction into Marionette recipient cells. Repressing expression of the secondary antigenic product during particle production resulted not only in increased titer but also in improved post-transduction protein expression (Figure 14B).

### 2.4. Bioprocessing

#### 2.4.1. Upstream optimization

Improving P4-like particle titer is essential for future preclinical evaluation of the constructs and later for manufacturing. Here we optimized P4-like particle production using the pP4-DiffJJ-Test phasmid to achieve a titer of (5.2 ± 0.34) × 10⁹ PFU/mL in 100 mL production volume before concentration (Figure 15).

**Fig. 15.**
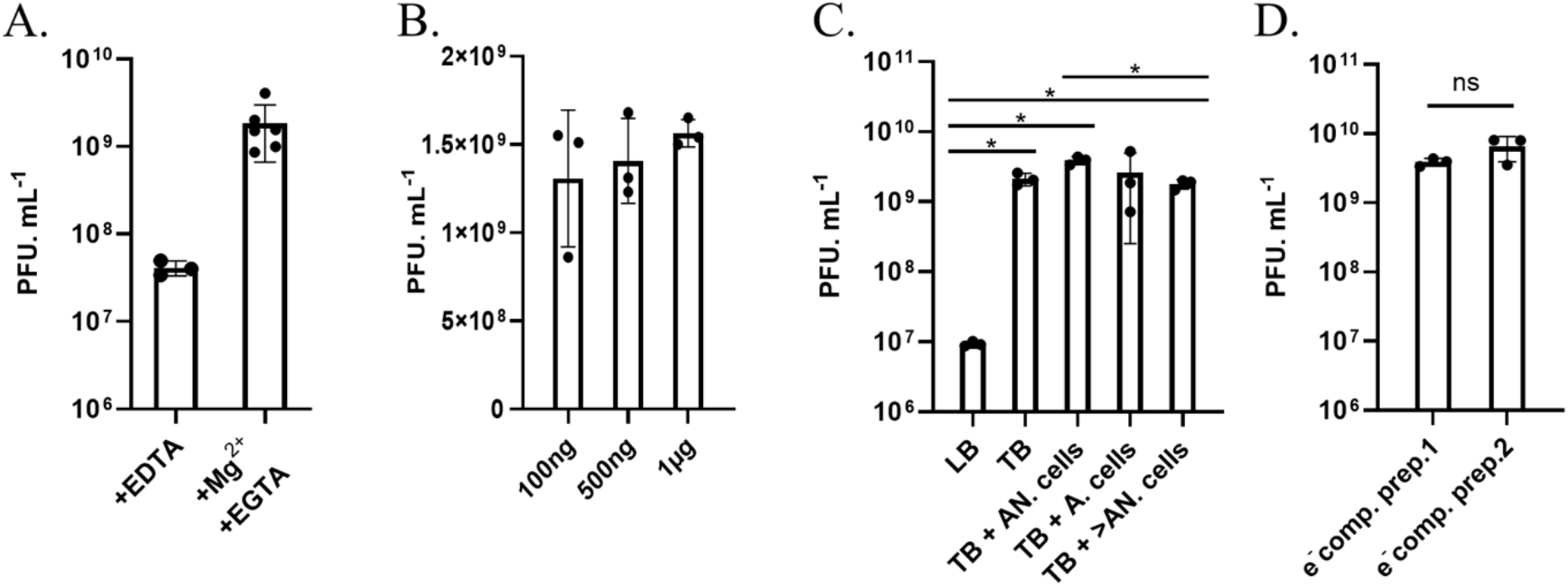
Upstream bioprocessing. **A**, Small-scale production (15 mL) with and without MgCl2 supplementation. EGTA was used (instead of EDTA) to avoid chelating Mg2+ while still chelating Ca2+ to prevent attachment to cell debris at the end of the production protocol; **B,** Small-scale production (15 mL) started with varying phasmid amount; **C,** Larger-scale production (100 mL) with media variations; 100 mL total volume; LB = Luria Bertani broth; TB = Terrific Broth; TB+AN.cells = Terrific Broth supplemented at the start with 2.5 mL anaerobic C-5545Δcos-Marionette cells grown overnight, stationary; TB+A.cells = same as TB+AN.cells, but the cells were grown aerobically; TB+>AN.cells = same as TB+AN.cells, but 10 mL of cells was added instead; **D,** Electrocompetent-cell variability between preparation lots on different days reflected in particle production using the same method as in panel C TB+AN.cells. Pns > 0.05; *P ≤ 0.05; (n ≥ 3).

The critical parameters identified were: addition of viral-particle stabilizers towards the end of the production (Mg^2+^ with EGTA), use of production media rich in nutrients (Terrific Broth) and addition of anaerobically grown cells for optimal particle-cell interactions (Figure 15A, C). The reproducibility of the titer yields was improved with increasing amount of starting phasmid and was demonstrated from two electro-competent cells production batches (Figure 15B, D).

#### 2.4.2. Downstream optimization

To concentrate the viral particles 100 times, we evaluated the following methods: PEG6000/NaCl precipitation and ultrafiltration with 100 kDa (UF100) and 300 kDa (UF300) cut-offs (Figure 16A). There was no statistical difference in titer yields between the concentration methods. By concentrating the crude lysate 100 times via the UF100 method, we achieved the highest average of ∼1.71 × 10¹¹ PFU/mL, with 90.4% recovery.

**Fig. 16.**
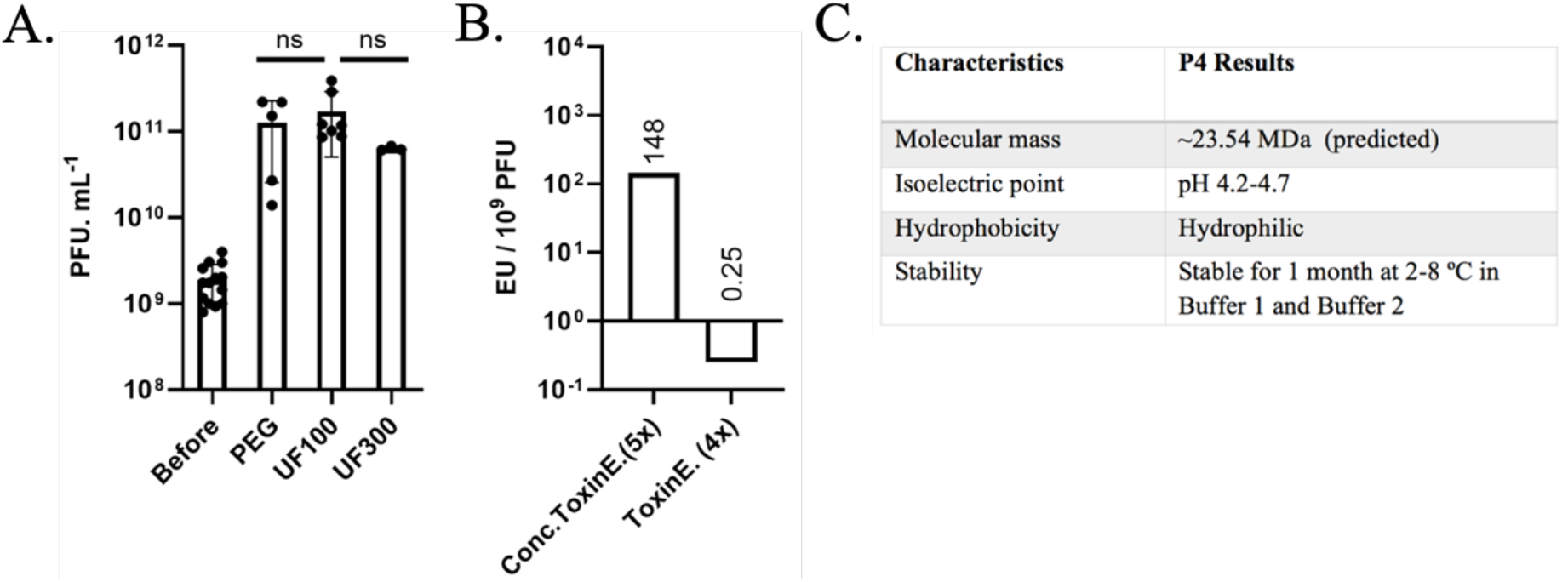
Downstream bioprocessing. **A**, 100× concentration methods; Before = titer of phage-like particles before concentrating; PEG = concentrating by PEG6000/NaCl precipitation; UF100 = concentrating by Vivaspin ultrafiltration with a cut-off of 100 kDa; UF300 = concentrating by Vivaspin ultrafiltration with a cut-off of 300 kDa. **B,** Endotoxin purification via ToxinEraser starting with concentrated phage stock at 10¹¹ PFU/mL and separately with phage stock at 10⁹ PFU/mL. **C,** P4 particle characteristics. Buffer 1 = 1× PBS, 20 mM MgSO4, pH 7.0; Buffer 2 = 20 mM Tris, 127 mM NaCl, 2 mM CaCl2, 5 mM MgSO4, pH 7. Pns > 0.05; (n ≥ 3).

The P4-like particles were evaluated for isoelectric point, mass, hydrophilicity, stability and solvent tolerance for tailored removal of endotoxins (Figure 16C; Supplementary Table S1 and Supplementary Figure S1.1). Endotoxins are bacterial lipopolysaccharides (LPSs) that can trigger toxic shock above threshold exposures. They are released upon gram-negative cell lysis and are predominant in phage stocks prepared using gram-negative production hosts. P2-derived particles use LPSs as receptors for cell attachment and thus may form particle-LPS complexes when in free solution [37].

For future preclinical planning, we evaluated the endotoxin removal methods applicable to the P4-like phage [38], [39], [40], [41], [42], [43], [44] (Supplementary Figure S1.4). We achieved the most significant endotoxin removal using the ToxinEraser kit resulting in 0.25 EU/10⁹ PFU with 30% particle recovery (Figure 16B). Under the specific example dose assumptions used here (e.g. 20 g mouse; <20 EU/mL oral route; <1 EU/mL parenteral route, 100 μL injection), this endotoxin level would fall within published endotoxin limits for oral and parenteral administration except intrathecal administration [38], [45]. However, the titer is marginally low at 6 × 10⁸ PFU/mL and 0.15 EU/mL considering the non-replicative attribute of the phage-like particles in the target strain. This would require optimization and/or upscaling before any animal-model experiments.

#### 2.4.3. Manufacturing upscaling feasibility

To prepare for future preclinical manufacturing tests, the technology was successfully transferred to a specialized contract manufacturing and research organization, Jafral Biosolutions. Upon upscaling to a 900 mL bioreactor, Jafral achieved a titer of 3 × 10⁹ PFU/mL before concentration. Upon two rounds of downstream bioprocessing, Jafral removed endotoxins down to 0.28 EU/10⁹ PFU, in a final concentrated stock of 1.07 × 10¹¹ PFU/mL with 30 EU/mL. Under the same dose assumptions, the required titer dilutions are within the number of non-replicative particles previously observed to produce a detectable signal in animal models [46]. Thus, the more concentrated final pure stock achieved at Jafral Biosolutions allows future dose optimization to inform preclinical proof-of-concept experiments.

## 3. Discussion

Engineering phage to deliver therapeutic cargo is an established concept, with current phage therapy research in preclinical and clinical trials. Recent examples include phage-mediated bacterial base editing in the mouse gut, CRISPR-Cas antibacterial delivery, tail-engineered P2 delivery into gut pathogens and sustained protein production in the mammalian gut [47], [48], [49], [50]. Here we demonstrated in bacterial cultures P4-like phage prototypes for antimicrobial functions beyond bacterial killing. We used a helper-free P4-like particle platform because it separates production from target delivery and allows non-replicative delivery into P2-nonlysogenic target strains. The broader host range remains inferred from published P2-derived particle data, because the present experiments were performed in laboratory *E. coli* strains. P2-derived particles can attach and inject DNA into *E. coli*, Klebsiella, Salmonella, Serratia, Shigella, Rhizobium and Pseudomonas [16], [51], [52]. When desirable, host range specificity could be modulated by swapping the tail fibers to produce chimeric particles as described in Yosef *et al.* 2017 [53] or by using strain-specific promoters that regulate the therapeutic cargo. This work also uncovers platform limitations and lessons that could inform further designs for phage-based antimicrobials.

Repressing cargo expression during phage-like particle production improved titer, preserved payload coding-region integrity and supported post-transduced cargo expression (i.e. pP4-pLtetO1-p72 vs pP4-LacIq-p72). The degree of impact was dependent on promoter strength, promoter leakiness and the size of the regulated therapeutic protein. Thus, production systems that allow cargo repression during production with constitutive expression in the target strain should be considered for improving the quality of phage-based antimicrobials.

The standard P4-sized vector used here has a cargo capacity of about 4 kb. We challenged this limitation by deleting the *sid* gene and testing engineered P4 phasmids expected to use P2-sized capsids. The resulting transductants and agarose-gel patterns were consistent with enlarged, likely multimeric phasmid forms and with an estimated cargo capacity of 25.8 kb, but at the expense of titer. We did not directly measure capsid size, packaged DNA length, copy number or particle homogeneity. The titer could be further optimized by adjusting the phasmid size and production conditions for improved packing efficiency into P2-sized capsids. As a result, the phasmid amount may also need to be optimized to account for potential loss in transformation efficiency.

P4-like phage stocks were difficult to purify using known phage-specific endotoxin purification methods (Supplementary Figure S1.4). This may reflect particle affinity to LPSs, forming particle-LPS complexes after bacterial lysis. Non-concentrated stocks resulted in the purest endotoxin-to-particle ratio when using the ToxinEraser kit at 0.25 EU/10⁹ PFU, but with 30% particle recovery. This was similar to the purity obtained at Jafral Biosolutions, 0.28 EU/10⁹ PFU, but with a concentrated phage stock of 1.07 × 10¹¹ PFU/mL, sufficient to prepare material for further animal-model dose-ranging studies under specified assumptions but not a general safety claim. In addition, endotoxin removal from manufactured stocks does not address endotoxin released by deliberate lysis of Gram-negative target bacteria in the dual-action design. The purity could be improved by transferring the production platform into cells containing LPSs genetically modified as non-toxic, e.g. ClearColi cells. Since the P2-derived particles may not infect cells with modified LPSs, the production method may require optimization to compensate for single production phage cycles.

Taken together, the work should be interpreted as a bacterial-culture engineering proof of concept with defined limitations. First, the experiments relied on laboratory *E. coli* strains, and activity, host range and payload delivery therefore require future validation in clinically relevant pathogens. Second, the *sid*-deletion evidence remains indirect because we did not perform TEM or packaged-DNA copy-number assays. Third, p65, p72 and PE were detected by sequencing or Western blot readouts, but functional antigen assays were not performed. Fourth, the dual-action strategy inherently releases endotoxins because it lyses Gram-negative target cells, so endotoxin exposure, microbiome disturbance, bacterial-load reduction and antigen-specific responses will need assessment in pathogen-based assays and animal models of infection. Within this scope, the constructs with repressed cargo expression during phage-like particle production (single-action: pP4-pLtetO1-p65 and pP4-pLtetO1-p72; dual-action: pP4-DiffJJ-Test, pP4-pLtetO1-tac and pP4Δ*sid*-pLtetO1-pVanCC) are the most promising candidates for further testing because they showed superior titer, payload coding-region integrity and detectable payload expression. After these validation studies, the P4-like platform could also be explored in future as part of a screening workflow for vaccine-candidate evaluation regardless of secretion, supporting future drug-discovery workflows.

Together, the single-and dual-action prototypes support two design routes for P4-like particles: payload delivery without programmed host lysis and bacterial killing coupled to release of a secondary antigenic payload. Here, antigenic model proteins provided experimentally tractable payloads; future designs could test other payload activities once payload function, target-pathogen activity, microbiome impact and endotoxin exposure are evaluated in appropriate pathogen and animal models. Repressing payload expression during particle production improved titre, preserved payload coding-region sequence integrity and enabled higher-dose transduction, identifying a practical design rule for P4-like particle development. The *sid*-deletion experiments further suggest a route to larger cargoes, although direct measurement of capsid size, packaged DNA length, copy number and particle homogeneity remains required.

## 4. Materials and Methods

### 4.1. Strains, plasmids and phasmids

Strains, plasmids and phasmids used in this study are listed in Table MM 1.

**Table MM1.** Strains, plasmids and phasmids.

| Name | Origin |
| --- | --- |
| EMG2 | ATCC 23716 |
| DH5α-Z1 | Expressys |
| DH10B-Marionette | Addgene (Kit #1000000137) [15] |
| BL21-DE3 | Thermofisher |
| C-5545Δcos-Marionette | Produced in AJ group [11] |
| pP4-DiffJJ-Test (pAJ1046) | Produced in AJ group [11] |
| Model protein gene constructs in pET-24b vectors | GSK, Rixensart, BE |

Constructs original to this study are listed in Table MM 2.

**Table MM2.**
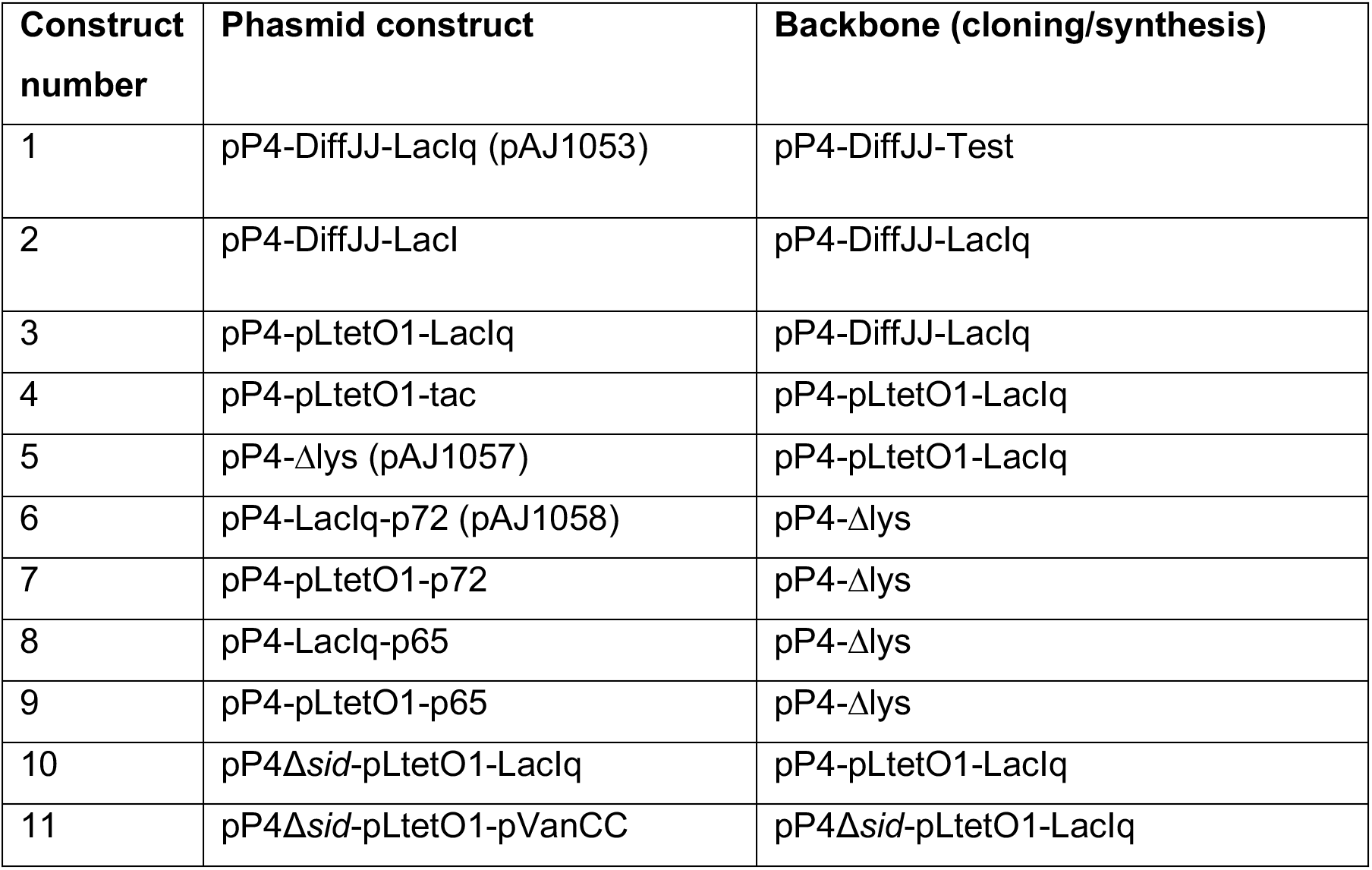
Engineered constructs original to this study.

The constructs were built by Genewiz synthesis and Gibson Assembly. Phusion High-Fidelity PCR 2x Master Mix (Thermo Scientific) and NEBuilder HiFi DNA Assembly were used following the manufacturer’s instructions. Primers and gBlocks were synthesised via Integrated DNA Technology. Sequencing information is provided in Supplementary Tables S9-S22.

### 4.2. Producing high-titer helper-free phage-like particles (first round)

Varying amounts of viral DNA (100 ng, 500 ng or 1 μg) were transformed by electroporation into 100 μL electrocompetent *E. coli* C-5545Δcos-Marionette. Immediately after electroporation, 1 mL of room-temperature Luria Bertani broth (LB) was added to the culture inside the electroporation cuvette and transferred to an Eppendorf tube. The cells were allowed to recover for 2 h at 37 °C with vigorous shaking (200 rpm). The recovered culture was supplemented with 28 μL of 1 M CaCl2 and mixed. The recovered, CaCl2-supplemented culture was then used for small-scale or larger-scale production, as follows:

For small-scale production, the recovered CaCl2-supplemented culture was added to a 150 mL Erlenmeyer flask containing 14 mL of terrific broth with 15 μg/mL trimethoprim. The culture mix was incubated at 37 °C with aeration and slow shaking (130 rpm) for 3 h or until clumps of debris were visible. To prevent particles from attaching to bacterial debris and to maintain particle stability, 0.28 mL of 4% EGTA (pH 8.8) and MgCl2 up to 80 mM were added. The culture was then incubated at 37 °C with aeration for an additional 1.5 h to complete the final lytic cycles. The lysed cells were pelleted by centrifugation at 3,428 rcf for 8-10 min. The supernatant containing the desired phage particles was collected and clarified by 0.2 μm syringe filtration. The lysate was then stored at 4 °C.

For larger-scale production, the recovered culture was added into a 1000 mL Erlenmeyer flask containing 100 mL total volume of terrific broth with 15 μg/mL trimethoprim. In the next steps the added components were adjusted proportionally. To account for the additional interstitial space between bacteria and phage-like particles in the larger volume, we also supplemented the terrific broth with varying amounts in the flask with none, 2.5 mL or 10 mL of *E. coli* C-5545Δcos-Marionette grown overnight (from one colony inoculation) at 37 °C stationary, aerobically or anaerobically.

### 4.3. Producing high-titer helper-free phage-like particles (second round)

Lysates generated in the first production round with a titer below 10⁹ PFU/mL were further concentrated down to 100 µL by Vivaspin ultrafiltration with 100 kDa cut-off, as recommended in the Vivaspin Sartorius manufacturer’s instructions. The concentrated lysates were used as feed material to begin a second round of production, as follows:

A 2.5 mL LB culture containing 15 μg/mL trimethoprim was inoculated with one colony of the C-5545Δcos-Marionette production strain in a 15 mL culture tube and vortexed briefly. The culture was incubated at 37 °C without aeration (with the dual-position cap in the down position), stationary, for 20 h overnight. The next day, the culture was supplemented with up to 2 mM CaCl2 and mixed. The culture with CaCl2 was then fed with a total of ∼1.8 × 10⁸ particles from the first round of production, briefly mixed and incubated stationary at 37 °C for 10 min.

Meanwhile, 200 μL 1 M CaCl_2_ were mixed with 800 μL LB in a 1.5 mL Eppendorf tube and vortexed well. The mixture was transferred to a 1000 mL Erlenmeyer flask containing 96.5 mL terrific broth supplemented with 15 μg/mL trimethoprim, drop-wise and whilst swirling the destination flask. To this flask, the culture with particles was transferred after 10 minutes incubation and allowed to grow at 37 °C with aeration, slowly shaking (130 rpm) for 3 hours (or until clumps of debris were visible). The culture was then supplemented with 2 mL of 4% EGTA (pH 8.8) and MgCl_2_ up to 80 mM and then incubated at 37 °C with aeration for an additional 1.5 hours. The culture was next transferred to two 50 mL falcon tubes and centrifuged at 3,428 rcf for 8-10 minutes. The resulting supernatants were collected and further clarified using a 500 mL 0.22 μm filtration system. The clarified lysates containing helper-free P4-like particles were then stored at 4 °C until further use.

### 4.4. Plaque-forming unit assay

4 mL of LB supplemented with 15 μg/mL trimethoprim were inoculated with a single colony of production strain, C-5545Δcos-Marionette. The culture was incubated overnight at 37 °C with vigorous shaking, 200 rpm. Next day, the overnight culture was refreshed in 10 mL LB with 15 μg/mL trimethoprim and grown until OD600 = 0.2-0.3, shaking at 37 °C. Meanwhile, dilutions of the lysate were prepared in LB and stored at room temperature until further use. 50 mL LB agar (LBA) plates supplemented with 5 mM CaCl_2_ were also prepared using large square petri-dishes. The exterior of the LBA bottom plates was drawn with a marker to form a 6 x 6 grid.

900 μL of 0.2-0.3 OD600 cells and 6 μL 1 M CaCl_2_ were mixed in a 15 mL culture tube, and incubated stationary at 37 °C for 10 minutes. 9-10 mL of 42-46 °C soft TBA (1:1 mix of LBA and terrific broth) were added to the culture tube and gently mixed. The content of the culture tube was poured onto a large LBA plate with CaCl_2_, to form an evenly distributed bacterial lawn. Immediately after, 10 μL of each lysate dilution was dropped on the surface of the bacterial lawn, in individual squares marked on the grid. The drops were allowed to become dry at room temperature to avoid the drops from merging. The plates were then flipped up-side-down and incubated at 37 °C, overnight. The plaques for each dilution contained in marked squares were counted next day and multiplied by 100x the dilution factors to calculate the phage-like particles per mL.

### 4.5. Transducing-forming unit assay

4 mL of LB supplemented with 50 µg/mL spectinomycin were inoculated with a single colony of target strain, DH5α-Z1. The culture was incubated overnight at 37 °C with vigorous shaking, 200 rpm. Next day, the culture was refreshed in 10 mL LB with 50 µg/mL spectinomycin and grown shaking at 37 °C until OD600 = 0.2-0.3. Meanwhile, the dilutions of the lysate were prepared in LB and stored at room temperature until further use. 20 mL LBA plates supplemented with kanamycin were also prepared using standard sized petri-dishes.

200 μL of 0.2-0.3 OD600 cells, 2 μL 1 M CaCl_2_ and 200 μL lysate dilutions were mixed in a 15 mL culture tube, and incubated stationary at 37 °C for 20 minutes. 4-5 mL of 42-46 °C soft LBA (1:1 mix of LBA with LB) was added to the culture tube and gently mixed. The content of the culture tube was poured onto an LBA plate with 50 µg/mL kanamycin, completely and evenly. The soft top was allowed to solidify for 2-3 minutes before incubating at 37 °C, overnight, in the up-side-down position. Next day, successfully transduced colonies were counted and, when required, further picked and grown overnight in 4 mL LB with 50 µg/mL kanamycin. The respective transduced plasmids were miniprepped next day using the QIAprep Spin Miniprep kit and confirmed by PCR and sequencing.

### 4.6. Transduction assay and kinetics

4 mL of LB were inoculated with *E. coli* EMG2 from a glycerol stock, and incubated overnight at 37 °C, 200 rpm. Next day, the overnight culture was refreshed in 10 mL LB supplemented with 2 mM CaCl_2_ and grown until OD600 = 0.3-0.4, shaking at 37 °C. The protocol continues either on a 96 well plate via reader monitoring growth kinetics or in a culture tube.

For the plate reader experiments, the concentrated stocks were diluted in LB to a titer respective of the multiplicity of infection (MOI) needed when mixing 1:1 with cells at OD600 = 0.2. 100 µL of each diluted lysate were added to a 96-well plate compatible with the Tecan plate reader. The Tecan programme (Supplementary Table S4) was set, and the instrument was heated in advance to 37 °C. At OD600 = 0.3-0.4, the EMG2 culture was diluted to an OD600 ≈ 0.2 in LB (+2 mM CaCl2) just before mixing with the particles. Then, 100 µL of OD600-adjusted culture corresponding to approximately 1.6 × 10⁸ cells/mL were transferred via a 12-channel pipette to the corresponding wells. OD600 measurements were recorded every 5 min.

For the transduction done in a tube instead of the 96 well plate, the culture tubes containing the cells and particles mixture (MOI of 0.1 or 10) were incubated instead for 3 hours, 6 hours and overnight at 37 °C, shaking with aeration.

At corresponding time points, respective transduced cultures (in a plate or from a tube) were transferred to a 1.5 mL Eppendorf tube and pelleted immediately (7 min; 13,000 rpm). The respective supernatants were concentrated individually down to 25 µL via Vivaspin 500 with 10 kDa cut-off, at 4 °C, and according to the Vivaspin manufacturer’s instructions. Each concentrated supernatant and/or the pellets were then mixed with 25 µL Gel Loading Buffer (100 mM Tris-Cl (pH 6.8), 4% (w/v) SDS, 0.2% (w/v) bromophenol blue, 20% (v/v) glycerol, 200 mM dithiothreitol) and boiled for 10 minutes at 95 °C. The end product was finally stored at −20 °C, ready for further analysis by SDS-PAGE and Western blot.

### 4.7. Detection and quantification of model proteins

The BIO-RAD mini-protean TGX precast gel was set up according to the manufacturer’s instructions, using the BIO-RAD Mini-PROTEAN Tetra system and power pack, in 1% TGX running buffer. 5 µL ladder and 25 µL of each denatured sample (concentrated supernatant mixed with loading dye and boiled at 95 °C) were added onto the gel. The instrument was set to electrophorese at 120 V until the dye front reached ∼50-80% down the gel. The proteins on the gel were then carefully transferred onto a PVDF membrane using the BIO-RAD Trans-Blot Turbo Transfer System. The non-specific sites were blocked overnight at 4 °C by soaking in 15 mL 5% albumin fraction V from bovine serum (BSA) prepared in TBS-T (0.05% Tween 20, TBS), where TBS is 20 mM Tris-HCl, 150 mM NaCl, pH 7.5. The membrane was washed twice in 0.1% BSA/TBS-T, followed by incubation (room temperature, gently rocking, for 45 minutes) in 15 mL primary antibody diluted in 0.1% BSA/TBS-T (anti-5xHis-HRP antibody, 1:1000; mouse anti-PE primary antibody, 0.25 µg/mL). The membrane was further washed three times with TBS-T (15 minutes x 1, 5 minutes x 2).

For the membrane treated with anti-PE primary antibody, the membrane was further incubated for another 45 min in 15 mL goat anti-mouse-HRP secondary antibody (0.25 µg/mL; diluted in 0.1% BSA/TBS-T), with slow mixing on a rocker at room temperature.

The membrane was finally washed three times in TBS-T and once in TBS before placing onto a clean flat surface. Amersham ECL reagent was added drop-wise covering the top of the membrane and left for 5 minutes before excess liquid was removed and the interacted bands were investigated for chemiluminescence via the ImageQuant instrument.

For the outer membrane protein E (PE) or PE variants, the band density was also quantified via the ImageJ gel analysis tool taking into account the known value of 300 ng PE ran on the same gel as control.

### 4.8. Indirect transduction assay

The indirect transduction protocol starts with a transducing-forming unit (TFU) assay. The difference is that transduction was done in the *E. coli* Marionette strain, DH10B-Marionette, and not DH5α-Z1; therefore, no spectinomycin was added where mentioned.

### 4.9. Indirect post-transduction cargo expression

4 mL of LB supplemented with 50 µg/mL kanamycin were inoculated with *E. coli* DH10B-Marionette with or without transduced phasmid from a glycerol stock, and incubated overnight at 37 °C, 200 rpm. The next day, the overnight culture was refreshed in 10 mL LB with kanamycin and grown until OD600 = 0.2-0.3, shaking at 37 °C. Then, 200 µL of each culture was transferred 12 times to a 96-well plate compatible with the Tecan instrument, preheated to 37 °C, including 12 wells containing 200 µL of LB only. The Tecan programme (Supplementary Table S4) was set, and the cells were allowed to grow until OD600 = 0.4, when the plate reader was paused to add inducer with a multichannel pipette, where relevant. Then, 10 µL of inducer stock solution was added to bring the culture to a final inducer concentration of 100 µM vanillic acid (Van) and, separately, 200 nM anhydrotetracycline (aTc). Where no inducer or one inducer only was added, the cultures were supplemented with 20 µL or 10 µL LB, respectively.

When both aTc and Van were supplemented, the plate reader was paused at half-time lysis to retrieve sample for protein expression analysis in the pellet and supernatant. When only Van was added, the plate reader was paused after 3 hours and next day to retrieve sample for protein expression analysis in the pellet only. The protein analysis was performed as detailed in Section 4.7, using anti-PE primary antibody and anti-mouse-HRP secondary antibody.

### 4.10. Preparation of electrocompetent cells for efficient phasmid transformation

One colony of *E. coli* C-5545Δcos-Marionette production strain was inoculated into 4 mL LB supplemented with 15 µg/mL trimethoprim. The culture was grown overnight at 37 °C, shaking 200 rpm. The overnight culture was refreshed (1:100 ratio) into 100 mL LB with 15 µg/mL trimethoprim. The culture was then incubated at 37 °C, shaking 200 rpm until OD600 = 0.5-0.8. To stop the growth, the culture was placed on ice for 20-30 minutes. The cells were centrifuged for 10 minutes at 2,254 rcf, the supernatant was discarded and the pellets were gently resuspended in 100 mL 10% cold glycerol solution. The resuspended pellets were kept on ice for another 20-30 minutes. The centrifugation, resuspension and ice cycle were repeated 3 more times, with less volume each time: 25 mL, 10 mL 10% cold glycerol solution. The final pellets were resuspended in 2 mL of 10% cold glycerol solution, and 100 µL aliquots were transferred to ice-cold Eppendorf tubes on ice and stored at −80 °C. These electro-competent cells were used to produce high titer helper-free phage-like particles, as detailed below.

### 4.11. Concentrating particles

The particles were concentrated by polyethylene glycol (PEG) precipitation or by Vivaspin Ultrafiltration at 100 kDa and 300 kDa cut-offs.

The PEG precipitation protocol was adjusted from M. Tridgett PhD thesis (2019) [54]. The PEG solution (2.5 M NaCl, 25% PEG 6000) was added to the filtered lysate, in a 1:5 volumetric ratio. The lysate with PEG solution was mixed for 20-24 hours at 4 °C, using a magnetic stirrer. The mixture was centrifuged next day for 15-16 hours at 3,428 rcf, 4 °C. The supernatant was discarded and the pellet was resuspended in a total of 5-7 mL P4 buffer (250 mM NaCl, 80 mM MgCl_2_, 0.08% EGTA). To remove the remaining cell debris, the resuspension was centrifuged for 5 minutes at 17,949 rcf, 4 °C. The supernatant was collected and the pellet was resuspended in 5-7 mL P4 buffer and centrifuged once more to dislocate any remaining phage-like particles stuck onto the cell debris. The first and second supernatants were mixed with PEG solution in a 1:5 volumetric ratio, and left to spin overnight on a rocker, at 4 °C. The mixture was centrifuged next day for 15 minutes at 17,949 rcf, 4 °C. The pellet containing phage-like particles was resuspended in 1 mL buffer of choice (*e.g.* for endotoxin removal, stability test).

The concentration by ultrafiltration took typically 6-8 hours using two Vivaspin 20 ultrafiltrators in parallel, spinning with 3,000 rcf, at 4 °C. The run-through lysate was discarded each time and the concentrator body was refilled with remaining lysate due for concentrating as needed. New concentrator was used if/when the filtrate got clogged.

### 4.12. Endotoxin removal and quantification

Endotoxin removal was performed with endotoxin-free pipette tips, and the respective buffers were prepared with endotoxin-free water or endotoxin-free PBS. Endotoxin removal was attempted by different variations of buffer exchange with or without Triton X-100 detergent via PEG precipitation and isoelectric-point (pI) precipitation, by 1-octanol extraction, and separately by affinity chromatography with the commercially available ToxinEraser kit.

Endotoxin removal by PEG buffer exchange was performed by mixing 100 µL phage-like particle concentrate (prepared by ultrafiltration with either 100 or 300 kDa cut-off; no statistical difference was found between the two concentration methods in endotoxin accumulation; Supplementary Figure S1.3) with 200 µL PEG solution (2.5 M NaCl, 25% PEG 6000) and with or without Triton X-100 at varying concentrations (2%, 4%, 6%). The sample was made up to 1 mL with Tris buffer (20 mM Tris pH 7.5, 100 mM NaCl). The sample was then mixed on a rocker for 1.5 h, 3 h or overnight at 4 °C. The phage-like particles were sedimented by centrifuging the sample for 10 min at 15,000 g, 22 °C. Particles were then resuspended in the respective PEG/Triton X-100 mix and filled up to 1 mL with Tris buffer to continue with the precipitation rounds, or the particles were resuspended directly in 1 mL Tris buffer. 50 µL of resuspended sample was collected between runs to examine endotoxin amount and particle recovery.

Endotoxin removal by ip buffer exchange was performed by first precipitating the phage-like particles with PEG as above, followed by resuspension in Tris Buffer at pH 4.7. The ip precipitation method was also used at the end of the buffer exchange by PEG with Triton X-100 to remove downstream contaminants.

Endotoxin removal by 1-octanol extraction was performed as documented in Bonilla *et al.* 2016 [38]. Endotoxin removal using the commercially available ToxinEraser kit was performed in a cold room at 4 °C. The final pellet after concentration of phage-like particles via PEG precipitation was resuspended in P4 TE buffer: 20 mM Tris-HCl pH 7.5, 250 mM NaCl. The buffer was prepared from 1 M Tris-HCl pH 7.5 buffer and endotoxin-free PBS supplemented with NaCl powder. The column was treated according to the manufacturer’s instructions, except for the additional equilibration step with P4 TE buffer, which was added before applying the sample containing phage-like particles onto the column. A 1 mL aliquot of the resuspended phage-like particles was added to the column and collected after a 1.5 mL dead volume. An extra 1 mL P4 TE buffer was added to the column to allow the flow-through of a total of 1 mL endotoxin-reduced phage-like particles. The column was then equilibrated with the ToxinEraser equilibration buffer and treated according to the manufacturer’s instructions for reloading the sample up to a total of five times or for storage. A 20 µL aliquot of endotoxin-reduced phage-like particles was collected between ToxinEraser run-throughs for quantification of particles and endotoxins.

Particle concentration was examined by a plaque-forming unit assay. The amount of endotoxin present in the sample was quantified with the Pierce LAL Chromogenic Endotoxin Quantification kit, according to the manufacturer’s instructions. Low amounts of Triton X-100 were found to mask the presence of endotoxins in the sample (Supplementary Figure S1.2). LAL interference was therefore actively monitored by measuring the unknown samples treated with Triton side by side with the respective samples spiked with 0.5 EU/mL. All plotted values were within the two-fold variation limit of the LAL assay.

### 4.13. Isoelectric precipitation and short-term stability

Phage-like particles concentrated by ultrafiltration with 300 kDa cut-off were diluted 10,000 times in buffer with varying pH values: pH 3, 4, 4.2, 4.4, 4.7, 4.9, 5, 6, 7 and 8, to a final volume of 1 mL (Supplementary Table S2). The pH was measured using the Mettler Toledo Seven2Go S2 pH meter. The diluted phage-like particles in respective pH buffer were mixed on a rocker at room temperature for 15 minutes. The particles in respective pH buffer were then centrifuged for 15 minutes at 13,000 rcf, at 4 °C. Each supernatant was transferred to a new tube and each pellet was resuspended in 1 mL pH 7 buffer supplemented with 80 mM MgCl_2_. The phage-like particles in both the resuspended pellet and supernatant were quantified by plaque-forming unit assay to monitor for isoelectric precipitation and short-term stability.

### 4.14. Hydrophobicity

The helper-free phage-like particles were grown at small scale, as described in Section 2. The method for testing hydrophobicity was adapted from Bonilla *et al.* (2016) [38], a promising method for purifying phage stocks based on endotoxin removal by extraction with organic solvents, inspired by Szermer-Olearnik *et al.* (2015) [40].

A 6 mL lysate containing helper-free phage-like particles was mixed with 4 mL concentrated 1-octanol (or 1-butanol) in a 15 mL Falcon tube. The sample was shaken on a rocker for 1.5 h at room temperature. The Falcon tube was placed on ice and the mixture was allowed to settle for 15 min, stationary. The mixture was then further partitioned into layers by centrifugation at 3,428 rcf for 15 min. The bottom layer, corresponding to the aqueous phase, was carefully extracted using a syringe by gently piercing through the bottom of the tube with the syringe needle, and transferred to a new 15 mL Falcon tube. The surviving particles present in the aqueous phase were quantified by plaque-forming unit assay and compared with the total number of particles before solvent treatment.

### 4.15. Long-term stability

Concentrated particles were resuspended in three buffers in parallel: Buffer 1 (1x PBS, 20 mM MgSO_4_, pH 7), Buffer 2 (20 mM Tris, 127 mM NaCl, 2 mM CaCl_2_, 5 mM MgSO_4_, pH 7) and Buffer 3 (20 mM Tris, 132 mM NaCl, 2 mM CaCl_2_, pH 7). The particles were then stored in triplicate in the fridge (2-8 °C) and at room temperature (22 ± 2 °C). The stability of the particles was monitored by a plaque-forming unit assay, as follows: the particles stored in the fridge were quantified every 2 weeks for 2 months and the particles stored at room temperature were quantified once after 24 hours.

## Data Availability

All data generated or analysed during this study are included in this article and its Supplementary Information files. Sequence information for engineered constructs is provided in Supplementary Tables S9-S22.

## Acknowledgements

We thank Frenk Smrekar and Kaja Malalan from Jafral for insightful phage bioprocessing discussions. We thank Dr Marta Vila Rico from GSK for helpful discussions on antigen/adjuvant design and testing. We thank Stéphanie Deroo, Steven Sijmons and Timothée Laloux from GSK for helpful manuscript comments. We thank Robert Ramirez-Garcia for early input related to this work. We also thank WISB for use of their facilities.

## Author Contributions

SG and AJ conceived the study and supervised the project. SG, AJ, MA, MT, CC and NB designed the methodology. MA performed the experiments and analysed the data with input from SG, AJ and NB. MT and CC performed preliminary experiments presented in the Supplementary Information. MA, AJ and SG provided resources. MA wrote the original draft. SG, CC and AJ edited the manuscript. All authors approved the final manuscript.

## References

[1] Hitchcock, N. M. et al. Current clinical landscape and global potential of bacteriophage therapy. Viruses 15, 1020; 10.3390/v15041020 (2023).

[2] Pirnay, J.-P. et al. Personalized bacteriophage therapy outcomes for 100 consecutive cases: a multicentre, multinational, retrospective observational study. Nat. Microbiol. 9, 1434–1453 (2024).

[3] Rohde, C. et al. Expert opinion on three phage therapy related topics: bacterial phage resistance, phage training and prophages in bacterial production strains. Viruses 10, 178; 10.3390/v10040178 (2018).

[4] Brives, C. & Pourraz, J. Phage therapy as a potential solution in the fight against AMR: obstacles and possible futures. Palgrave Commun. 6, 100; 10.1057/s41599-020-0478-4 (2020).

[5] Principi, N., Silvestri, E. & Esposito, S. Advantages and limitations of bacteriophages for the treatment of bacterial infections. Front. Pharmacol. 10, 513; 10.3389/fphar.2019.00513 (2019).

[6] Dedrick, R. M. et al. Engineered bacteriophages for treatment of a patient with a disseminated drug-resistant Mycobacterium abscessus. Nat. Med. 25, 730–733 (2019).

[7] Hatfull, G. F., Dedrick, R. M. & Schooley, R. T. Phage therapy for antibiotic-resistant bacterial infections. Annu. Rev. Med. 73, 197–211 (2022).

[8] Pires, D. P., Cleto, S., Sillankorva, S., Azeredo, J. & Lu, T. K. Genetically engineered phages: a review of advances over the last decade. Microbiol. Mol. Biol. Rev. 80, 523–543 (2016).

[9] Gibb, B., Hyman, P. & Schneider, C. L. The many applications of engineered bacteriophages-an overview. Pharmaceuticals 14, 634; 10.3390/ph14070634 (2021).

[10] Elois, M. A., da Silva, R., Pilati, G. V. T., Rodríguez-Lázaro, D. & Fongaro, G. Bacteriophages as biotechnological tools. Viruses 15, 349; 10.3390/v15020349 (2023).

[11] Tridgett, M., Ababi, M., Osgerby, A., Ramirez-Garcia, R. & Jaramillo, A. Engineering bacteria to produce pure phage-like particles for gene delivery. ACS Synth. Biol. 10, 107–114 (2021).

[12] Westwater, C., Schofield, D. A., Schmidt, M. G., Norris, J. S. & Dolan, J. W. Development of a P1 phagemid system for the delivery of DNA into Gram-negative bacteria. Microbiology 148, 943–950 (2002).

[13] Cronan, J. E. Two neglected but valuable genetic tools for Escherichia coli and other bacteria: in vivo cosmid packaging and inducible plasmid replication. Mol. Microbiol. 120, 783–790 (2023).

[14] Rao, V. B. & Zhu, J. Bacteriophage T4 as a nanovehicle for delivery of genes and therapeutics into human cells. Curr. Opin. Virol. 55, 101255 (2022).

[15] Meyer, A. J., Segall-Shapiro, T. H., Glassey, E., Zhang, J. & Voigt, C. A. Escherichia coli “Marionette” strains with 12 highly optimized small-molecule sensors. Nat. Chem. Biol. 15, 196–204 (2019).

[16] Six, E. W. & Klug, C. A. C. Bacteriophage P4: a satellite virus depending on a helper such as prophage P2. Virology 51, 327–344 (1973).

[17] Geisselsoder, J. et al. Mutants of satellite virus P4 that cannot derepress their bacteriophage P2 helper. J. Mol. Biol. 148, 1–19 (1981).

[18] Kahn, M., Ow, D., Sauer, B., Rabinowitz, A. & Calendar, R. Genetic analysis of bacteriophage P4 using P4-plasmid ColE1 hybrids. Mol. Gen. Genet. 177, 399–412 (1980).

[19] Souza, L., Calendar, R., Six, E. W. & Lindqvist, B. H. A transactivation mutant of satellite phage P4. Virology 81, 81–90 (1977).

[20] Sauer, B., Ow, D., Ling, L. & Calendar, R. Mutants of satellite bacteriophage P4 that are defective in the suppression of transcriptional polarity. J. Mol. Biol. 145, 29–46 (1981).

[21] Geisselsoder, J., Chidambaram, M. & Goldstein, R. Transcriptional control of capsid size in the P2:P4 bacteriophage system. J. Mol. Biol. 126, 447–456 (1978).

22. Shore, D., Dehò, G., Tsipis, J. & Goldstein, R. Determination of capsid size by satellite bacteriophage P4. Proc. Natl. Acad. Sci. USA 75, 400-404 (1978).

[23] Gross, M. K., Au, D. C., Smith, A. L. & Storm, D. R. Targeted mutations that ablate either the adenylate cyclase or hemolysin function of the bifunctional cyaA toxin of Bordetella pertussis abolish virulence. Proc. Natl. Acad. Sci. USA 89, 4898–4902 (1992).

[24] Barbieri, J. T. & Cortina, G. ADP-ribosyltransferase mutations in the catalytic S-1 subunit of pertussis toxin. Infect. Immun. 56, 1934–1941 (1988).

[25] Guan, Y. et al. An equation to estimate the difference between theoretically predicted and SDS PAGE-displayed molecular weights for an acidic peptide. Sci. Rep. 5, 13370; 10.1038/srep13370 (2015).

[26] Ramirez-Garcia, R., Sagona, A. P., Barr, J. J. & Jaramillo, A. High-yield bioproduction of virus-free virus-like P4-EKORhE multi-lysin transducing particles as an antimicrobial gene therapeutic. Front. Cell. Infect. Microbiol. 15, 1561443; 10.3389/fcimb.2025.1561443 (2025).

[27] Ronander, E. et al. Identification of a novel *Haemophilus influenzae* protein important for adhesion to epithelial cells. Microbes Infect. 10, 87–96 (2008).

[28] Ronander, E. et al. Nontypeable *Haemophilus influenzae* adhesin protein E: characterization and biological activity. J. Infect. Dis. 199, 522–531 (2009).

[29] Hallström, T., Blom, A. M., Zipfel, P. F. & Riesbeck, K. Nontypeable *Haemophilus influenzae* protein E binds vitronectin and is important for serum resistance. J. Immunol. 183, 2593–2601 (2009).

30. Shore, D., Dehò, G., Tsipis, J. & Goldstein, R. Determination of capsid size by satellite bacteriophage P4. Proc. Natl. Acad. Sci. USA 75, 400-404 (1978).

[31] Kim, K.-J., Sunshine, M. G., Lindqvist, B. H. & Six, E. W. Capsid size determination in the P2-P4 bacteriophage system: suppression of sir mutations in P2’s capsid gene N by Supersid mutations in P4’s external scaffold gene *sid*. Virology 283, 49–58 (2001).

[32] Okazaki, N., Matsuo, S., Saito, K., Tominaga, A. & Enomoto, M. Conversion of the Salmonella phase 1 flagellin gene fliC to the phase 2 gene fljB on the Escherichia coli K-12 chromosome. J. Bacteriol. 175, 758–766 (1993).

[33] Didierlaurent, A. et al. Flagellin promotes myeloid differentiation factor 88-dependent development of Th2-type response. J. Immunol. 172, 6922–6930 (2004).

[34] McSorley, S. J., Ehst, B. D., Yu, Y. & Gewirtz, A. T. Bacterial flagellin is an effective adjuvant for CD4+ T cells *in vivo*. J. Immunol. 169, 3914–3919 (2002).

[35] Eom, J. S. et al. Enhancement of host immune responses by oral vaccination to Salmonella enterica serovar Typhimurium harboring both FliC and FljB flagella. PLoS ONE 8, e74850; 10.1371/journal.pone.0074850 (2013).

[36] Summers, D. K. & Sherratt, D. J. Multimerization of high copy number plasmids causes instability: ColE1 encodes a determinant essential for plasmid monomerization and stability. Cell 36, 1097–1103 (1984).

[37] Luong, T., Salabarria, A.-C., Edwards, R. A. & Roach, D. R. Standardized bacteriophage purification for personalized phage therapy. Nat. Protoc. 15, 2867–2890 (2020).

[38] Bonilla, N. et al. Phage on tap-a quick and efficient protocol for the preparation of bacteriophage laboratory stocks. PeerJ 4, e2261; 10.7717/peerj.2261 (2016).

[39] Ma, R. et al. Endotoxin removal during the purification process of human-like collagen. Sep. Sci. Technol. 45, 2400–2405 (2010).

[40] Szermer-Olearnik, B. & Boratyński, J. Removal of endotoxins from bacteriophage preparations by extraction with organic solvents. PLoS ONE 10, e0122672; 10.1371/journal.pone.0122672 (2015).

[41] Yamashita, M., Murahashi, H., Tomita, T. & Hirata, A. Effect of alcohols on Escherichia coli phages. Biocontrol Sci. 5, 9–16 (2000).

[42] Branston, S. D., Wright, J. & Keshavarz-Moore, E. A non-chromatographic method for the removal of endotoxins from bacteriophages. Biotechnol. Bioeng. 112, 1714–1719 (2015).

[43] Tao, H. et al. Purifying natively folded proteins from inclusion bodies using sarkosyl, Triton X-100 and CHAPS. BioTechniques 48, 61–64 (2010).

[44] Dong, D., Sutaria, S., Hwangbo, J. Y. & Chen, P. A simple and rapid method to isolate purer M13 phage by isoelectric precipitation. Appl. Microbiol. Biotechnol. 97, 8023–8029 (2013).

[45] U.S. Food and Drug Administration. Setting endotoxin limits during development of investigational oncology drugs and biological products: draft guidance for industry. https://www.fda.gov/regulatory-information/search-fda-guidance-documents/setting-endotoxin-limits-during-development-investigational-oncology-drugs-and-biological-products (2020).

[46] Scholl, D. & Martin, D. W. Antibacterial efficacy of R-type pyocins towards Pseudomonas aeruginosa in a murine peritonitis model. Antimicrob. Agents Chemother. 52, 1647–1652 (2008).

[47] Brödel, A. K. et al. In situ targeted base editing of bacteria in the mouse gut. Nature 632, 877–884 (2024).

[48] Gencay, Y. E. et al. Engineered phage with antibacterial CRISPR-Cas selectively reduce *E. coli* burden in mice. Nat. Biotechnol. 42, 265–274 (2024).

[49] Baker, Z. R. et al. Sustained in situ protein production and release in the mammalian gut by an engineered bacteriophage. Nat. Biotechnol. 44, 90–99 (2026).

[50] Fa-Arun, J., Huan, Y. W., Darmon, E. & Wang, B. Tail-engineered phage P2 enables delivery of antimicrobials into multiple gut pathogens. ACS Synth. Biol. 12, 596–607 (2023).

[51] Kahn, M. L. et al. Bacteriophage P2 and P4. Methods Enzymol. 204, 264–280 (1991).

[52] Ow, D. W. & Ausubel, F. M. Recombinant P4 bacteriophages propagate as viable lytic phages or as autonomous plasmids in Klebsiella pneumoniae. Mol. Gen. Genet. 180, 165–175 (1980).

[53] Yosef, I., Goren, M. G., Globus, R., Molshanski-Mor, S. & Qimron, U. Extending the host range of bacteriophage particles for DNA transduction. Mol. Cell 66, 721–728.e3 (2017).

[54] Tridgett, M. Development of nanomaterials based on bacteriophage-chromophore bioconjugates. PhD thesis, University of Birmingham (2019).

