## Supplementary Information for "Engineering P4 satellite phage particles with single and dual antimicrobial actions"

[†] Joint last authors.

### Supplementary Figures
#### S1. Bioprocessing

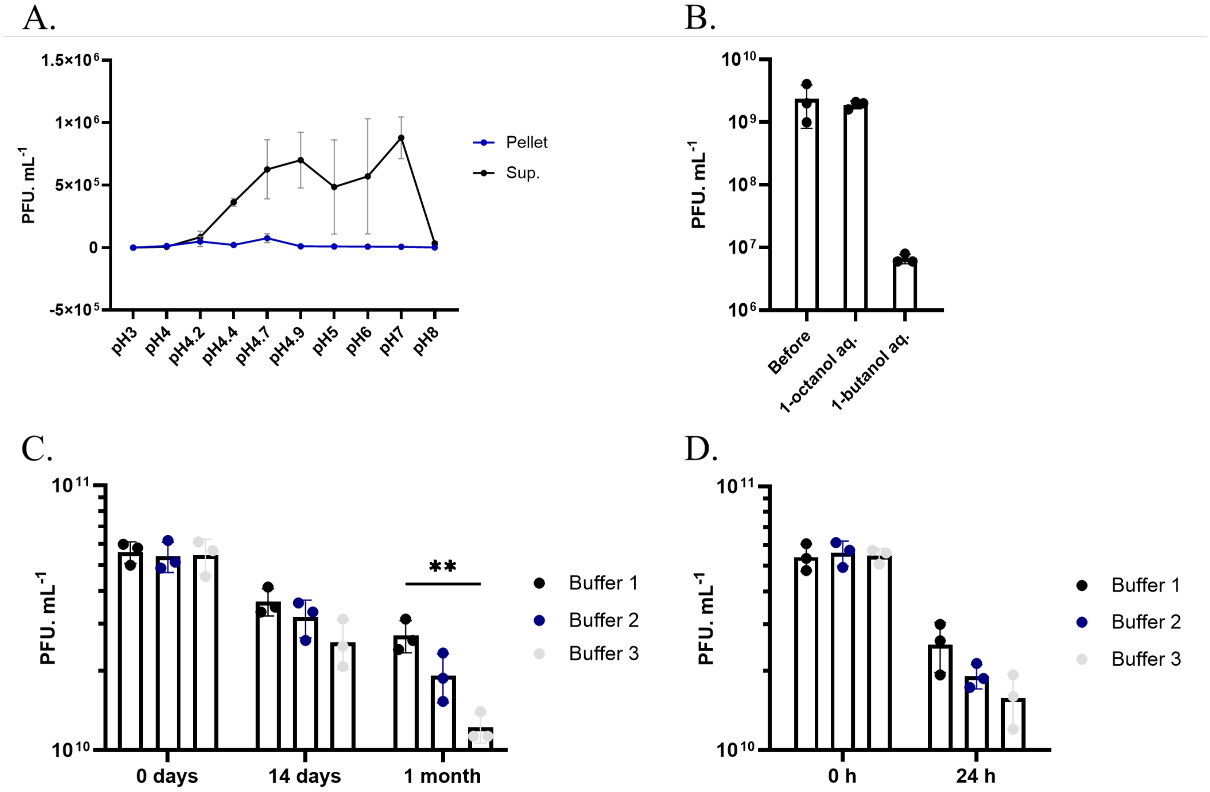


**Supplementary Figure S1.1 | P4 characteristics. A,** Isoelectric point precipitation and short-term stability, using P4-DiffJJ-Test particles. Pellet = phage-like particles accumulation based on isoelectric point precipitation during centrifugation at 13,000 x g; resuspended in 1 mL Tris Buffer (20 mM Tris, 50 mM NaCl, pH 7). Sup. = stable phage-like particles remaining after isoelectric point precipitation; in original buffer with varying pH values. **B,** Hydrophobicity, using P4-DiffJJ-Test particles. 1-octanol/butanol aq. = the number of particles present in the aqueous phase after phase partitioning using 40% 1-octanol or 1-butanol, respectively. C-**D,** Long-term stability (one month; 2-8 °C) and accelerated stability (24 h; 22 ± 2 °C), respectively; using P4-pLtetO1-LacIq particles; Buffer 1 = 1x PBS, 20 mM $MgSO_4$, pH 7; Buffer 2 = 20 mM Tris, 127 mM NaCl, 2 mM $CaCl_2$, 5 mM $MgSO_4$, pH 7; Buffer 3 = 20 mM Tris, 132 mM NaCl, 2 mM $CaCl_2$, pH 7. **$P \leq 0.01$; (n = 3).

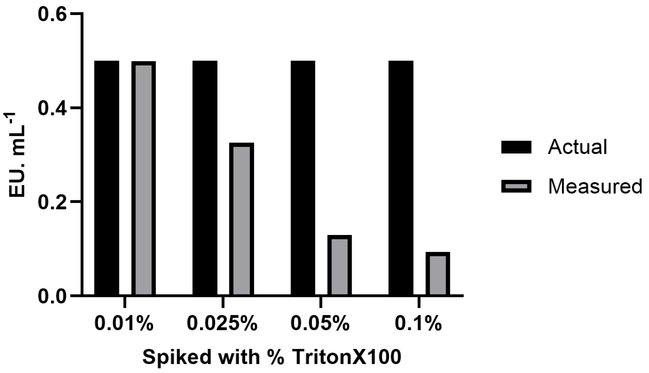


**Supplementary Figure S1.2 | Triton X-100 interference with the LAL endotoxin assay.** P4 particle samples were tested to determine whether Triton X-100 affected limulus amebocyte lysate endotoxin quantification. n = 1.

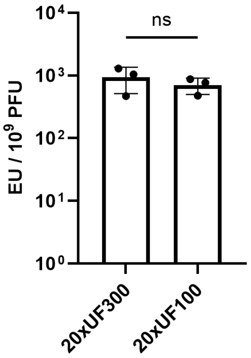


**Supplementary Figure S1.3 | Endotoxin accumulation after particle concentration by ultrafiltration.** P4 particle stocks were concentrated with 100 kDa or 300 kDa molecular-weight cut-off ultrafiltration membranes before endotoxin quantification. Pns > 0.05. n = 3.

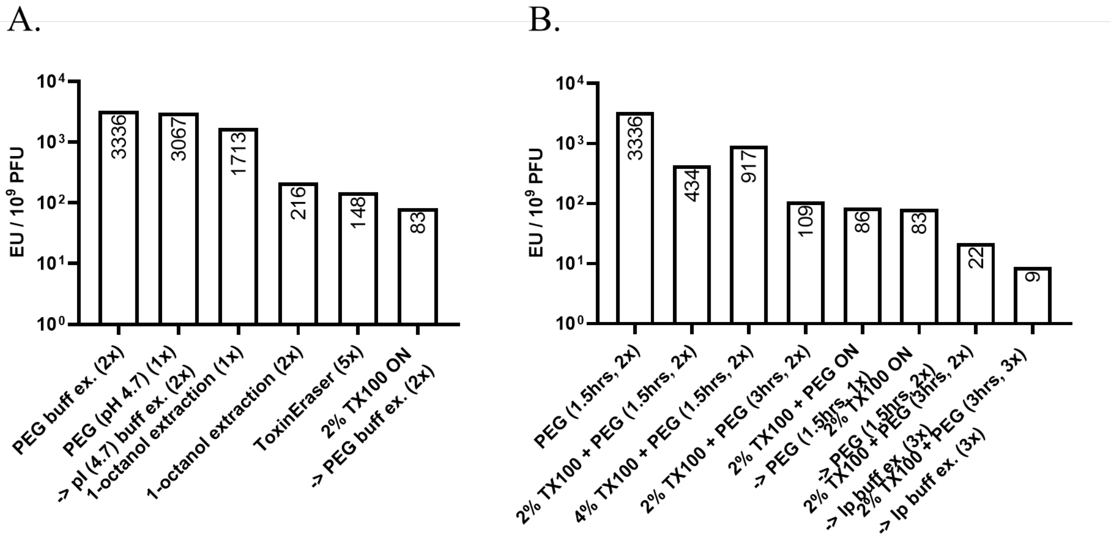


**Supplementary Figure S1.4 | Endotoxin removal from P4 cargo stocks. A,** Methods: Buffer exchange by PEG and isoelectric point (pH 4.7) precipitation; two-phase separation by 1-octanol extraction; polymyxin B affinity chromatography via the ToxinEraser kit; treatment with Triton X-100 overnight before two rounds of PEG6000/NaCl buffer exchange. **B,** PEG precipitation varying Triton X-100 concentration (0%, 2%, 4%), precipitation order (Triton X-100 mixed with PEG; Triton X-100 alone overnight, followed by PEG precipitation), incubation times (1.5 h, 3 h, overnight) and total precipitation rounds (2x, 3x, 6x). TX100 = Triton X-100; ON = overnight; PEG = PEG6000/NaCl precipitation; (n = 1).

#### S2. Phage-based single-action prototype

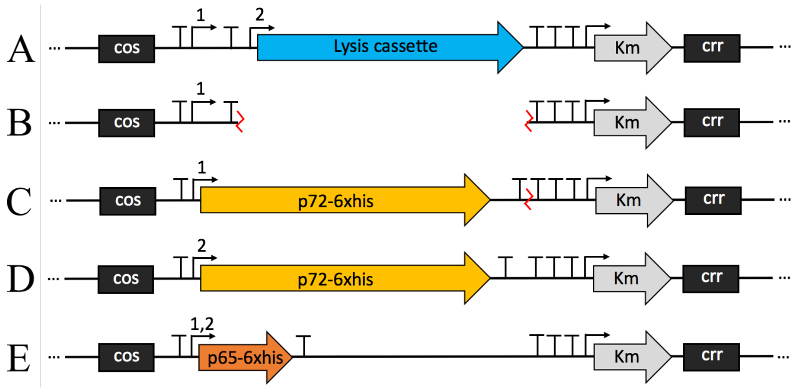


**Supplementary Figure S2.1 | P4 phasmid designs containing cargo. A,** The vector backbone, pP4-pLtetO1-LacIq (11,627 bp), used for cloning and Genewiz synthesis. **B,** Intermediate construct, pP4-Δlys, used to build pP4-LacIq-p72 (constitutive). **C,** pP4-LacIq-p72 (11,626 bp), built by Gibson Assembly of opened pP4-Δlys vector and amplified p72 fragment. **D,** pP4-pLtetO1-p72 (repressed) (11,624 bp), built by Genewiz synthesis; **E,** pP4-LacIq-p65 (constitutive) and pP4-pLtetO1-p65 (repressed) (11,624 bp), built by Genewiz synthesis. 1 = $P_{lacI}^{q}$ promoter; 2 = $P_{LtetO\text{-}1}$ promoter.

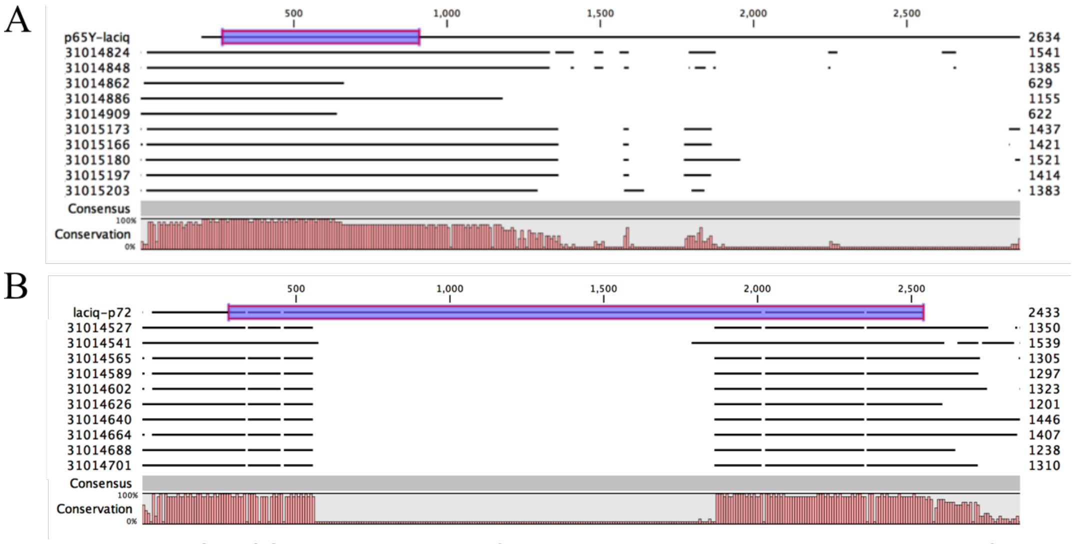


**Supplementary Figure S2.2 | Sequencing results for $P_{lacI}^{q}$ regulated constructs minipreppped from transduced EMG2, mixed with P4_col1_F primer. A,** pP4-LacIq-p65; **B,** pP4-LacIq-p72. Original sequence is on first row, with *p65*/*p72* gene highlighted.

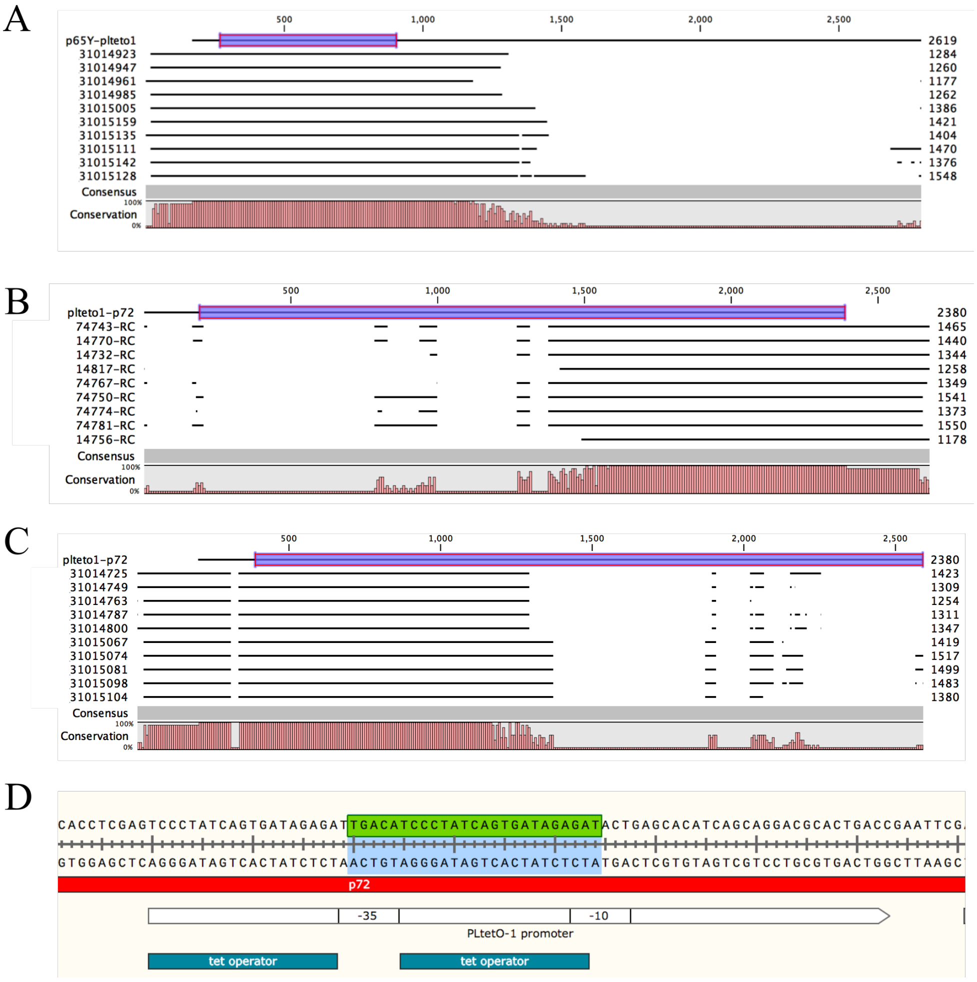


**Supplementary Figure S2.3 | Sequencing results for PLtetO-1 regulated constructs minipreppped from transduced EMG2. A,** pP4-pLtetO1-p65, mixed with P4-col1-F primer; **B,** pP4-pLtetO1-p72, mixed with pP4_Del2_Rev primer; **C,** pP4-pLtetO1-p72, mixed with P4-col1-F primer; A-**C,** Original sequence is on first row, with p65/p72 gene highlighted. **D,** The fragment deletion from panel C is highlighted on the PLtetO-1 promoter sequence.

#### S3. Phage-based dual-action prototype

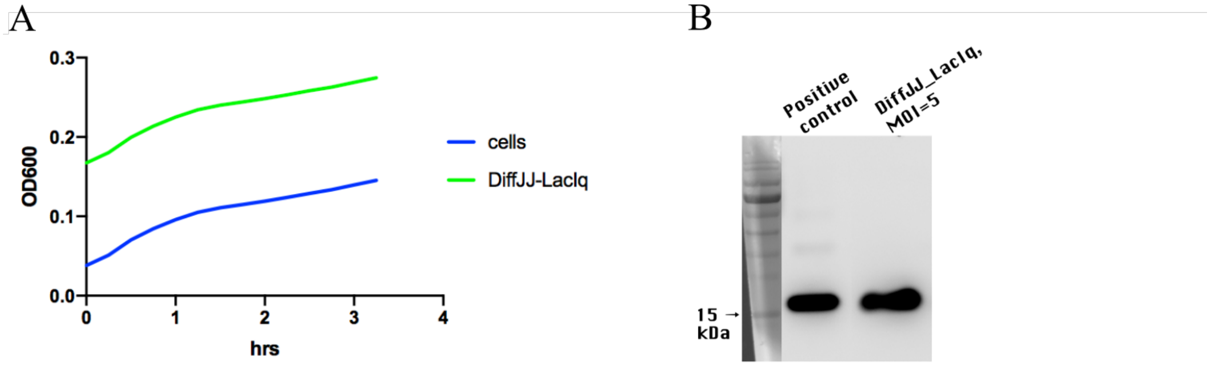


**Supplementary Figure S3.1 | Post-transduction differential expression with P4-DiffJJ-LacIq particles.** The particles carry a lytic cassette regulated by PDiffJJ and PE regulated by PlacIq. *E. coli* EMG2 was used as the target strain. **A,** Lysis kinetics after transduction into *E. coli* EMG2. Cells = EMG2-only control. PDiffJJ-PlacIq = EMG2 mixed with P4-DiffJJ-LacIq particles. **B,** Post-transduction PE detection in the pellet fraction. Positive control = 300 ng PE. PDiffJJ_PlacIq, MOI = 5 = pellet from the panel A transduction condition.

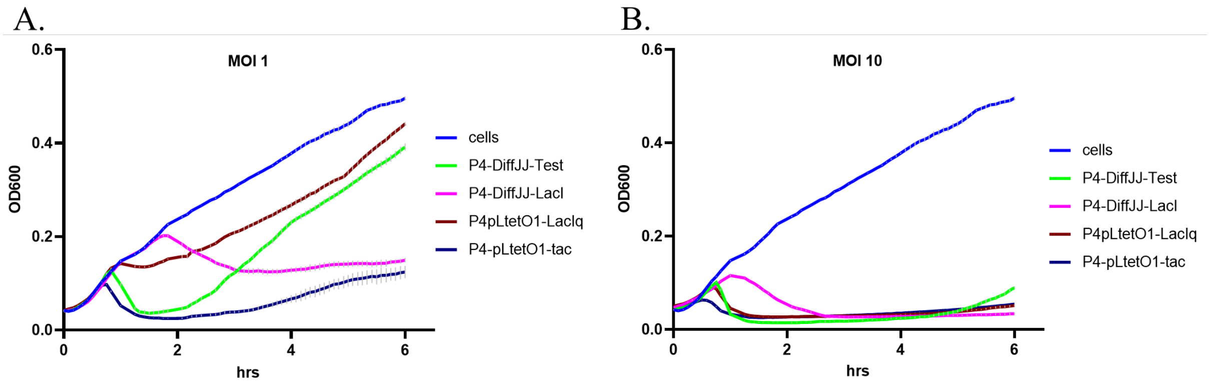


**Supplementary Figure S3.2 | EMG2 post-transduction lysis kinetics when mixed with P4-like particles. A,** at MOI = 1; **B,** at MOI = 10. The starting cell density was OD600 = 0.2. The data in the graph is normalized against the OD600 of the dilution media used for bringing the particles to set MOI: LB + 80 mM $MgCl_2$. P4-DiffJJ-Test (lysis-only) = Lysis kinetics of EMG2 mixed with P4-like particles encapsulating a lytic cassette regulated by the JJ weaker variant of the $P_{LtetO-1}$ promoter; P4-DiffJJ-LacI (dual weak/PE), P4-pLtetO1-LacIq (dual PLtetO-1/PE) and P4-pLtetO1-tac (dual PLtetO-1/PE) = Lysis kinetics of EMG2 mixed with P4-like particles encapsulating a lytic cassette and the PE antigen regulated by the JJ weaker variant of the $P_{LtetO-1}$ promoter ($P_{DiffJJ}$) or $P_{LtetO-1}$ and by the $P_{lacI}$, $P_{lacI}^{q}$ or $P_{tac}$ promoters, respectively.

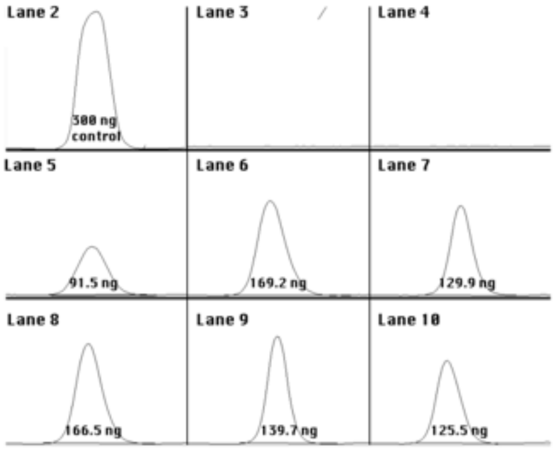


**Supplementary Figure S3.3 | Semi-quantification of the Figure 10B PE Western blot.** ImageJ/Fiji area-under-the-peak analysis was used to estimate PE band density relative to the 300 ng PE control in lane 2. The panel shows antigen detected in *E. coli* EMG2 culture supernatants at perceived half-time and full-time lysis after transduction with the indicated particles. Lane 1 = ladder, not shown in the quantification panel. Lane 2 = 300 ng PE control. Lanes 3-4 = P4-DiffJJ-Test (MOI = 2). Lanes 5-6 = P4-DiffJJ-LacI (MOI = 4). Lanes 7-8 = P4-pLtetO1-LacIq (MOI = 4). Lanes 9-10 = P4-pLtetO1-tac (MOI = 2).

#### S4. Sid-deletion large-cargo dual-action prototype

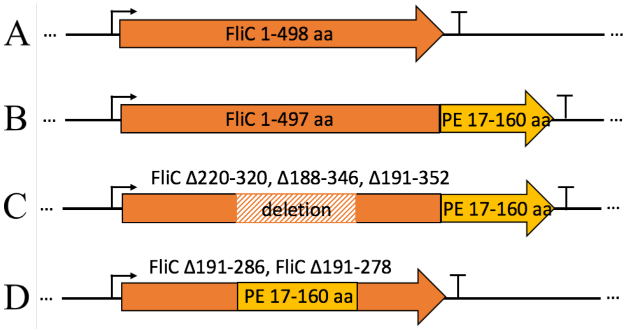


**Supplementary Figure S4.1 | FliC-PE and its variants in pET-24b vector. A,** FliC; **B,** FliC-PE; **C,** FliC-PE deletion variants: FliCΔ220-320-PE, FliCΔ188-346-PE, FliCΔ191-352-PE; **D,** FliCΔ191-286-PE, FliCΔ191-278-PE. Regulated by the T7 promoter.

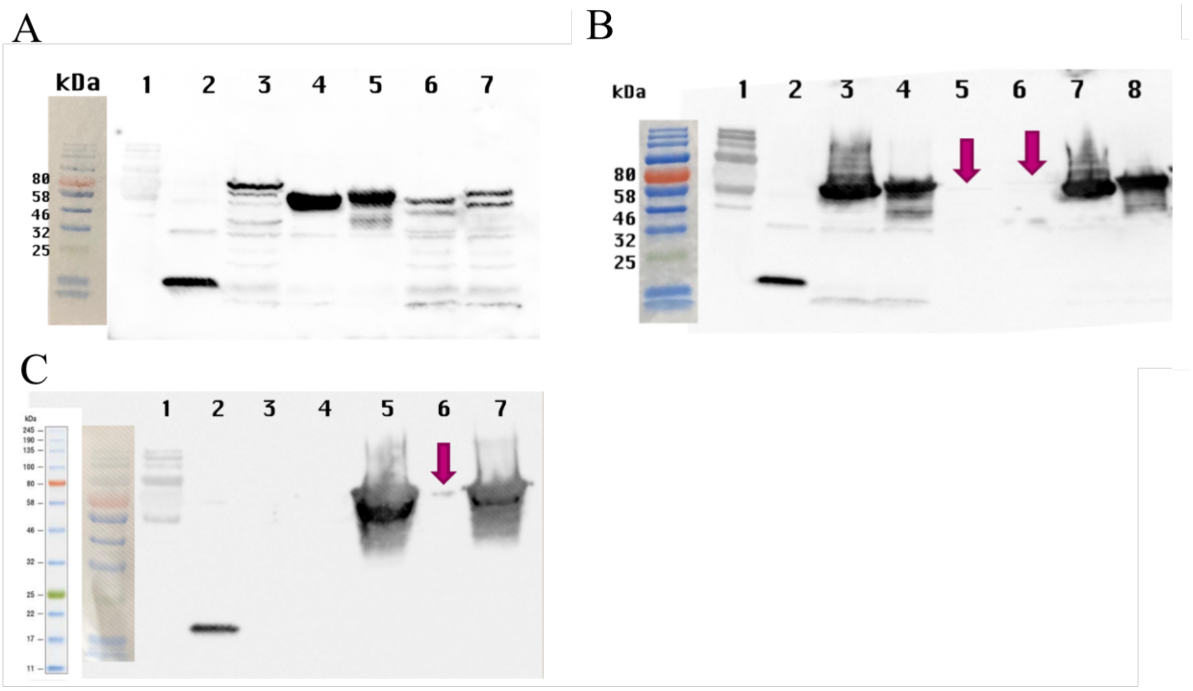


**Supplementary Figure S4.2 | Expression of FliC-PE and variants in BL21-DE3. A,** Pellets of induced BL21-DE3 cells containing pET-24b coding for FliC-PE and its variants; Lane 1 = Color Prestained Protein Standard, Broad Range (11-245 kDa); Lane 2 = 300 ng PE control; Lane 3 = FliC-PE; Lane 4 = FliCΔ188-346-PE; Lane 5 = FliCΔ191-286-PE; Lane 6 = FliCΔ191-352-PE; Lane 7 = FliCΔ220-320-PE. **B,** Total, soluble and insoluble fractions of FliCΔ188-346-PE and FliCΔ191-286-PE, respectively. Lane 1 = Color Prestained Protein Standard, Broad Range (11-245 kDa); Lane 2 = 300 ng PE control; Lanes 3, 4 = total fraction; Lanes 5, 6 = soluble fraction; Lanes 7, 8 = insoluble fraction. **C,** Total, soluble and insoluble fractions of FliCΔ191-278-PE; Lane 1 = Color Prestained Protein Standard, Broad Range (11-245 kDa); Lane 2 = 300 ng PE control; Lanes 3, 4 = BL21-DE3-only soluble and insoluble fractions, respectively; Lanes 5, 6, 7 = BL21-DE3 with FliCΔ191-278-PE total, soluble and insoluble fractions, respectively.

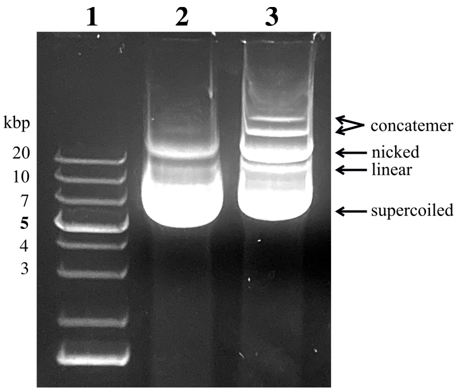


**Supplementary Figure S4.3 | Agarose-gel pattern consistent with enlarged *sid*-deleted P4 phasmids.** Lane 1 = Thermo Scientific GeneRuler 1 kb Plus DNA Ladder; Lane 2 = transduced *sid*-intact P4 phasmid with cargo (pP4-pLtetO1-LacIq (standard-size)); Lane 3 = transduced P4 phasmid with *sid* knockout and large cargo (pP4Δ*sid*-pLtetO1-LacIq (Δ*sid* constitutive)). This assay supports transfer of enlarged phasmid forms but does not directly measure capsid size or packaged-DNA copy number.

A. B.

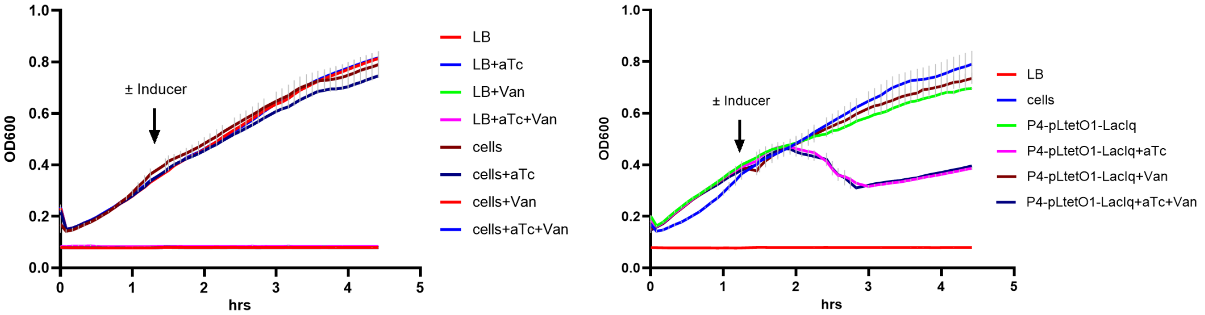


**Supplementary Figure S4.4 | Inducer toxicity and expression-control assays. A,** Testing for inducer toxicity. **B,** Testing inducer effects using the positive-control phasmid pP4-pLtetO1-LacIq (standard-size). LB = Luria Bertani; aTc = anhydrotetracycline; Van = vanillic acid; cells = *E. coli* Marionette; P4-pLtetO1-LacIq (standard-size) = Marionette cells containing the pP4-pLtetO1-LacIq phasmid; Inducer = aTc, Van or both. A 10 µL inducer volume were added at OD600 ~ 0.4. (n = 3).

#### Source blot and membrane images

**Supplementary Figures S4.**5-S4.12 show source Western blot images and Figure 10B source-analysis panels for Figures 5C, 5D, 10B, 14A and 14B. These panels are included for transparency and are not used to make new quantitative claims beyond the semi-quantitative analyses stated in the manuscript.

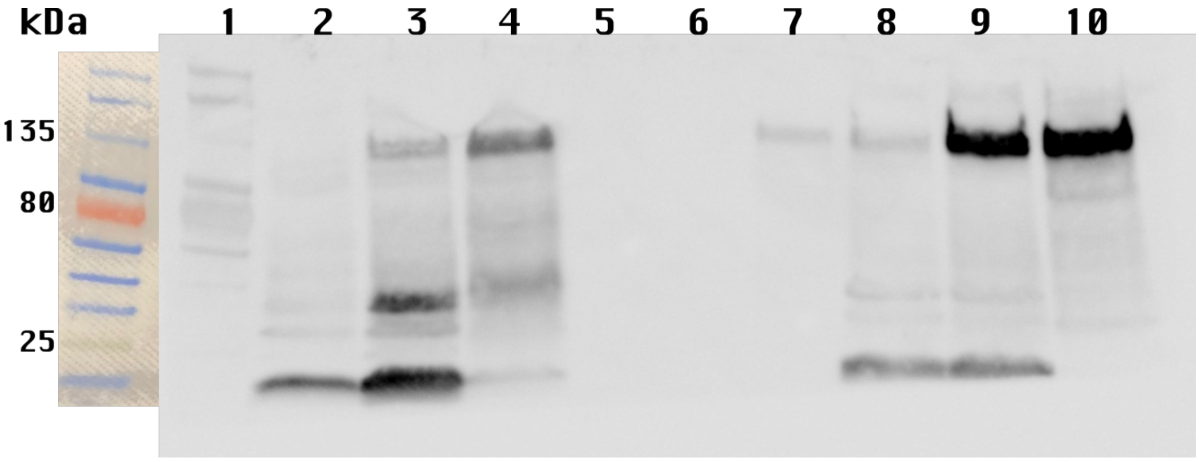


**Supplementary Figure S4.5 | Source image for the Figure 5C p72 MOI Western blot.** Concentrated *E. coli* EMG2 supernatants after transduction with P4-LacIq-p72 or P4-pLtetO1-p72 at the indicated MOIs and times; membrane/lane edges retained where present.

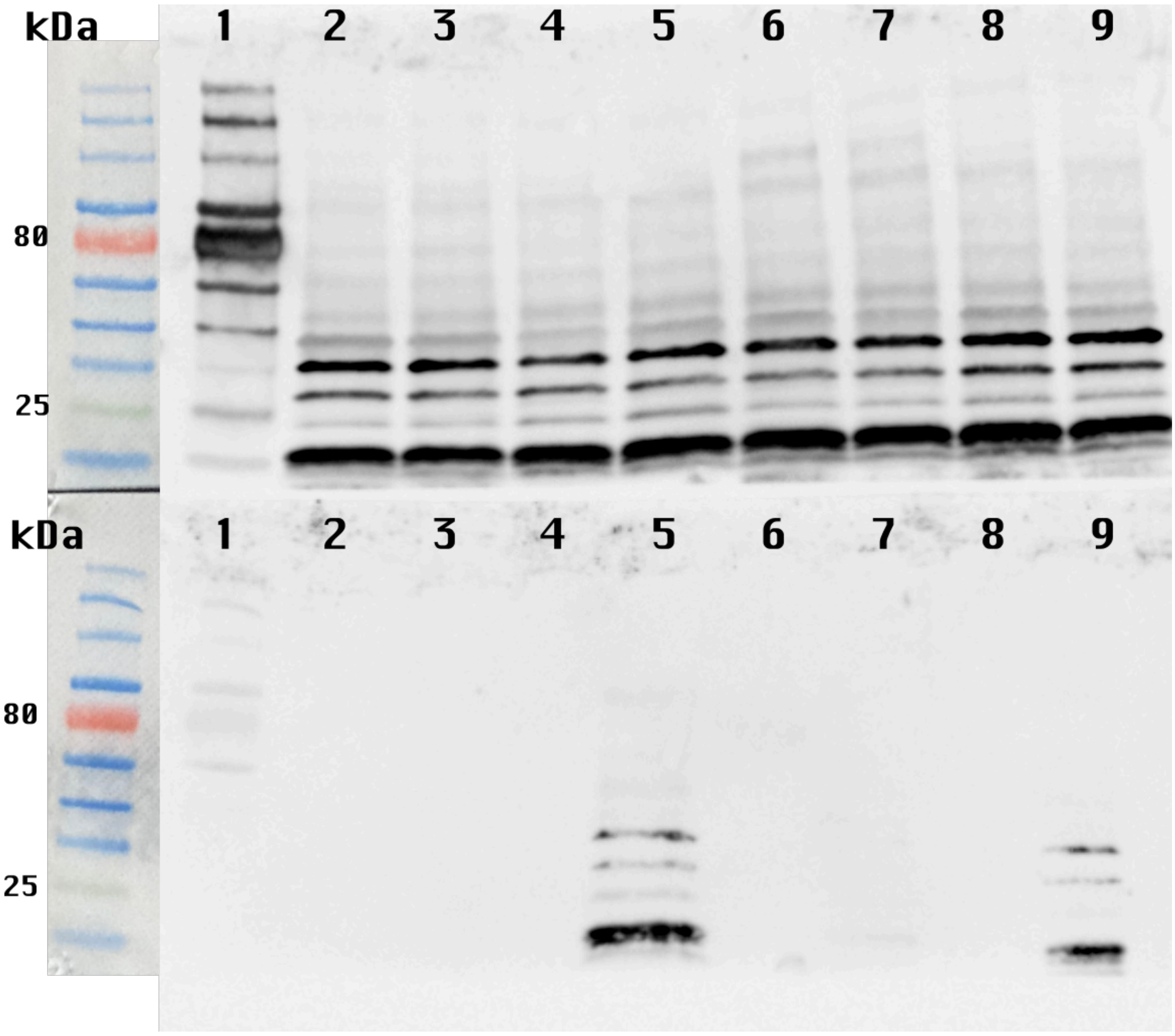


**Supplementary Figure S4.6 | Source image for the Figure 5D p72 and p65 Western blot.** Concentrated *E. coli* EMG2 supernatants after transduction with P4-LacIq-p65, P4-pLtetO1-p72 or P4-pLtetO1-p65 at the indicated times; membrane/lane edges retained where present.

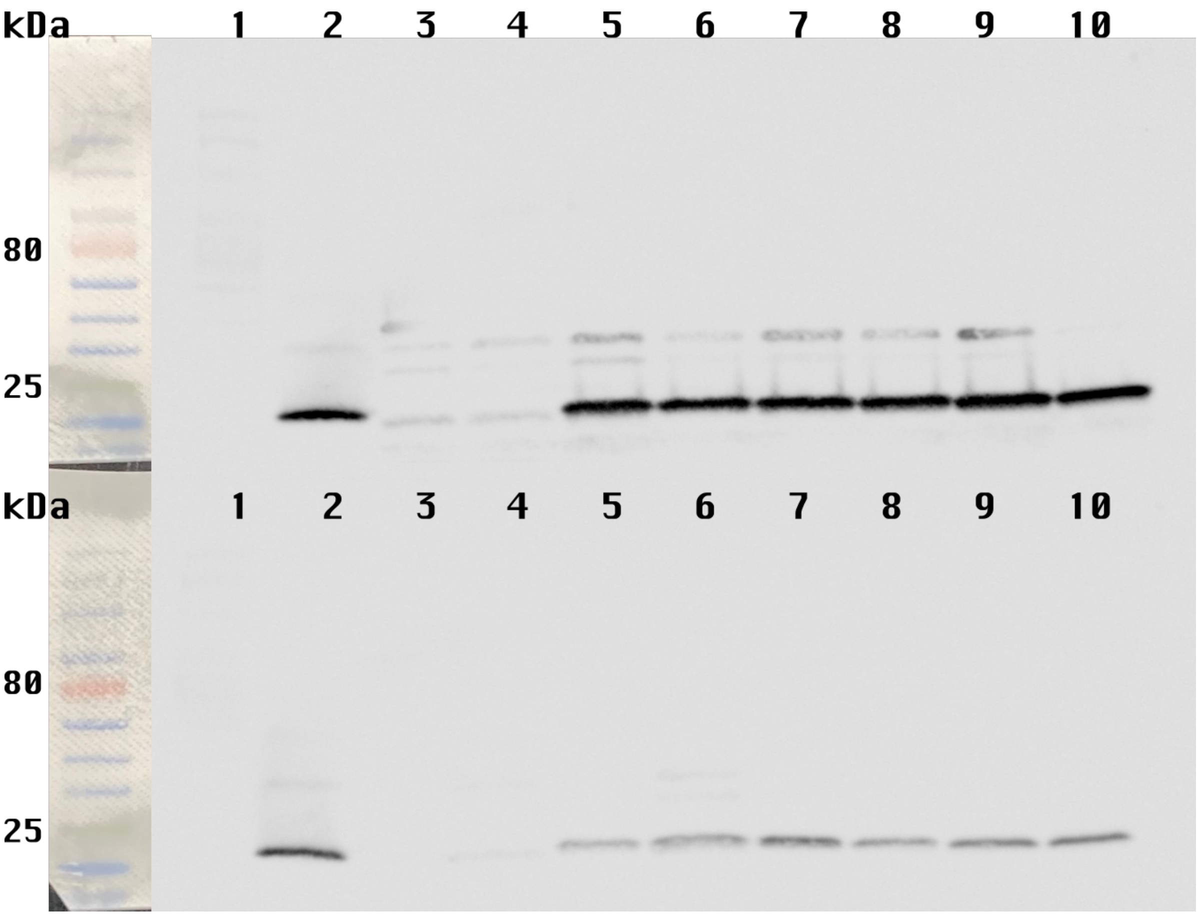


**Supplementary Figure S4.7 | Source image for the Figure 10B PE Western blot, 30 min antibody-exposure version.** PE detection in *E. coli* EMG2 supernatants at perceived half-time and full-time lysis after transduction with P4-DiffJJ-Test, P4-DiffJJ-LacI, P4-pLtetO1-LacIq or P4-pLtetO1-tac; lane 2 = 300 ng PE control.

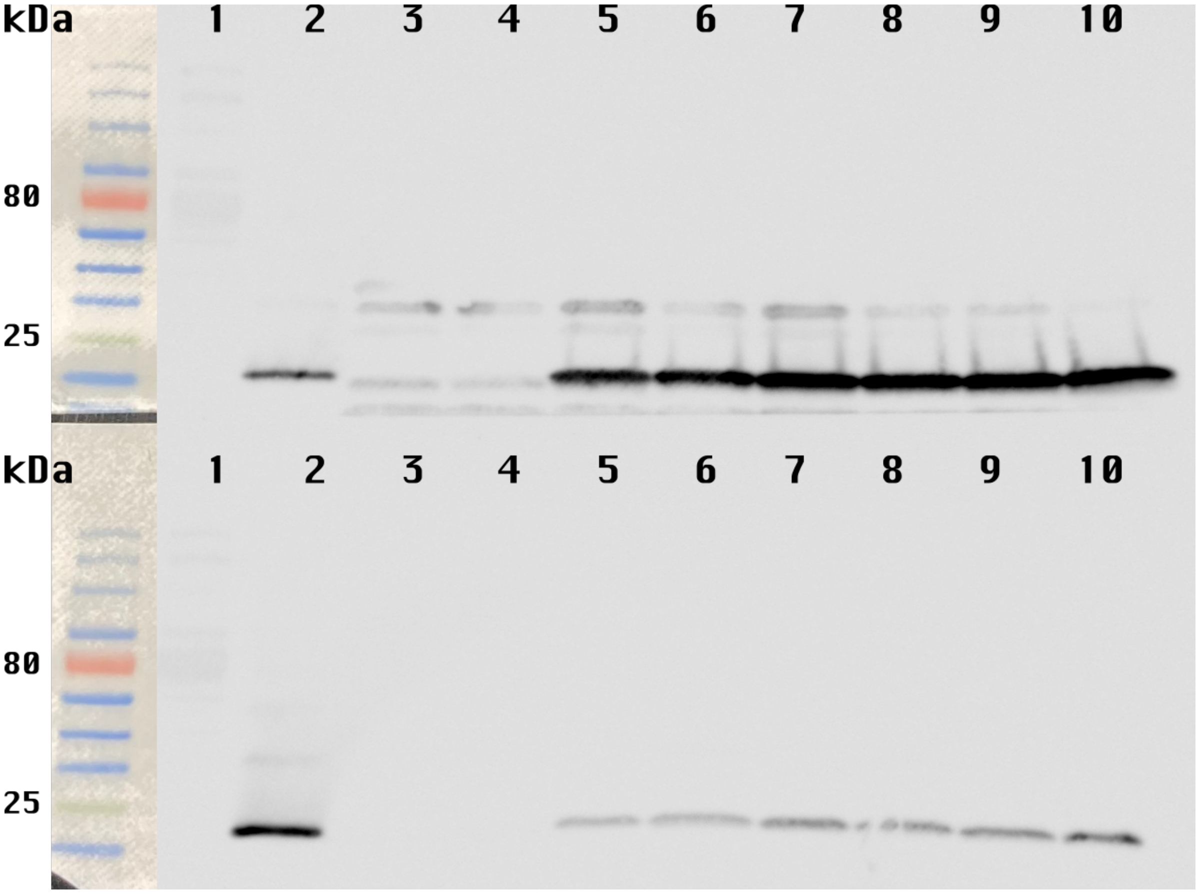


**Supplementary Figure S4.8 | Source image for the Figure 10B PE Western blot, 1 h antibody-exposure version.** Same Figure 10B lane set as Supplementary Figure S4.7 after longer antibody exposure; PE control retained.

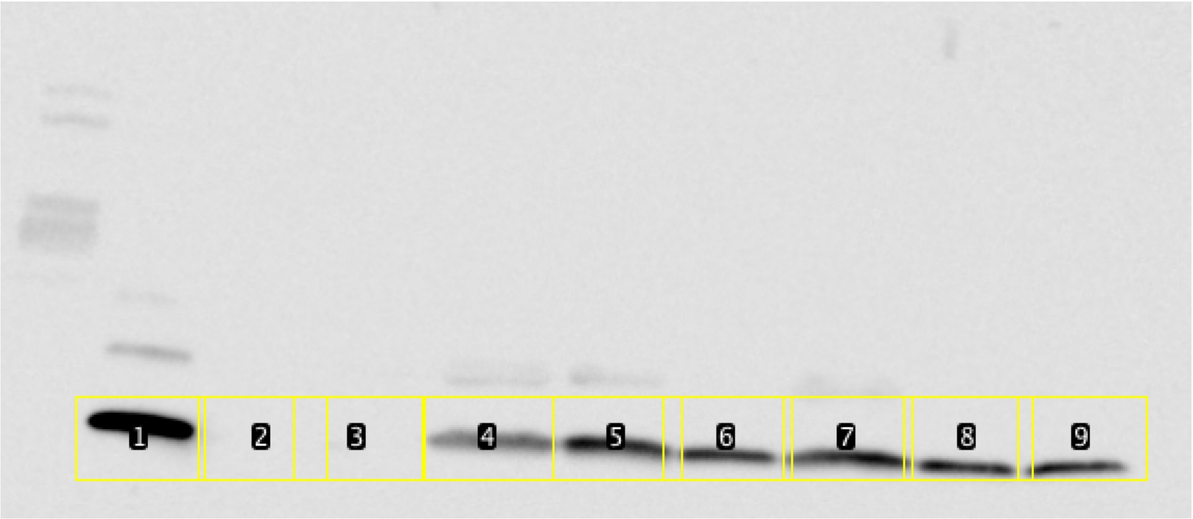


**Supplementary Figure S4.9 | Figure 10B ImageJ/Fiji quantification support panel.** Semi-quantitative Figure 10B band-density analysis by ImageJ/Fiji area-under-the-peak measurement, using the 300 ng PE control in lane 2 and normalization by MOI.

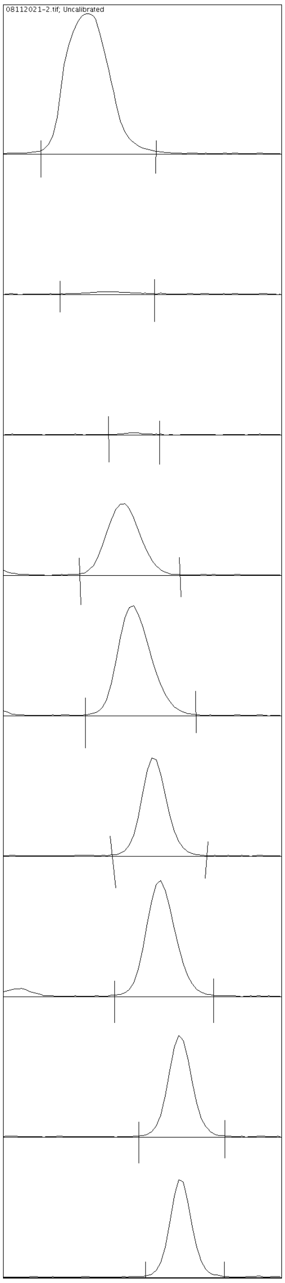


**Supplementary Figure S4.10 | Figure 10B Fiji gel-analysis trace export.** Band-density profiles used for the ImageJ/Fiji quantification support panel in Supplementary Figure S4.9.

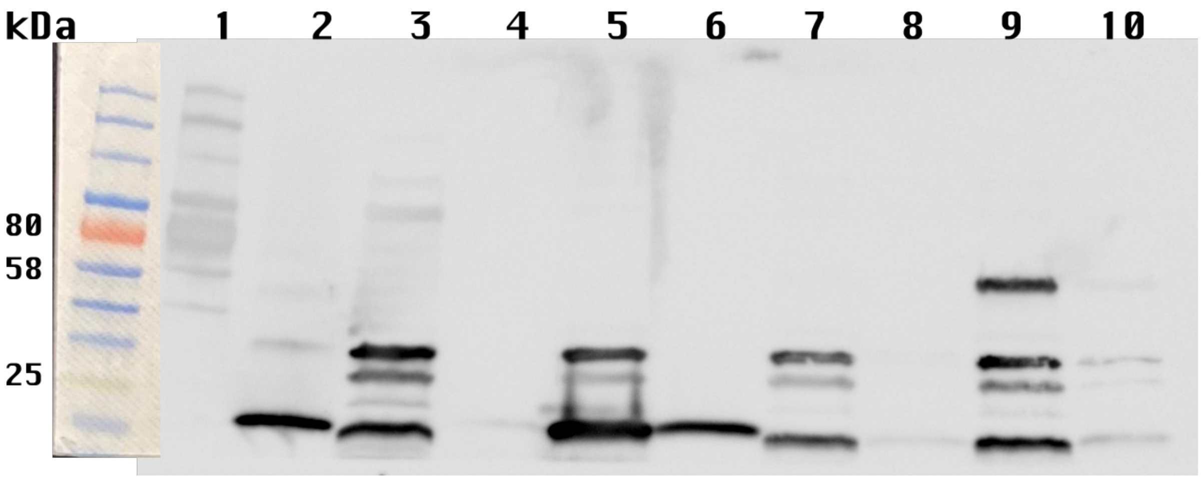


**Supplementary Figure S4.11 | Source image for the Figure 14A FliC-PE Western blot.** FliC-PE detection in pellet and supernatant fractions from *E. coli* Marionette cells carrying pP4Δ*sid*-pLtetO1-LacIq or pP4Δ*sid*-pLtetO1-pVanCC after aTc/Van induction and sampling at perceived half-time lysis; membrane/lane edges retained where present.

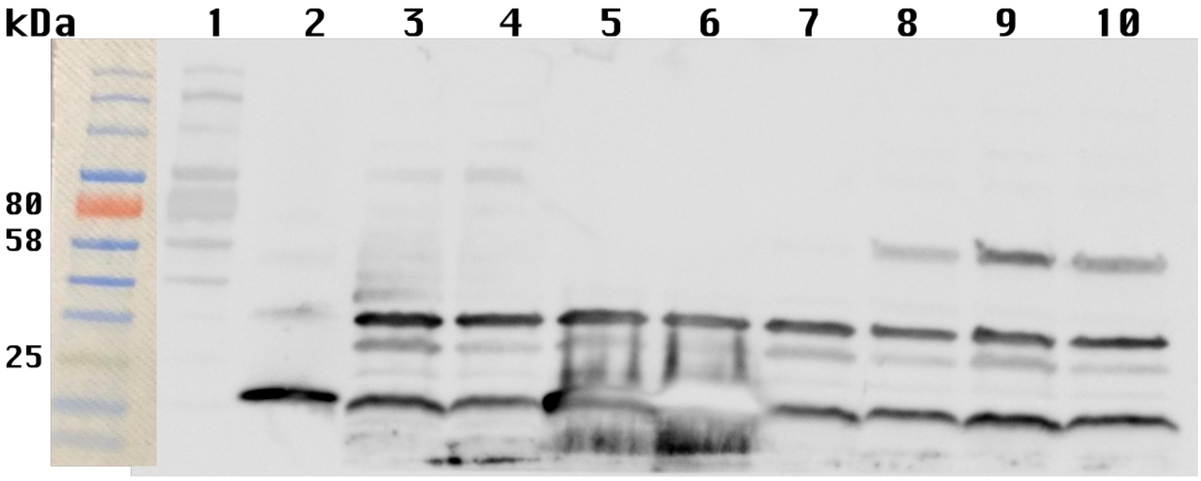


**Supplementary Figure S4.12 | Source image for the Figure 14B FliC-PE Western blot.** FliC-PE detection in pellet fractions from *E. coli* Marionette cells carrying pP4Δ*sid*-pLtetO1-LacIq or pP4Δ*sid*-pLtetO1-pVanCC after Van induction at 3 h and overnight time points; membrane/lane edges retained where present.

#### Supplementary Tables

**Supplementary Table S1. Structural P4 characteristics, predicted or literature derived.** § = the number of monomer copies was inferred by sequence homology; $ = the § was backed up by experimental data; ND = not determined.

| Mature P4 Structural proteins | Monomer mass (kDa) | Oligomeric state | No. of monomer copies | Location and remarks | References |
|---|---|---|---|---|---|
| **gpQ** | 39.11 | dodecamer | 12 | Capsid-tail junction | Doan & Dokland, 2007 [1] |
| **gpX**$^{§,\ Mu\text{-}Mup46,\ T4\ gp25}$ | 7.08 | ND | 6 | Baseplate-tail sheet interface | Christie & Calendar, 2016; Büttner *et al.*, 2016; Leiman *et al.*, 2010 [2]–[4] |
| **gpV** | 22.24 | trimer | 3 | Spike | Christie & Calendar, 2016 [2] |
| **gpW**$^{§,\ T4\text{-}gp25}$ | 12.63 | dimer | 6 | Baseplate wedge | Christie & Calendar, 2016; Leiman *et al.*, 2010 [2], [4] |
| **gpJ**$^{\$,\ Mu\text{-}\ Mup47}$ | 32.78 | ND | 12 | Baseplate wedge | Büttner *et al.*, 2016 [3] |
| **gpI**$^{§,\ Mu\text{-}Mup48}$ | 19.58 | dimer | 12 | Baseplate wedge | Büttner *et al.*, 2016 [3] |
| **gpH** | 71.50 | trimer | 18 | Tail fiber | Christie & Calendar, 2016 [2] |
| **gpFI** | 43.14 | hexamer | ~200 | Tail tube | Lengyel *et al.*, 1974 [5] |
| **gpFII** | 19.06 | hexamer | ~200 | Tail sheet | Lengyel *et al.*, 1974 [5] |
| **gpT** | 86.52 | ND | 5 | Tail tube base | Lengyel *et al.*, 1974 [5] |
| **gpU**$^{§,\ T4,\ gp48/gp54}$ | 17.45 | ND | 6 | Baseplate-tail tube junction | Maxwell *et al.*, 2013 [6] |
| **gpD** | 42.79 | trimer | 3 | Baseplate hub | Christie & Calendar, 2016 [2] |
| **gp psu** | 21.36 | dimer | 60 | Capsid decoration | Kizziah, Rodenburg, & Dokland, 2020 [7] |
| **gpN*, h1, h2** | 36, 39, 38 | hexamer | 180, ratio 9:1:2 | Cleaved major capsid protein | Dokland, Lindqvist, & Fuller, 1992 [8] |

**Supplementary Table S2. Buffer formulations used for the isoelectric-point precipitation and short-term stability experiment.**

| Buffer number | 20 mM Buffer (+ 50 mM NaCl) | pH on graph | pH measured |
|---|---|---|---|
| 1 | Tris | 8 | 7.87 |
| 2 | Bis-Tris-Propane, HCl adjusted | 7 | 6.95 |
| 3 | Bis-Tris, HCl adjusted | 6 | 6.03 |
| 4 | Bis-Tris, HCl adjusted | 5 | 4.93 |
| 5 | 0.5 mL Buffer #10 + 4.5 mL Buffer #4 | 4.9 | 4.86 |
| 6 | Bis-Tris, HCl adjusted | 4.7 | 4.67 |
| 7 | 1.5 mL Buffer #10 + 3.5 mL Buffer #4 | 4.4 | 4.44 |
| 8 | 1.8 mL Buffer #10 + 3.2 mL Buffer #4 | 4.2 | 4.17 |
| 9 | 2 mL Buffer #10 + 3 mL Buffer #4 | 4 | 4.01 |
| 10 | Bis-Tris, HCl adjusted | 3 | 2.94 |

**Supplementary Table S3. Primers for cloning.**

| Construct | Description | Primer name | 5'-3' |
|---|---|---|---|
| **pP4-Δlys** | Open the pP4-pLtetO1-LacIq vector to delete the | P4_T7gp10term_Fwd | `ctagcataaccccttgg` |

| | | | |
|---|---|---|---|
| | lysis cassette; with 3' overhang | P4_term_termover_Rev | aaggggttatgctagGGACCAAAACGAAAAAAGGCC |
| **pP4-LacIq-p72** | Amplify the p72 fragment from pet24-p72 vector; with 5' overhang | P4_RBSover_p72_Fwd | GAGGAGAAAGGAGATcatatgaaaaaggccctgg |
| | | p72_Rev | ctcgagttattaatgatggtgg |
| | Open the pP4-Δlys vector; with 5' overhang | P4_p72over_PE_Fwd | cattaataactcgagACAAAGGTTTATACGTGTATCCG |
| | | P4_RBS_Rev | ATCTCCTTTCTCCTCTTTCGAATTC |

**Supplementary Table S4. Tecan plate-reader programme used for OD600 kinetic measurements.**

| Programme section | Parameter | Setting |
|---|---|---|
| Plate setup | Plate model | [GRE96ft] Greiner 96 Flat Transparent |
| Plate setup | Plate cover | Yes |
| Plate setup | Barcode | No |
| Plate layout | Measurement wells | User-selected wells matching the experimental plate layout |
| Temperature control | Mode | On |
| Temperature control | Set point | 37.0 °C |
| Temperature control | Start condition | Wait for temperature before measurement |
| Temperature control | Accepted range | 36.5-37.5 °C |
| Kinetic acquisition | Number of cycles | 150 |
| Kinetic acquisition | Interval | 00:05:00 (5 min) |
| Absorbance measurement | Wavelength | 600 nm |
| Absorbance measurement | Bandwidth | 10 nm |
| Absorbance measurement | Reads per well | 10 |
| Absorbance measurement | Settle time | 0 ms |
| Absorbance measurement | Data label | OD |
| Absorbance measurement | Beam diameter | 3800 µm |

| Programme section | Parameter | Setting |
|---|---|---|
| Absorbance measurement | Multiple reads | Filled-square 2 × 2 pattern; 1000 µm well border |
| Shaking | Duration | 60 sec before each read |
| Shaking | Mode | Orbital |
| Shaking | Amplitude | 1.5 mm |
| Shaking | Frequency | 351.9 rpm |

**Supplementary Table S5. Primers for sequencing the cargo region.**

| Name | 5'-3' |
|---|---|
| P4_col1_F | catacctcgttaatacattggagc |
| pP4_Del2_Rev | cagggttttccttaactttgattccattcttttgtttg |

**Supplementary Table S6. Predicted expected sizes for FliC-PE and its variants.**

| Construct | FliC-PE | FliCΔ188-346-PE | FliCΔ191-286-PE | FliCΔ191-352-PE | FliCΔ220-320-PE | FliCΔ191-278-PE |
|---|---|---|---|---|---|---|
| Size (kDa) | 68.34 | 52.31 | 58.47 | 52.38 | 58.15 | 59.27 |

**Supplementary Table S7. Primer sequences.**

| DNA name | 5'-3' |
|---|---|
| delta_Fwd | ATGATTTACTGTCCGTCGTGTGGACATGTTGCTC |
| P4_delsidall_deltaover_Rev | cggacagtaaatcattttccggctcctcgtcacttcaggttaagaaaatt |

**Supplementary Table S8. Promoter sequences used in this study.** Sequences are shown 5'-3'; spaces are inserted every 10 nt for readability.

| Promoter | Sequence (5'-3') | bp | Source / note |
|---|---|---|---|
| $P_{lacI}{}^{q}$ | CGAATGGTGC AAAACCTTTC GCGGTATGGC ATGATAGCGC C | 41 | lacIq custom/source-feature sequence; detected in pAJ1053 source map in reverse-complement orientation. |
| $P_{lacI}$ | GACACCATCG AATGGCGCAA AACCTTTCGC GGTATGGCAT GATAGCGCCC GGAAGAGAGT CAATTCAGGG TGGTGAAT | 78 | lacI standard source-feature sequence. |
| $P_{tac}$ | TGTTGACAAT TAATCATCGG CTCGTATAAT GTGTGGAATT GTGAGCGCTC ACAATT | 56 | pTac source-map sequence with symmetric lac operator. |
| $P_{LtetO-1}$ | TCCCTATCAG TGATAGAGAT TGACATCCCT ATCAGTGATA GAGATACTGA GCACATCAGC AGGACGCACT GACC | 74 | PLtetO-1 source-map sequence; modified phage lambda PL promoter with tet operator sites. |
| $P_{DiffJJ}$ | ACTCTATCAT TGATAGAGTT TGACCTCCCT ATCAGTGATA GAGATACTGA GCAC | 54 | pLtetO12 vJJ non-sym weak 0.14x truncated custom feature; JJ weaker variant of PLtetO-1. |
| $P_{VanCC}$ | ATTGGATCCA ATTGACAGCT AGCTCAGTCC TAGGTACCAT TGGATCCAAT | 50 | pVanCC custom source-feature sequence; evolved Marionette promoter. |

For Supplementary Tables S9-S22, nucleotide sequences are shown 5'-3'. Only nucleotide sequences are set in Menlo. Feature colours mark curated genetic parts from source annotations and promoter-feature files; uncoloured positions are outside the curated feature set.

**Supplementary Table S9. pP4 wt sequence (5'-3'; 11,624 bp; cargo region highlighted).** Feature key: cos packaging region; P4 gop promoter; cargo region; P4 attP site; P4 replication origin; P4 crr replication region; P4 alpha terminator; P4 alpha; P4 epsilon; P4 cI; P4 pLE promoter; P4 gp11/vis; P4 pLL promoter; P4 *sid* promoter; P4 *sid*; P4 delta; P4 psu; P4 tsid terminator

```
GGCGAGGCGGGGAAAGCACCGCGCGCAAACCGACAAGTTAGTTAATTATTTGTGTAGTCAAAGTGCCTTCAGTACATACC
TCGTTAATACATTGGAGCATAATGAAGAAAATCTATGGCCTATGGTCCAAAACTGTCTTTTTTGATGGCACTATCCTGAA
AAATATGCAAAAAATAGATTGATGTAAGGTGGTTCTTGTCAGTGTCGCAAGATCCTTAAGAATTCGTGGCATGAGAGAGT
TAAAGGATGCTGAATCATGTATATGGATTAATTGGCGTTGTTGGGACCATAATCACTATTGCTACTAGCTTCATCCGCAA
CCAAGATATTTCACAATGGTTGCTCGTTTGTTCTGGATGGTTAGCTGCTTTACTTATTGGATGGTTTACACACAGAACGA
TTAAAGCAATTAGCAACAACCACACGAATGTTATAAAAAGTAATATGGAGGTAATCAACAGTCATAATGAATCCAATCAA
AACTTGATGGCTGAAAACAAAGAATTGATTAAAGAACTTGCAAGAACTTCAGAACAGAAAGAGAAGATGGAAAGTATAGC
TGCTTATCTTGCTACTCAGAACCCCCAAATTAACGCTATGCCAAGAACAGCAAGCCGACCAGAAAATATCGATTCGGAGG
CTAATTAAATATGAAAGTATATTTTGAAAATTACTCATACTATCCAGCCCTAAGAACACGATCAGCAGAAATGACCGGAC
TGAATAATTTATCATATGAGAATAAGAAAAAAATATTACCTTTAATTTCTTTGGGGAAATGGCCTCGCTCGGAAGAAATA
CAGGTATCGTTAGATAAAAGTTTAGAAGTGATGAGTAATCTCCCATTTATTTTGGATGTCACAAAGGATAATTCTCATCA
TTGCGCGTCAAGCTTCGAACTACTCTCTCCTGAGAATGGTTTTAAAAACTGGATAGAGTTTTGTTCCAGAAATGATAATA
TTATTCCAGTAGTCCAGATGCCGGATTCAGCAAAACTTAGAGATATATCTATCCAAGCCAGGGTTTTAGAGGAACTTAAA
GGTTCAATTGCATTTAGAATTAGAAATCTAAATACAGACATTAATAAAACCCTGACGTCACTAGTATCGATGAACTCTCC
AGAAAATGCCATAGTTTTCATAGATTTGGGCTACATTAGAGGGAATGTATCTGCTATAACAGCTGCAGCAATAAATTCGA
TTAATCAGATTAGAACTGAGATTCCTGAAGCCATAATTAGTGTGCTAGCAACAAGCTTCCCTAGTTCAGTTACAAATTTT
TGCCGTGAAAATGGACAGTCCGGATATATTGATGTTATTGAGAGAGAATTGCATCAAAATATTGGTGGAAGTGATGTCGC
TATTTATGGCGATCATGGTTCAATTCATTCAGTAGTGTATGACAACATAATTGGGCGATATGTTCCTAGAATTGATATAG
CGTTAAATGATTCATGGTATTTCGAGCGCCGCCCTGGGATGAATAAAGAAGGGTTCATTGAAGCTGCGAAATCAATTTTG
GCAGAGTATCCACATTATCAAAGGGAGGACTCTTGGGGAGCTGCAATGATAAGAAATGCTGCTATAGGGGATATAGCAGG
TATGGGAAGTCCAGCTAAATGGATTGCTGTTCGAGTGAATCTACACTTAAATAAGCAAATTGAGCTATCTGAAGCTCTAC
AGTATGGCTTTGATCATGATGAGGAAGATCTTATCTAATAAAAAACTAGGGAAGGTTAGCCTTCCCTAGCCCACAACCCT
TTTAAGTACTTTTTCCGGAATATAGGAAACATCATCGCCATATTTCCTAACTATAGAGTCAACATCAACCTTATGGTATT
CATTTAAAGATTTCTCCGAACAAGCCTTAAACTCAAACTTAACCAAAGAATTATTCCTTCTTTTAGCTTTGCTCAATAAG
GCAAGATCGTTTATAGATATTTCTTGATTACCATGGAGTCTAGATAAAAAATCATTAGATGTCGCATGATACCTATTTTT
AATAGATGTAATGACAAATTCTCTAATCTTGTTTAAAGATAACTTAGATGCCAAAACCTCAAGACAATCCCTATGTAGTT
CATTTGAAAATGTAATTCCATACTGATTAAGCAGATCTTTAAGCTCAGACTTAAGCAAAAAACCCAAATAAACAAATTTA
TTTAAGCGAGAGCAAACCCTTCCTTTTTGAATTATTTTAAAGCGTACGCTACTATTATTAGATTGAAATTCTACAACGCC
AACATCTGATGAAACCCGTTTAGATACTTCACTGGTAAATTTTGGCGCGCACACAATTGTAACTTTTTCGAAGCTAGATT
TAAAAATTTTTAACTGACCATCCAAGCGCTTAAGAGAGTCAAAATCGCTTTTTATTTCAAAAGCTTGTAGGCTTCCATTG
GCAATAGCCAAATCAGCTCTTCTGCTCCAGTTAGCAACAACCATTTCATTAATGATCGTTGCATCACTAAGCATGCCTTT
TTTATATAACCATTCAATTAGGGCTATTTTAACTTCCTGCTCTCGCAATAACTCAGCCATGTTAATCACACTCACTTCAC
GTTGATTTTTATTATAGCAACTAACAATTAAAAACTCCAGATTATGTTACTTAACACTCATTTTAAAAATCAACCAATAA
CTATTTTTATTTAAGTATTTCTTTGTATTTTGAATAATCATAAGGCTAATATATTTATTACGGTTCATATCCAAATAATC
AGCCCACCACTGCATCATTAAACGCCGCTCATCCAAATGCTCGGAGGTATGGATATATGCCGCACGTACATTATTACGCT
CTGAGTGGCTCAGTTGCCTCTCTATCGCGTCATCACTCCATAACCCCGACTCCCCCAACGCACCACGCGCCATAGTCCTA
AACCCATGCCCACACACCTCGGTTTTAGTATCATAGCCCATCGCACGCAATGCGCTGTTTACCGTGTTTTCACTCATAAC
CTTAGTTGCGTCATGATCCCCCGGAAAAAGCAGCTCTTTATCACCACTAATCTGCTTTAACTGGTTTAATAAAATCATCG
CCTGTCGACTAAGCGGAACGATATGTTCCTCTTTCATCTTCATGCCTCGGTACGAATAACGCACACCTTTAATTTCTTCT
CGCTTTGCAGGTATACGCCAAAGAGATTTATCGAAGTCGAATTCATCCCAACGTGCGAAACGTAATTCACTGGAACGCAC
AAAAGTTAGCAAGGAAAGCTTGACCGCAATCCGTGTCATTACACGGCCACGATATGCAGCAAGACGTGCAAGAAACTCAG
GGAATCGGCTAGAAGGTAAAGCAGGGTAATGTCGCGCTTTGGTTGTCGATAGCGCACCAGCCATATCGCTGGCTGGATTT
GAGTCGATGTAATCGTTCTGTACGGCATAACGCATAATGGCTGTGACGCGCTGTTGCAGGCGCTGAGCGACATCGTGTTT
GCCACTGGCATCAACTTTTTTAATCGGGGCTAACAGGTGGCTGGTTTTAAGCTGCCGAATGTCGGACGAACCGATATGAG
GGAATATATAAAGCTCAAGATAGCGAAGAACGCGCGATCGATGGTCTTCACTCCAGCGCTTGTTACTGGCATGCCATTCT
CGGGCGATAGCTTCAAAAGTATATGCCCCCGAATTCTCGGCCCGAGCTTCTTTTTGTACGACTTTTGGGTCTATGCCCTG
TACTAAAAGCTTTTTAGCTTCATCGCGTTTTGCTCTTGCCTGAGCAAGCGTCACAGTAGGCCAAACACCAAACGCAAGAC
GATCCTCTTTTTTGTCAGAGGGACGTCTGTATTTCATGCGCCAGTATTTGGAACCTTTGGCCGAAACCTCAAGATACAAA
CCACCACCATCGGCCATTTTGTAGGTTTTGTCTTTTGGCTTTGCGGTCTCGACCTGTCTGGCGTTGAGCTTCATTTGGGG
GCACATTTCTAATCGAAGTTAAGATGCCCCCAATTATGCCCCCAATGATATCCGGATTTCAACGGACAACCTCGGAAGAC
GCAGGACGAAAAAATCGCTGCAAGCATTGATTTTAAAGGGATATTTGGACTTTCTCGGATGGTCTTGGAAGTAAGAATGG
TGCCGAAGGCCGGACTCAAACATCAAAATAAGTTAATGATAAAAAACAAATAATAAAACACAACAATGAAATATGCCCCC
TTTTGTGCCCCCACTGTTTTTCTGACCAATCTATTTTCAGCCCATCAATAAATCGGAAAGTTAAATCATTTTTAATCAGT
AAGTTTGGATCCGTAGCTCGGATCCAAACCAGTGCATCTTTTATCCACATAAAAAATTTTTTTTCGAAAGAACTGTTCAC
ACTGTTCACCTTTCTGTTTTCTCCTTTTATTTCAGAGTGATAGGTGGTGAATAATGGGTGAAGGGTGAACATTCGATTCT
TCACCTCCGGCATTCTGCCGATGTGACTCATACCGGTGATTAATCCTCCGCACTGAAATCACTCAGGAAGAAAAAAGTTT
TTTTTGATTTGATTGTTCACACTGTTCACCTTTCGTTTTTCTCTTTTAATTTCAGTGTGATAACGGGTGAATATACGGTG
AAGGGTGAACAGTGGATTGTTCACCTTCGGGGGATATCGGGATAAAAAAAGACCGGCAGATGCCGGTCAGGTGGGTCAGG
CTGTTGTAGGGTCGTCACATTTTGGCAGCCAGTCGCCGTAGCTTTCCTCTTTCAGCGTCAGGTTGGTCTGTATCCCCTGT
TTGGTATGGCGTTTCTCGTAATTCAGTCCGTATTCCTTCAGCATCACCGGCAGCCCCAGCCCGAACATTTTCAGACTGAG
TACATTCCGGTAGCCGTTTGCCTCCATGTAGGCCAGATAGGCGTGATAGAGGTATTTACGGTAATTGCGCGGGATGATAC
TGGCGTTCCCCATATACATGCCGCTGGTCTGCGGCAGGGTTTCCAGATAGCCGATAAAATCAAACGTCGGGTCGGCATCC
CGTTTGATGTTCAGTGCCTCGTCTGAGTTCTGCTGGGACTGAAGCAGTGACCGGGCGAGCATCGGGTCGCTGAACTTCTG
CATCAGGTGACGCACGATGACCGCCAGCTCGCGGGTGATTTTGTCCTTAAGCTGCGGGTCGCGCTCCTGCGGGGCTATCT
GTTCCGGGAAGTGAATAATCACCCGTCGGCGTGACACGCCGCCGCTGCGGTCGGTGAAGCGCATCGGGTTATTGTTCACG
GCCAGAATCACCGCCGGGATGTGCGTGGAGTACGCATCCCGGTATTTCGGGTCAACGGACACCGCATCGCCGCCGGTGAT
GGCCTTGAGTCCGGCACCGTCGCCGCTCCATTTTTCCTGGTCCGGCAGGCGTATCAGTGAGAAGCCAGTTAACGCGGCAC
GTTCACGCGGGGATTCCAGCGTCTCGATGGTGGCCGACGTGGCGTTATCCTCCCCGGCCAGCAGGGTGGCTATTTCGGCC
ATGATACTTTTGCCGCTGCCGCCGGGACCGGTCACCTCCAGAAAGAGCTGCCAGTCGTAGCGGTTTGCCAGCACCATAAA
CAGTGCAGCCAGAATCACGTCGCGTTTTTCCGCACGGCCACCGGCGGCACGGTCAAGCCAGCGCCAGAAGGCGGGGGCGT
GGGTTTCCAGCGTTTCACCGTCCACCGGCGGGGTGAAATCCACATCGCACAGGGTGCGCATCCAGTGTGACGGACTGTGC
GGGTGGAACGTGCCGTTCTGCGTGTCGAGCACGCCGTTACGAAAGCCAATCAGGCGGCGGGAGGGGGCTTCCTGCTGCGG
AATAATCAGCTTCAGGGTATCCACCACGGAGGCCACCTTCCCGGAGGAGAACGGCGCACGCAGACGCTGAAACAGCCCGG
```

```
CCACATCCCGGGCAAAGTCCTGTGGCGGCAGCACCTTCCAGACACCATTTTCATAGCGGGACAGAAGCTGGCCGTTGGCA
TCGACCGCGAGCGCCTCGCCGTAATGCTCATAGATACGCATGGCCTTTTCGCTGGTACTCATGGCGGAAAACTCCGCTTC
GCTCATGGTGTCGAACGGGCTTTCAGCCGGTGGCCGGATGGCATCGTAAATGGCCTTACGGGTGGCCTCCCCGCCGTACT
GCGTGAAGGCATCATTCCAGTCACCGAAGACCGGCGGCAGGGCAACAACACCTTCACACGCATCTGCGGCTGCGGCGGCT
TTTTTCTGGCCGTCACCACTGAGGTCACGGTCAGCGGCAAGGACAATCTGACAGGCGGGGTGCTTCTGCCGGGCAAGGCT
GGCCAGAGAAAGGAGGTTCACGGAAGAAAGCGCCACCATCACCGTTTCACCGGTCAGGTGATGTACGGTAAGTGCGGTCG
CGTATCCCTCCGCTATCCACAGACGTTTTCCGGCCTGATTCTGTCCTTCAAGGGTGTGACAGGTGCCCCTGACCTGTCCG
CCTTTCAGGGTGCGCTTACGGCCGTCAGCACTGATTAACTGAAGGTTAACCAGTTCGCCGCTGTCGTCATACAGTGGCAC
CACAAGGTCACCGGCGCGCCAGCTCACGCCACCGGCTCTGTGTGTGCCGGTCAGCATCCGGCATTCCCGGCCGGGAAAGC
CCTTGCGGGTCAGGTAGGCGTTACCGGTTCCGGTACGGGTTTTCGCCATCAGGGTTTGTGCCAGTGCGGCGGCGTTCTTC
CGGGCAGCGTCTGTTTCATCAACGGCGGCGGTCGTCACTGCCGGGTCAGCCGGTGGCAGGCTGCCGGTCACGGCAGCCAC
CTTTGCGGCCGCGTCGGACGGGGAAACACCAAAAACCTTTTCAACCAGTTTCAGGCCGTCACCGGCACCACACTGATTGC
AGTACCAGGTGCCGCGCCCCTCCCTGTCATCAAAACGGAAGCGGTCACTCCCGCCACAGACCGGACAGGGCTGATGACGG
TTCTTCAGCACCTGAATCCCCAGCGCCGGGAGAATACGCGGCCAGTGGCCGAGCGCATGGCTGACGGTGGCGGTTACGTT
CATTTTCATGGTGTTGTTCTCCTTCAGTGCAGTACCGGCGCTTTTATGTGACGGGCACAGAGTTCATCCATCACAACCAG
CCCGAGAAAGGACAGCGACGGCGCGGCCTTCAGGGGGCCGGATTCCATTAAATCTTCCAGCAGGGCACAGGCTATCTGAC
GCCCTTTTTCCTCACCGTGCTGGCGCAGATAAAAGCCTTCCAGCTCAGCGGCGATGGCCGCCTCCAGTGACTCAAGGGTG
AGATGCGGGTAGCGGTGCTGACGTTCGCACACGGTCAGCCAGGCACAGGCGACAGCGCGACGGTAAAGGGCAGCGCGTAA
GACGGGCGGTAAGGGTGTTTTCATTTGCTTTTCTCCCTGTGACAGATGACTGCATTCCGTGCCGGTTGCATTAACTGATA
AGGCATATCTGCGTCTCCTGAAGACGTGCGTATCCCTGCGCGAATACGCACATTTAATTTTTCGGGGGTCGTTTTTTAAT
TACAGATAATTGCGGTAACTGTTATCCGGGGTGGTTTCCGGGTCAGGCTCCGTGCGGGGAATTTCCCGCCATTCCCGCGC
CACCGGTGCTGCCCGGCTGACCGGAACAGTGTCCTGCGGGTAAATATCCAGATATTTTTCCCGCCATTTCTGTAATTCCG
GGTCTCCGGCCATTTCTTTCAGTACCGCATGCCGGTTTACGGGGCTGCGTTTAAACAGGTCAGGACGGTCACAGGTAAAT
TCCCGCAGAAAACGCCCCAGCGGGATGTCTGTGGTGCGTCCGTCAGCGAGGATACGCACAAGGATACTGAATTTACGGCG
GTACGGGTTCCAGACAATGTCCGGGCAGCGGTACGGCATTTCCCACGGAATACCGTCTTCCAGAATGCCGACCACGGCCA
CATCGGGAAAACCGGCAGAACGGTAAATCTCACCGGGCTGGGGAAAATCAAACATGCGTCCTGTCTCCCCGGTCTTTCTG
CTGGGCGAGAAAATCGCGGCACAGGCCTTTGGCTTTCAGCTCATTCAGCACAAAATCAATATCTTCATTCAGGTAGCTGA
AAATATGTGGAATGTAGAGCTGATGCAGGCCGGAGAGTTCACGGTGAATCAAATCACCCCCAACAAACCGGGATACGGCG
CTGGCGCGGTTGAGCTTATGGTAAGCCTCAATGCTGAGGTGTTCACGGGCGTCATGACGCGCTGAGACGGTCTGAGGGGC
TTTTTTATTACGCACGGGACACCTCCACCACCGGCAGACGGGCAGCAAGGGAGAGCACATAGTCACGGACAAGGGAACGG
CGGGCACTGCGTTCATCACCGGCGACGGTGCGAAGCATACAGATACGGGGATGACGGTCTGCGCGACGGACAGCCGCAAA
CACAAAGACAAATTCAGGGTGTGAGGGGGTAAGGGTTGTAGCCATGATGGCAGCCTCCTGTGAATAGCAAATAACGCTAT
CGCCGGAGTTCTCACGCTCGATGGCGATAGCCCAGACGGGGGTGAGAATACCGGCTTCACAGGATACCGGCCAGCCCGGA
GGCTGCCCCGCCTGAGCTACCATTGACTCTGCGGCATAATGAGCGGACGCGGGCAGGATGCACGGAATGCCATCTGCACG
ACTGACCACACACCACACCATAATCTGGCGCTCTGTGGCATTGATTGCGACACAAAAAAAGACGCGTGGCGCGTCATATG
TCGCCTGTGAATTGCTCGGGTTCTCACGCCCGGCTGCCGATTTTGCGGCAGGCGAAAAACTATATCCGCAAATGCCGGAA
AAAGGCAAGCCAGAAAAAGGGAGTTTTTGCAGAGCGGGCATCATCATGCGTCGTACCCCCGTTTGCGTCCGGCAATGCGT
CCGGCCATCCATGCGGTGACTTCAGAGTGCAGCCAGGCCACATTTTTACCGCCAAGACTCACCTGCGGCGGAAATTCCCC
CTTACGGATGAGTTCGTAGATGGTCGAGCGTGACAGGCCGCACAGGTGCATCACTTCCGGCACACGTAAAAAACGCTCCT
GCGTGATGTCCGGCAGCGGCATCAGTGGCGTCACTGGGGCGGGAGACGGGGAAGAAAAAACAGCTTGCATCGGGCTACCT
CGTTAATGTCCATACAGCACCGGATAAGTCCGTCCGGCTTCGGGTAGCGCTTTATTTTGTGAATATTTTCAGCAGACGCA
ACAGGGGGGATTTGTTCAGGCTGTCTTACAATGGCTGTGTGTTTTTTGTTCATCTCCACTTAAAGTCATTTAAAGCCACT
TAAAGCAATTTGTAATTTTTATAGTGAAATACAAATCGTTTCTTCTTATTCATTCCCGGCGAATTAATAAAAACAAACAG
TAGTAAACAGCACAAAAAGCCCATCAACGGGTGAACAGTGGTGAACAGACGGTGAACAGTCATTACTGCGATTGTTCACC
CTTTAACTTACTGTATTACTTATCTTTTTTATTAAGGTGAACAGAGGTGAACAGTAAAATATAAAAAAACAAACAGTAAG
CCGGTTTTTCCTGCGACCTTTTCCTGGCTTGCCGGTCTGAGGATGAGTCTCCTGTGTCAGGGCTGGCACATCTGCAATGC
GTCGTGTTGTTGTCCGGTGTACGTCACAATTTTCTTAACCTGAAGTGACGAGGAGCCGGAAAATGTCTGACCACACTATC
CCTGAATATCTGCAACCCGCACTGGCACAACTGGAAAAGGCCAGAGCCGCCCATCTTGAGAACGCCCGCCTGATGGATGA
GACCGTCACGGCCATTGAACGGGCAGAGCAGGAAAAAAATGCGCTGGCGCAGGCCGACGGAAACGACGCTGACGACTGGC
GCACGGCCTTTCGTGCAGCCGGTGGTGTCCTGAGCGACGAGCTGAAACAGCGCCACATTGAGCGCGTGGCACGCCGGGAG
CTGGTACAGGAATATGACAATCTGGCCGTGGTGCTGAATTTCGAACGTGAACGCCTGAAAGGGGCGTGTGACAGCACGGC
CACCGCCTACCGGAAGGCACATCATCACCTTCTGAGTCTGTATGCAGAGCATGAGCTGGAACACGCCCTGAATGAAACCT
GTGAGGCGCTTGTCCGGGCAATGCATCTGAGCATTCTGGTACAGGAAAATCCGCTCGCCAACACCACCGGCCATCAGGGC
TACGTCGCACCGGAAAAGGCTGTCATGCAGCAGGTGAAATCATCGCTGGAACAGAAAATTAAACAGATGCAAATCAGCCT
CACCGGCGAGCCGGTTCTCCGGCTGACCGGACTGTCAGCGGCAACACTCCCGCACATGGATTATGAGGTGGCAGGCACAC
CGGCACAGCGCAAGGTGTGGCAGGACAAAATAGACCAGCAGGGAGCAGAGCTTAAGGCCAGAGGGCTGCTGTCATGATTT
ACTGTCCGTCGTGTGGACATGTTGCTCACACCCGTCGCGCACATTTCATGGACGATGGCACCAAGATAATGATTGCACAG
TGCCGGAATATTTATTGCTCTGCGACATTTGAAGCGAGTGAAAGCTTTTTCTCTGACAGTAAAGATTCAGGAATGGAATA
CATTTCAGGCAAACAGAGATACCGCGATTCACTGACGTCAGCCTCCTGCGGTATGAAACGCCCGAAAAGAATGCTTGTTA
CCGGATATTGTTGTCGGAGATGTAAAGGCCTTGCACTGTCAAGAACATCGCGGCGTCTGTCTCAGGAAGTCACCGAGCGT
TTTTATGTGTGCACGGATCCGGGCTGTGGTCTGGTGTTTAAAACGCTTCAGACCATCAACCGCTTCATTGTCCGCCCGGT
CACGCCGGACGAACTGGCAGAACGCCTGCATGAAAAACAGGAACTGCCGCCAGTACGGTTAAAAACACAATCATATTCGC
TGCGTCTGGAATGAGGGCTGCCGGTTAACACCGGCCGTCGCCGCACACCGTATTTTTATTCTTCAGCATGATGAGAAAGA
GATAACGATGGAAAGCACAGCCTTACAGCAGGCCTTTGACACCTGTCAGAATAACAAAGCAGCATGGCTGCAACGCAAAA
ATGAGCTGGCAGCGGCCGAACAGGAATATCTGCGGCTTCTGTCAGGAGAAGGCAGAAACGTCAGTCGCCTGGACGAATTA
CGCAATATTATCGAAGTCAGAAAATGGCAGGTGAATCAGGCCGCCGGTCGTTATATTCGTTCGCATGAAGCCGTTCAGCA
CATCAGCATCCGCGACCGGCTGAATGATTTTATGCAGCAGCACGGCACAGCACTGGCGGCCGCACTGGCACCGGAGCTGA
TGGGCTACAGTGAGCTGACGGCCATTGCCCGAAACTGTGCCATACAGCGTGCCACAGATGCCCTGCGTGAAGCCCTTCTG
TCCTGGCTTGCGAAGGGTGAAAAAATTAATTATTCCGCACAGGATAGCGACATTTTAACGACCATCGGATTCAGGCCTGA
CGTGGCTTCGGTGGATGACAGCCGTGAAAAATTCACCCCTGCGCAGAACATGATTTTTTCGCGTAAAAGTGCGCAACTGG
CATCACGTCAGTCAGTGTAAAATTCCCCGAAAATCCGCCCGTTTTTACTGAAAAAAGCCATGCATCGATAAGGTGCATGG
CTTTGCATGCGTTTTCCTGCCTCATTTTCTGCAAACCGCGCCATTCCCGGCGCGGTCTGAGCGTGTCAGTGCAACTGCAT
TAAAACCGCCCCGCAAAGCGGGCG
```

**Supplementary Table S10. PEwt sequence (5'-3'; 483 bp).** Feature key: PE antigen

```
ATGAAGAAAATTATTTTAACTCTGTCTTTAGGTTTACTGACCGCATGCAGCGCCCAGATTCAGAAAGCCAAACAGAACGA
TGTGAAACTGGCCCCTCCGACCGATGTGCGCAGCGGTTATATCCGTTTAGTGAAAAACGTGAATTACTATATTGACAGCG
AAAGCATCTGGGTGGACAATCAAGAACCGCAGATTGTGCACTTCGATGCCGTGGTGAATTTAGACAAAGGTTTATACGTG
TATCCGGAGCCGAAACGTTACGCCCGCAGCGTTCGTCAGTATAAAATTCTGAACTGCGCCAACTACCATCTGACCCAAGT
TCGCACCGATTTCTACGATGAGTTTTGGGGCCAAGGTCTGCGTGCCGCACCGAAAAAGCAGAAGAAACACACTTTATCTC
TGACCCCGGATACCACTTTATACAATGCCGCCCAGATCATTTGCGCTAACTATGGCGAAGCCTTTAGCGTGGATAAAAAA
TAA
```

**Supplementary Table S11. p65 model-protein fragment sequence (5'-3'; 633 bp).** Feature key: p65 model-protein fragment

```
ATGCTCTCCACACTCCGCCGCACTCTATTTGCGCTGCTGGCTTGTGCGTCTTTTATCGTCCATGCCGATCCGCCGGCCAC
CGTGTATAAATATGATAGCCGCCCGCCGGAAGATGTGTTTCAGAATGGCTTCACCGCCTGGGGCAACAATGATAACGTGC
TGGACCACCTGACCGGTCGTAGCGGTCAGGTGGGTAGCAGCAACAGTGCCTTTGTGAGCACCAGCAGCAGCCGTCGCTAT
ACCGAAGTGTACCTGGAACATCGCATGCAGGAAGCAGTGGAAGCAGAACGTGCAGGCCGTGGCACCGGCCATTTTATTGG
CTACATTTATGAGGTGCGCGCCGATAATAACTTTTACGGCGCCGCCAGCAGCTACTTTGAATACGTGGACACCTACGGCG
ACAATGCCGGTCGTATCCTGGCAGGTGCCCTGGCAACCTATCAGAGCGGTTATCTGGCCCATCGCCGCATTCCGCCGGAA
AATATTCGCCGTGTGACCCGCGTGTATCATAACGGCATTACCGGCGAAACCACCACCACCGAGTATAGCAATGCCCGCTA
TGTTAGCCAGCAGACCCGCGCCAATCCGAATCCGTATACCAGCGGCGGTCATCATCACCACCACCATTAATAA
```

**Supplementary Table S12. p72 model-protein fragment sequence (5'-3'; 2,184 bp).** Feature key: p72 model-protein fragment

```
ATGAAAAAGGCCCTGGCAACACTGATTGCACTGGCACTGCCGGCAGCAGCACTGGCAGAAGGTACCGAAAACGTGCAGTA
TCGCCACGTTGAACTGGCACGCGTTGGCCAATTAGTTGAGGTGGACACCCTGGAACACGTGCAGCATATTATTGGCGGTG
CCGGCAATGACAGCATTACAGGCAACGCCCACGACAACTTTCTGGCAGGCGGCAGCGGCGATGATCGTCTGGATGGTGGC
GCCGGTAATGATACCCTGGTGGGTGGCGAGGGTCAAAATACCGTTATTGGCGGTGCCGGCGACGATGTTTTCCTGCAGGA
TCTGGGCGTGTGGAGCAACCAGCTGGACGGTGGTGCAGGTGTGGACACCGTGAAATACAACGTTCATCAGCCGAGCGAGG
AACGTCTGGAACGCATGGGTGACACCGGTATTCATGCCGATCTGCAGAAAGGCACCGTGGAAAAATGGCCGGCCCTGAAC
CTGTTTAGCGTGGATCATGTTAAAAATATTGAAAATCTGCATGGCAGCCGTCTGAACGACCGCATTGCCGGCGATGACCA
AGATAACGAACTGTGGGGCCATGACGGTAATGACACCATTCGCGGCCGTGGTGGCGACGACATTCTGCGTGGTGGTCTGG
GTCTGGATACACTGTACGGTGAGGATGGCAATGATATCTTTCTGCAGGATGACGAAACAGTTAGCGACGACATTGACGGT
GGCGCAGGTCTGGACACAGTGGACTACAGCGCCATGATTCATCCGGGCCGTATTGTGGCCCCGCACGAATATGGTTTTGG
TATCGAAGCCGATCTGAGCCGTGAATGGGTGCGCAAAGCAAGTGCCCTGGGTGTGGACTATTACGACAACGTGCGCAATG
TGGAAAACGTGATTGGCACCAGCATGAAAGACGTGCTGATTGGCGACGCCCAGGCCAACACCCTGATGGGCCAGGGTGGT
GACGATACAGTGCGCGGTGGTGATGGTGATGACCTGCTGTTCGGCGGCGATGGTAACGACATGCTGTATGGCGATGCCGG
TAACGACACCCTGTACGGCGGTCTGGGTGACGATACCCTGGAGGGCGGTGCCGGCAACGATTGGTTTGGCCAAACCCAGG
CCCGTGAACATGATGTGCTGCGCGGTGGCGATGGCGTGGATACCGTTGACTACAGTCAGACAGGCGCCCATGCAGGTATC
GCAGCCGGTCGCATTGGTCTGGGTATTCTGGCAGACCTGGGTGCCGGTCGTGTGGATAAACTGGGTGAGGCCGGCAGTAG
TGCCTATGACACAGTGAGCGGCATCGAAAACGTGGTGGGCACAGAGCTGGCAGACCGTATCACAGGCGATGCCCAAGCCA
ATGTTCTGCGCGGCGCAGGTGGTGCCGATGTGTTAGCCGGTGGTGAGGGTGATGACGTTCTGCTGGGCGGTGACGGCGAT
GATCAGCTGAGTGGTGATGCAGGTCGCGACCGCCTGTATGGCGAAGCAGGCGATGATTGGTTCTTTCAGGACGCAGCCAA
TGCCGGTAATCTGTTAGACGGTGGCGACGGTCGTGATACAGTGGATTTTAGCGGTCCTGGTCGTGGCCTGGATGCCGGCG
CAAAGGGTGTGTTTCTGAGCCTGGGCAAAGGCTTTGCCAGCTTAATGGACGAACCGGAAACCAGCAACGTGTTACGCAAC
ATTGAGAACGCCGTGGGTAGTGCCCGCGATGACGTGCTGATCGGTGATGCCGGTGCAAATGTTCTGAACGGCCTGGCCGG
CAACGACGTGCTGAGTGGTGGTGCAGGTGATGATGTTCTGCTGGGTGATGAGGGCAGTGACTTACTGAGCGGTGACGCCG
GCAATGACGATTTATTCGGCGGCCAGGGTGATGACACCTACCTGTTTGGCGTGGGTTATGGCCACGACACCATTTATGAG
AGCGGTGGTGGTCACGACACAATCCGCATCAATGCAGGCGCCGATCAGCTGTGGTTTGCCCGCCAGGGTAACGATCTGGA
GATTCGTATCCTGGGCACAGACGATGCCTTAACAGTGCATGATTGGTACCGCGATGCCGATCATCGCGTGGAGATTATCC
ATGCAGCAAACCAGGCAGTGGATCAGGCCGGCATCGAGAAGCTGGTTGAGGCCATGGCCCAGTATCCTGATCCTGGTGGT
CATCATCACCACCATCATTAATAA
```

**Supplementary Table S13. FliC sequence (5'-3'; 1,515 bp).** Feature key: FliC flagellin protein

```
ATGGCCCAAGTGATTAACACCAACAGCTTGAGCCTGCTGACCCAAAACAACCTTAACAAAAGTCAGAGTGCCCTGGGCAC
TGCCATTGAAAGACTGAGCAGTGGCCTGAGAATTAATAGTGCCAAAGATGATGCAGCTGGCCAAGCCATTGCCAACAGAT
TTACAGCAAACATTAAGGGCCTGACCCAAGCAAGCAGAAATGCCAATGATGGAATATCTATTGCACAGACCACTGAAGGT
GCCCTCAATGAAATTAACAACAACTTACAGAGAGTGAGAGAACTGGCAGTGCAGAGTGCCAACAGCACCAACAGTCAGAG
TGATCTGGATAGCATTCAAGCAGAAATTACTCAGAGACTAAATGAAATTGATAGAGTGAGTGGTCAGACTCAGTTTAATG
GTGTGAAAGTGCTGGCACAAGATAACACCCTGACCATTCAAGTGGGTGCCAATGATGGTGAAACCATTGATATTGATCTG
AAACAGATTAACAGTCAGACCCTGGGCCTGGATACCCTGAATGTGCAGCAGAAATATAAAGTGAGTGATACAGCAGCCAC
TGTGACTGGCTATGCAGATACCACCATTGCCCTGGATAACAGCACCTTTAAGGCTAGTGCCACTGGCCTGGGTGGCACTG
ATCAAAAAATTGATGGTGATCTGAAATTTGATGATACCACTGGCAAATATTATGCCAAAGTGACTGTGACTGGTGGCACT
GGCAAAGATGGCTATTATGAAGTGAGTGTGGATAAGACTAATGGTGAGGTTACCCTTGCTGGTGGTGCCACAAGCCCACT
AACTGGTGGCCTGCCTGCCACTGCCACAGAAGATGTCAAAAATGTGCAAGTGGCCAATGCAGATCTGACTGAAGCCAAAG
CAGCCCTGACAGCAGCTGGTGTGACTGGCACTGCAAGTGTGGTGAAAATGAGCTATACTGATAACAATGGCAAAACCATT
GATGGTGGCCTGGCAGTGAAAGTGGGTGATGATTATTATAGTGCCACTCAGAATAAAGATGGCAGCATTAGCATTAACAC
CACCAAATATACAGCAGATGATGGCACTAGCAAAACTGCCCTGAACAAGCTGGGTGGTGCAGATGGCAAAACTGAAGTGG
TGAGCATTGGTGGCAAAACCTATGCAGCAAGCAAAGCAGAAGGCCATAACTTTAAAGCACAGCCTGATCTGGCAGAAGCA
GCAGCCACCACCACTGAAAACCCATTACAGAAGATTGATGCAGCCCTGGCCCAAGTGGATACCCTGAGAAGTGATCTGGG
TGCAGTGCAGAACAGATTTAACAGTGCCATCACCAACCTGGGCAACACTGTGAACAACCTGACCTCTGCAAGAAGCAGAA
TTGAAGATAGTGATTATGCCACTGAAGTGAGCAACATGAGTAGAGCACAGATTTTACAACAAGCTGGCACAAGTGTGCTG
GCCCAAGCCAATCAAGTGCCACAGAATGTGTTGTCTCTGCTGAGAGGTGGTCATCATCATCATCATCATTAATAA
```

**Supplementary Table S14. MS2 gpL lysis-protein sequence (5'-3'; 228 bp).** Feature key: MS2 gpL lysis protein

```
ATGGAgACCCGATTCCCTCAGCAATCGCAGCAAACTCCGGCATCTACcAAcAGACGCCGGCCATTCAAgCATGAGGATTA
CCCATGTCGAAGACAACAAAGAAGTTCAACTCTTTATGTATTGATCTTCCTCGCGATCTTTCTCTCGAAATTTACCAATC
AATTGCTTCTGTCGCTACTGGAAGCGGTGATCCGCACAGTGACGACTTTACAGCAATTGCTTACTTAA
```

**Supplementary Table S15. PhiX174 gpE lysis-protein sequence (5'-3'; 276 bp).** Feature key: PhiX174 gpE lysis protein

```
ATGGTACGCTGGACTTTGTGGGATACCCTCGCTTTCCTcCTgtTGctcAGTTTATTGCTGCCGTCATTGCTgATcATGTT
CATCCCGTCAACATTCAAACGGCCTGTCTCATCATGGAAGGCGCTGAATTTACGGAAAACActgTTAATGGCGTCGAGCG
TCCGGctgAAGCCGCTGAATTGTTCGCGTTTACCTTGCGTGTACGCGCAGGAAACACTGACGTTCTTACTGACGCAGAAG
AAAACGTGCGTCAAAAATTACGTGCGGAAGGAGTGA
```

**Supplementary Table S16. Lambda LysS lysis-protein sequence (5'-3'; 324 bp).** Feature key: Lambda LysS lysis protein

```
ATGAAGATGCCAGAAAAACATGACCTGTTGGCCGCCATTCTCGCGGCAAAGGAACAAGGCATCGGGGCAATCCTTGCGTT
TGCAATGGCGTACCTTCGCGGCAGATATAATGGCGGTGCGTTTACAAAAACAGTAATCGACGCAACGATGTGCGCCATTA
TCGCCTGGTTCATTCGTGACCTTCTCGACTTCGCCGGACTAAGTAGCAATCTCGCTTATATAACGAGCGTGTTTATCGGC
TACATCGGTACTGACTCGATTGGTTCGCTTATCAAACGCTTCGCTGCTAAAAAAGCCGGAGTAGAAGATGGTAGAAATCA
ATAA
```

**Supplementary Table S17. Lambda LysR lysis-protein sequence (5'-3'; 477 bp).** Feature key: Lambda LysR lysis protein

```
ATGGTAGAAATCAATAATCAACGTAAGGCGTTCCTCGATATGCTGGCGTGGTCGGAGGGAACTGATAACGGACGTCAGAA
AACCAGAAATCATGGTTATGACGTCATTGTAGGCGGAGAGCTATTTACTGATTACTCCGATCACCCTCGCAAACTTGTCA
CGCTAAACCCAAAACTCAAATCAACAGGCGCCGGACGCTACCAGCTTCTTTCCCGTTGGTGGGATGCCTACCGCAAGCAG
CTTGGCCTGAAAGACTTCTCTCCGAAAAGTCAGGACGCTGTGGCATTGCAGCAGATTAAGGAGCGTGGCGCTTTACCTAT
GATTGATCGTGGTGATATCCGTCAGGCAATCGACCGTTGCAGCAATATCTGGGCTTCACTGCCGGGCGCTGGTTATGGTC
AGTTCGAGCATAAGGCTGACAGCCTGATTGCAAAATTCAAAGAAGCGGGCGGAACGGTCAGAGAGATTGATGTATGA
```

**Supplementary Table S18. Lambda Rz lysis-protein sequence (5'-3'; 461 bp).** Feature key: Lambda Rz lysis protein

```
ATGAGCAGAGTCACCGCGATTATCTCCGCTCTGGTTATCTGCATCATCGTCTGCCTGTCATGGGCTGTTAATCATTACCG
TGATAACGCCATTACCTACAAAGCCCAGCGCGACAAAAATGCCAGAGAACTGAAGCTGGCGAACGCGGCAATTACTGACA
TGCAGATGCGTCAGCGTGATGTTGCTGCGCTCGATGCAAAATACACGAAGGAGTTAGCTGATGCTAAAGCTGAAAATGAT
GCTCTGCGTGATGATGTTGCCGCTGGTCGTCGTCGGTTGCACATCAAAGCAGTCTGTCAGTCAGTGCGTGAAGCCACCAC
CGCCTCCGGCGTGGATAATGCAGCCTCCCCCCGACTGGCAGACACCGCTGAACGGGATTATTTCACCCTCAGAGAGAGGC
TGATCACTATGCAAAAACAACTGGAAGGAACCCAGAAGTATATTAATGAGCAGTGCAGATA
```

**Supplementary Table S19. Full plasmid sequence from the source map for pP4-DiffJJ-Test (pAJ1046; 11,627 bp).** Source file: pAJ1046_DiffTestP4Cosmid_Assembly_warwick-caja3-c1_willscott.dna. Feature key: cos packaging region; P4 gop promoter; EGFP fragment; L3S2P21 terminator; PDiffJJ promoter; MS2 gpL lysis protein; PhiX174 gpE lysis protein; Lambda LysS lysis protein; Lambda LysR lysis protein; Lambda Rz lysis protein; T7 terminator; lambda t0 terminator; kanamycin-resistance promoter; kanamycin resistance gene; P4 replication origin; P4 crr replication region; P4 alpha terminator; P4 alpha; P4 epsilon; P4 cI; P4 pLE promoter; P4 pLL promoter; P4 *sid* promoter; P4 *sid*; P4 delta; P4 psu; P4 tsid terminator

```
GGCGAGGCGGGGAAAGCACCGCGCGCAAACCGACAAGTTAGTTAATTATTTGTGTAGTCAAAGTGCCTTCAGTACATACC
TCGTTAATACATTGGAGCATAATGAAGAAAATCTATGGCCTATGGTCCAAAACTGTCTTTTTTGATGGCACTATCCTGAA
AAATATGCAAAAAATAGATTGATGTAAGGTGGTTCTTGTCAGTGTCGCAAGATCCTTAAGAATTCGTGGCATGAGAGAGT
TAAAGGCTGAAGTTCATCTGCACCACCGGCAAGCTGCCCGTGCCCTGGCCCACCCTCGTGACCACCCTGACCTACGGCGT
GCAGTGCTTCAGCCGCTACCCCGACCACATGAAGCAGCACGACTTCTTCAAGTCCGCCATGCCCGAAGGCTACGTCCAGG
AGCGCACCATCTTCTTCAAGGACGACGGCAACTACAAGACCCGCGCCGAGGTGAAGTTCGAGGGCGACACCCTGGTGAAC
CGCATCGAGCTGAAGGGCATCGACTTCAAGGAGGACGGCAACATCCTGGGGCACAAGCTGGAGTACAACTACAACAGCCA
CAACGTCTATATCATGGCCGACAAGCAGAAGAACGGCATCAAGGTGAACTTCAAGATCCGCCACAACATCGAGGACGGCA
GCGTGCAGCTCGCCGACCACTACCAGCAGAACACCCCCATCGGCGACGGCCCCGTGCTGCTGCCCGACAACCACTACCTG
AGCACCCAGTCCGCCCTGAGCAAAGACCCCAACGAGAAGCGCGATCACATGGTCCTACTGGAGTTCGTGACCGCCGCCGG
GATCACTCTCGGCATGGACGAGCTGTACAAGTAACCCGGGCTCGGTACCAAATTCCAGAAAAGAGGCCTCCCGAAAGGGG
GGCCTTTTTTCGTTTTGGTCCGAGCTCCAATTCCGACGTCTAAGAAACCATTATTATCATGACATTAACCTATAAAAATA
GGCGTATCACGAGGCCCTTTCGTCTTCACCTCGAGTCCCTATCAGTGATAGAGATTGACCTCCCTATCAGTGATAGAGAT
ACTGAGCACATCAGCAGGACGCACTGACCCCATGGTCTAGAGAAAGAGGAGAAAGGAGATATGGAGACCCGATTCCCTCA
GCAATCGCAGCAAACTCCGGCATCTACCAACAGACGCCGGCCATTCAAGCATGAGGATTACCCATGTCGAAGACAACAAA
GAAGTTCAACTCTTTATGTATTGATCTTCCTCGCGATCTTTCTCTCGAAATTTACCAATCAATTGCTTCTGTCGCTACTG
GAAGCGGTGATCCGCACAGTGACGACTTTACAGCAATTGCTTACTTAAGGTAAAGAGGGGAAAGGAGATATGGTACGCTG
GACTTTGTGGGATACCCTCGCTTTCCTCCTGTTGCTCAGTTTATTGCTGCCGTCATTGCTGATCATGTTCATCCCGTCAA
CATTCAAACGGCCTGTCTCATCATGGAAGGCGCTGAATTTACGGAAAACACTGTTAATGGCGTCGAGCGTCCGGCTGAAG
CCGCTGAATTGTTCGCGTTTACCTTGCGTGTACGCGCAGGAAACACTGACGTTCTTACTGACGCAGAAGAAAACGTGCGT
CAAAAATTACGTGCGGAAGGAGTGATGTAATGAAGATGCCAGAAAAACATGACCTGTTGGCCGCCATTCTCGCGGCAAAG
GAACAAGGCATCGGGGCAATCCTTGCGTTTGCAATGGCGTACCTTCGCGGCAGATATAATGGCGGTGCGTTTACAAAAAC
AGTAATCGACGCAACGATGTGCGCCATTATCGCCTGGTTCATTCGTGACCTTCTCGACTTCGCCGGACTAAGTAGCAATC
TCGCTTATATAACGAGCGTGTTTATCGGCTACATCGGTACTGACTCGATTGGTTCGCTTATCAAACGCTTCGCTGCTAAA
AAAGCCGGAGTAGAAGATGGTAGAAATCAATAATCAACGTAAGGCGTTCCTCGATATGCTGGCGTGGTCGGAGGGAACTG
ATAACGGACGTCAGAAAACCAGAAATCATGGTTATGACGTCATTGTAGGCGGAGAGCTATTTACTGATTACTCCGATCAC
CCTCGCAAACTTGTCACGCTAAACCCAAAACTCAAATCAACAGGCGCCGGACGCTACCAGCTTCTTTCCCGTTGGTGGGA
TGCCTACCGCAAGCAGCTTGGCCTGAAAGACTTCTCTCCGAAAAGTCAGGACGCTGTGGCATTGCAGCAGATTAAGGAGC
GTGGCGCTTTACCTATGATTGATCGTGGTGATATCCGTCAGGCAATCGACCGTTGCAGCAATATCTGGGCTTCACTGCCG
GGCGCTGGTTATGGTCAGTTCGAGCATAAGGCTGACAGCCTGATTGCAAAATTCAAAGAAGCGGGCGGAACGGTCAGAGA
GATTGATGTATGAGCAGAGTCACCGCGATTATCTCCGCTCTGGTTATCTGCATCATCGTCTGCCTGTCATGGGCTGTTAA
TCATTACCGTGATAACGCCATTACCTACAAAGCCCAGCGCGACAAAAATGCCAGAGAACTGAAGCTGGCGAACGCGGCAA
TTACTGACATGCAGATGCGTCAGCGTGATGTTGCTGCGCTCGATGCAAAATACACGAAGGAGTTAGCTGATGCTAAAGCT
GAAAATGATGCTCTGCGTGATGATGTTGCCGCTGGTCGTCGTCGGTTGCACATCAAAGCAGTCTGTCAGTCAGTGCGTGA
```

AGCCACCACCGCCTCCGGCGTGGATAATGCAGCCTCCCCCCGACTGGCAGACACCGCTGAACGGGATTATTTCACCCTCA
GAGAGAGGCTGATCACTATGCAAAAACAACTGGAAGGAACCCAGAAGTATATTAATGAGCAGTGCAGATAACTAGCATAA
CCCCTTGGGGCCTCTAAACGGGTCTTGAGGGGTTTTTTGAGACAAACAAAAGAATGGAATCAAAGTTAATGCTAGCAGCC
GCCGCTGCAGGCATGCAAGCTTGCGGCCGCGTCGTGACTGGGAAAACCCTGGCGACTAGTCTTGGACTCCTGTTGATAGA
TCCAGTAATGACCTCAGAACTCCATCTGGATTTGTTCAGAACGCTCGGTTGCCGCCGGGCGTTTTTTATTGGTGAGAATC
CAGGGGTCCCCAATAATTACGATTTAAATTTGTGTCTCAAAATCTCTGATGTTACATTGCACAAGATAAAAATATATCAT
CATGAACAATAAAACTGTCTGCTTACATAAACAGTAATACAAGGGGTGTTATGAGCCATATTCAGCGTGAAACGAGCTGT
AGCCGTCCGCGTCTGAACAGCAACATGGATGCGGATCTGTATGGCTATAAATGGGCGCGTGATAACGTGGGTCAGAGCGG
CGCGACCATTTATCGTCTGTATGGCAAACCGGATGCGCCGGAACTGTTTCTGAAACATGGCAAAGGCAGCGTGGCGAACG
ATGTGACCGATGAAATGGTGCGTCTGAACTGGCTGACCGAATTTATGCCGCTGCCGACCATTAAACATTTTATTCGCACC
CCGGATGATGCGTGGCTGCTGACCACCGCGATTCCGGGCAAAACCGCGTTTCAGGTGCTGGAAGAATATCCGGATAGCGG
CGAAAACATTGTGGATGCGCTGGCCGTGTTTCTGCGTCGTCTGCATAGCATTCCGGTGTGCAACTGCCCGTTTAACAGCG
ATCGTGTGTTTCGTCTGGCCCAGGCGCAGAGCCGTATGAACAACGGCCTGGTGGATGCGAGCGATTTTGATGATGAACGT
AACGGCTGGCCGGTGGAACAGGTGTGGAAAGAAATGCATAAACTGCTGCCGTTTAGCCCGGATAGCGTGGTGACCCACGG
CGATTTTAGCCTGGATAACCTGATTTTCGATGAAGGCAAACTGATTGGCTGCATTGATGTGGGCCGTGTGGGCATTGCGG
ATCGTTATCAGGATCTGGCCATTCTGTGGAACTGCCTGGGCGAATTTAGCCCGAGCCTGCAAAAACGTCTGTTTCAGAAA
TATGGCATTGATAATCCGGATATGAACAAACTGCAATTTCATCTGATGCTGGATGAATTTTTCTAACTTGGAAGTAAGAA
TGGTGCCGAAGGCCGGACTCAAACATCAAAATAAGTTAATGATAAAAAACAAATAATAAAACACAACAATGAAATATGCC
CCCTTTTGTGCCCCCACTGTTTTTCTGACCAATCTATTTTCAGCCCATCAATAAATCGGAAAGTTAAATCATTTTTAATC
AGTAAGTTTGGATCCGTAGCTCGGATCCAAACCAGTGCATCTTTTATCCACATAAAAAATTTTTTTTCGAAAGAACTGTT
CACACTGTTCACCTTTCTGTTTTCTCCTTTTATTTCAGAGTGATAGGTGGTGAATAATGGGTGAAGGGTGAACATTCGAT
TCTTCACCTCCGGCATTCTGCCGATGTGACTCATACCGGTGATTAATCCTCCGCACTGAAATCACTCAGGAAGAAAAAAG
TTTTTTTTGATTTGATTGTTCACACTGTTCACCTTTCGTTTTTCTCTTTTAATTTCAGTGTGATAACGGGTGAATATACG
GTGAAGGGTGAACAGTGGATTGTTCACCTTCGGGGGATATCGGGATAAAAAAAGACCGGCAGATGCCGGTCAGGTGGGTC
AGGCTGTTGTAGGGTCGTCACATTTTGGCAGCCAGTCGCCGTAGCTTTCCTCTTTCAGCGTCAGGTTGGTCTGTATCCCC
TGTTTGGTATGGCGTTTCTCGTAATTCAGTCCGTATTCCTTCAGCATCACCGGCAGCCCCAGCCCGAACATTTTCAGACT
GAGTACATTCCGGTAGCCGTTTGCCTCCATGTAGGCCAGATAGGCGTGATAGAGGTATTTACGGTAATTGCGCGGGATGA
TACTGGCGTTCCCCATATACATGCCGCTGGTCTGCGGCAGGGTTTCCAGATAGCCGATAAAATCAAACGTCGGGTCGGCA
TCCCGTTTGATGTTCAGTGCCTCGTCTGAGTTCTGCTGGGACTGAAGCAGTGACCGGGCGAGCATCGGGTCGCTGAACTT
CTGCATCAGGTGACGCACGATGACCGCCAGCTCGCGGGTGATTTTGTCCTTAAGCTGCGGGTCGCGCTCCTGCGGGGCTA
TCTGTTCCGGGAAGTGAATAATCACCCGTCGGCGTGACACGCCGCCGCTGCGGTCGGTGAAGCGCATCGGGTTATTGTTC
ACGGCCAGAATCACCGCCGGGATGTGCGTGGAGTACGCATCCCGGTATTTCGGGTCAACGGACACCGCATCGCCGCCGGT
GATGGCCTTGAGTCCGGCACCGTCGCCGCTCCATTTTTCCTGGTCCGGCAGGCGTATCAGTGAGAAGCCAGTTAACGCGG
CACGTTCACGCGGGGATTCCAGCGTCTCGATGGTGGCCGACGTGGCGTTATCCTCCCCGGCCAGCAGGGTGGCTATTTCG
GCCATGATACTTTTGCCGCTGCCGCCGGGACCGGTCACCTCCAGAAAGAGCTGCCAGTCGTAGCGGTTTGCCAGCACCAT
AAACAGTGCAGCCAGAATCACGTCGCGTTTTTCCGCACGGCCACCGGCGGCACGGTCAAGCCAGCGCCAGAAGGCGGGGG
CGTGGGTTTCCAGCGTTTCACCGTCCACCGGCGGGGTGAAATCCACATCGCACAGGGTGCGCATCCAGTGTGACGGACTG
TGCGGGTGGAACGTGCCGTTCTGCGTGTCGAGCACGCCGTTACGAAAGCCAATCAGGCGGCGGGAGGGGGCTTCCTGCTG
CGGAATAATCAGCTTCAGGGTGTCCACCACGGAGGCCACCTTCCCGGAGGAGAACGGCGCACGCAGACGCTGAAACAGCC
CGGCCACATCCCGGGCAAAGTCCTGTGGCGGCAGCACCTTCCAGACACCATTTTCATAGCGGGACAGAAGCTGGCCGTTG
GCATCGACCGCGAGCGCCTCGCCGTAATGCTCATAGATACGCATGGCCTTTTCGCTGGTACTCATGGCGGAAAACTCCGC
TTCGCTCATGGTGTCGAACGGGCTTTCAGCCGGTGGCCGGATGGCATCGTAAATGGCCTTACGGGTGGCCTCCCCGCCGT
ACTGCGTGAAGGCATCATTCCAGTCACCGAAGACCGGCGGCAGGGCAACAACACCTTCACACGCATCTGCGGCTGCGGCG
GCTTTTTTCTGGCCGTCACCACTGAGGTCACGGTCAGCGGCAAGGACAATCTGACAGGCGGGGTGCTTCTGCCGGGCAAG
GCTGGCCAGAGAAAGGAGGTTCACGGAAGAAAGCGCCACCATCACCGTTTCACCGGTCAGGTGATGTACGGTAAGTGCGG
TCGCGTATCCCTCCGCTATCCACAGACGTTTTCCGGCCTGATTCTGTCCTTCAAGGGTGTGACAGGTGCCCCTGACCTGT
CCGCCTTTCAGGGTGCGCTTACGGCCGTCAGCACTGATTAACTGAAGGTTAACCAGTTCGCCGCTGTCGTCATACAGTGG
CACCACAAGGTCACCGGCGCGCCAGCTCACGCCACCGGCTCTGTGTGTGCCGGTCAGCATCCGGCATTCCCGGCCGGGAA
AGCCCTTGCGGGTCAGGTAGGCGTTACCGGTTCCGGTACGGGTTTTCGCCATCAGGGTTTGTGCCAGTGCGGCGGCGTTC
TTCCGGGCAGCGTCTGTTTCATCAACGGCGGCGGTCGTCACTGCCGGGTCAGCCGGTGGCAGGCTGCCGGTCACGGCAGC
CACCTTTGCGGCCGCGTCGGACGGGGAAACACCAAAAACCTTTTCAACCAGTTTCAGGCCGTCACCGGCACCACACTGAT
TGCAGTACCAGGTGCCGCGCCCCTCCCTGTCATCAAAACGGAAGCGGTCACTCCCGCCACAGACCGGACAGGGCTGATGA
CGGTTCTTCAGCACCTGAATCCCCAGCGCCGGGAGAATACGCGGCCAGTGGCCGAGCGCATGGCTGACGGTGGCGGTTAC
GTTCATTTTCATGGTGTTGTTCTCCTTCAGTGCAGTACCGGCGCTTTTATGTGACGGGCACAGAGTTCATCCATCACAAC
CAGCCCGAGAAAGGACAGCGACGGCGCGGCCTTCAGGGGGCCGGATTCCATTAAATCTTCCAGCAGGGCACAGGCTATCT
GACGCCCTTTTTCCTCACCGTGCTGGCGCAGATAAAAGCCTTCCAGCTCAGCGGCGATGGCCGCCTCCAGTGACTCAAGG
GTGAGATGCGGGTAGCGGTGCTGACGTTCGCACACGGTCAGCCAGGCACAGGCGACAGCGCGACGGTAAAGGGCAGCGCG
TAAGACGGGCGGTAAGGGTGTTTTCATTTGCTTTTCTCCCTGTGACAGATGACTGCATTCCGTGCCGGTTGCATTAACTG
ATAAGGCATATCTGCGTCTCCTGAAGACGTGCGTATCCCTGCGCGAATACGCACATTTAATTTTTCGGGGGTCGTTTTTT
AATTACAGATAATTGCGGTAACTGTTATCCGGGGTGGTTTCCGGGTCAGGCTCCGTGCGGGGAATTTCCCGCCATTCCCG
CGCCACCGGTGCTGCCCGGCTGACCGGAACAGTGTCCTGCGGGTAAATATCCAGATATTTTTCCCGCCATTTCTGTAATT
CCGGGTCTCCGGCCATTTCTTTCAGTACCGCATGCCGGTTTACGGGGCTGCGTTTAAACAGGTCAGGACGGTCACAGGTA
AATTCCCGCAGAAAACGCCCCAGCGGGATGTCTGTGGTGCGTCCGTCAGCGAGGATACGCACAAGGATACTGAATTTACG
GCGGTACGGGTTCCAGACAATGTCCGGGCAGCGGTACGGCATTTCCCACGGAATACCGTCTTCCAGAATGCCGACCACGG
CCACATCGGGAAAACCGGCAGAACGGTAAATCTCACCGGGCTGGGGAAAATCAAACATGCGTCCTGTCTCCCCGGTCTTT
CTGCTGGGCGAGAAAATCGCGGCACAGGCCTTTGGCTTTCAGCTCATTCAGCACAAAATCAATATCTTCATTCAGGTAGC
TGAAAATATGTGGAATGTAGAGCTGATGCAGGCCGGAGAGTTCACGGTGAATCAAATCACCCCCAACAAACCGGGATACG
GCGCTGGCGCGGTTGAGCTTATGGTAAGCCTCAATGCTGAGGTGTTCACGGGCGTCATGACGCGCTGAGACGGTCTGAGG
GGCTTTTTTATTACGCACGGGACACCTCCACCACCGGCAGACGGGCAGCAAGGGAGAGCACATAGTCACGGACAAGGGAA
CGGCGGGCACTGCGTTCATCACCGGCGACGGTGCGAAGCATACAGATACGGGGATGACGGTCTGCGCGACGGACAGCCGC
AAACACAAAGACAAATTCAGGGTGTGAGGGGGTAAGGGTTGTAGCCATGATGGCAGCCTCCTGTGAATAGCAAATAACGC
TATCGCCGGAGTTCTCACGCTCGATGGCGATAGCCCAGACGGGGGTGAGAATACCGGCTTCACAGGATACCGGCCAGCCC
GGAGGCTGCCCCGCCTGAGCTACCATTGACTCTGCGGCATAATGAGCGGACGCGGGCAGGATGCACGGAATGCCATCTGC
ACGACTGACCACACACCACACCATAATCTGGCGCTCTGTGGCATTGATTGCGACACAAAAAAAGACGCGTGGCGCGTCAT
ATGTCGCCTGTGAATTGCTCGGGTTCTCACGCCCGGCTGCCGATTTTGCGGCAGCGGAAAAACTATATCCGCAAATGCCG
GAAAAAGGCAAGCCAGAAAAAGGGAGTTTTTGCAGAGCGGGCATCATCATGCGTCGTACCCCCGTTTGCGTCCGGCAATG
CGTCCGGCCATCCATGCGGTGACTTCAGAGTGCAGCCAGGCCACATTTTTACCGCCAAGACTCACCTGCGGCGGAAATTC
CCCCTTACGGATGAGTTCGTAGATGGTCGAGCGTGACAGGCCGCACAGGTGCATCACTTCCGGCAGACGTAAAAAACGCT
CCTGCGTGATGTCCGGCAGCGGCATCAGTGGCGTCACTGGGGCGGGAGACGGGGAAGAAAAAACAGCTTGCATCGGGCTA
CCTCGTTAATGTCCATACAGCACCGGATAAGTCCGTCCGGCTTCGGGTAGCGCTTTATTTTATGAATATTTTCAGCAGAC
GCAACAGGGGGGATTTGTTCAGGCTGTCTTACAATGGCTGTGTGTTTTTTGTTCATCTCCACTTAAAGTCATTTAAAGCC
ACTTAAAGCAATTTGTAATTTTTATAGTGAAATACAAATCGTTTCTTCTTATTCATTCCCGGCGAATTAATAAAAACAAA
CAGTAGTAAACAGCACAAAAAGCCCATCAACGGGTGAACAGTGGTGAACAGACGGTGAACAGTCATTACTGCGATTGTTC
ACCCTTTAACTTACTGTATTACTTATCTTTTTTATTAAGGTGAACAGAGGTGAACAGTAAAATATAAAAAAACAAACAGT

```
AAGCCGGTTTTTCCTGCGACCTTTTCCTGGCTTGCCGGTCTGAGGATGAGTCTCCTGTGTCAGGGCTGGCACATCTGCAA
TGCGTCGTGTTGTTGTCCGGTGTACGTCACAATTTTCTTAACCTGAAGTGACGAGGAGCCGGAAAATGTCTGACCACACT
ATCCCTGAATATCTGCAACCCGCACTGGCACAACTGGAAAAGGCCAGAGCCGCCCATCTTGAGAACGCCCGCCTGATGGA
TGAGACCGTCACGGCCATTGAACGGGCAGAGCAGGAAAAAAATGCGCTGGCGCAGGCCGACGGAAACGACGCTGACGACT
GGCGCACGGCCTTTCGTGCAGCCGGTGGTGTCCTGAGCGACGAGCTGAAACAGCGCCACATTGAGCGCGTGGCACGCCGG
GAGCTGGTACAGGAATATGACAATCTGGCCGTGGTGCTGAATTTCGAACGTGAACGCCTGAAAGGGGCGTGTGACAGCAC
GGCCACCGCCTACCGGAAGGCACATCATCACCTTCTGAGTCTGTATGCAGAGCATGAGCTGGAACACGCCCTGAATGAAA
CCTGTGAGGCGCTTGTCCGGGCAATGCATCTGAGCATTCTGGTACAGGAAAATCCGCTCGCCAACACCACCGGCCATCAG
GGCTACGTCGCACCGGAAAAGGCTGTCATGCAGCAGGTGAAATCATCGCTGGAACAGAAAATTAAACAGATGCAAATCAG
CCTCACCGGCGAGCCGGTTCTCCGGCTGACCGGACTGTCAGCGGCAACACTCCCGCACATGGATTATGAGGTGGCAGGCA
CACCGGCACAGCGCAAGGTGTGGCAGGACAAAATAGACCAGCAGGGAGCAGAGCTTAAGGCCAGAGGGCTGCTGTCATGA
TTTACTGTCCGTCGTGTGGACATGTTGCTCACACCCGTCGCGCACATTTCATGGACGATGGCACCAAGATAATGATTGCA
CAGTGCCGGAATATTTATTGCTCTGCGACATTTGAAGCGAGTGAAAGCTTTTTCTCTGACAGTAAAGATTCAGGAATGGA
ATACATTTCAGGCAAACAGAGATACCGCGATTCACTGACGTCAGCCTCCTGCGGTATGAAACGCCCGAAAAGAATGCTTG
TTACCGGATATTGTTGTCGGAGATGTAAAGGCCTTGCACTGTCAAGAACATCGCGGCGTCTGTCTCAGGAAGTCACCGAG
CGTTTTTATGTGTGCACGGATCCGGGCTGTGGTCTGGTGTTTAAAACGCTTCAGACCATCAACCGCTTCATTGTCCGCCC
GGTCACGCCGGACGAACTGGCAGAACGCCTGCATGAAAAACAGGAACTGCCGCCAGTACGGTTAAAAACACAATCATATT
CGCTGCGTCTGGAATGAGGGCTGCCGGTTAACACCGGCCGTCGCCGCACACCGTATTTTTATTCTTCAGCATGATGAGAA
AGAGATAACGATGGAAAGCACAGCCTTACAGCAGGCCTTTGACACCTGTCAGAATAACAAAGCAGCATGGCTGCAACGCA
AAAATGAGCTGGCAGCGGCCGAACAGGAATATCTGCGGCTTCTGTCAGGAGAAGGCAGAAACGTCAGTCGCCTGGACGAA
TTACGCAATATTATCGAAGTCAGAAAATGGCAGGTGAATCAGGCCGCCGGTCGTTATATTCGTTCGCATGAAGCCGTTCA
GCACATCAGCATCCGCGACCGGCTGAATGATTTTATGCAGCAGCACGGCACAGCACTGGCGGCCGCACTGGCACCGGAGC
TGATGGGCTACAGTGAGCTGACGGCCATTGCCCGAAACTGTGCCATACAGCGTGCCACAGATGCCCTGCGTGAAGCCCTT
CTGTCCTGGCTTGCGAAGGGTGAAAAAATTAATTATTCCGCACAGGATAGCGACATTTTAACGACCATCGGATTCAGGCC
TGACGTGGCTTCGGTGGATGACAGCCGTGAAAAATTCACCCCTGCGCAGAACATGATTTTTTCGCGTAAAAGTGCGCAAC
TGGCATCACGTCAGTCAGTGTAAAATTCCCCGAAAATCCGCCCGTTTTTACTGAAAAAAGCCATGCATCGATAAGGTGCA
TGGCTTTGCATGCGTTTTCCTGCCTCATTTTCTGCAAACCGCGCCATTCCCGGCGCGGTCTGAGCGTGTCAGTGCAACTG
CATTAAAACCGCCCCGCAAAGCGGGCG
```

**Supplementary Table S20. Full plasmid sequence from the source map for pP4-DiffJJ-LacIq (pAJ1053; 6,850 bp).** Source file: pAJ1053_DiffP4_FullAssembly_warwick-caja2-b8.dna. Feature key: Lambda LysR lysis protein; Lambda LysS lysis protein; MS2 gpL lysis protein; PLtetO-1 promoter; L3S2P21 terminator; EGFP fragment; PE antigen; PlacIq promoter; rrnB T1 terminator; M13 origin; RSF/p15A origin region; oriT transfer region; kanamycin resistance gene; kanamycin-resistance promoter; lambda t0 terminator; T7 terminator; Lambda Rz lysis protein

```
TGCGGTAGGCATCCCACCAACGGGAAAGAAGCTGGTAGCGTCCGGCGCCTGTTGATTTGAGTTTTGGGTTTAGCGTGACA
AGTTTGCGAGGGTGATCGGAGTAATCAGTAAATAGCTCTCCGCCTACAATGACGTCATAACCATGATTTCTGGTTTTCTG
ACGTCCGTTATCAGTTCCCTCCGACCACGCCAGCATATCGAGGAACGCCTTACGTTGATTATTGATTTCTACCATCTTCT
ACTCCGGCTTTTTTAGCAGCGAAGCGTTTGATAAGCGAACCAATCGAGTCAGTACCGATGTAGCCGATAAACACGCTCGT
TATATAAGCGAGATTGCTACTTAGTCCGGCGAAGTCGAGAAGGTCACGAATGAACCAGGCGATAATGGCGCACATCGTTG
CGTCGATTACTGTTTTTGTAAACGCACCGCCATTATATCTGCCGCGAAGGTACGCCATTGCAAACGCAAGGATTGCCCCG
ATGCCTTGTTCCTTTGCCGCGAGAATGGCGGCCAACAGGTCATGTTTTTCTGGCATCTTCATTACATCACTCCTTCCGCA
CGTAATTTTTGACGCACGTTTTCTTCTGCGTCAGTAAGAACGTCAGTGTTTCCTGCGCGTACACGCAAGGTAAACGCGAA
CAATTCAGCGGCTTCAGCCGGACGCTCGACGCCATTAACAGTGTTTTCCGTAAATTCAGCGCCTTCCATGATGAGACAGG
CCGTTTGAATGTTGACGGGATGAACATGATCAGCAATGACGGCAGCAATAAACTGAGCAACAGGAGGAAAGCGAGGGTAT
CCCACAAAGTCCAGCGTACCATATCTCCTTTCCCCTCTTTACCTTAAGTAAGCAATTGCTGTAAAGTCGTCACTGTGCGG
ATCACCGCTTCCAGTAGCGACAGAAGCAATTGATTGGTAAATTTCGAGAGAAAGATCGCGAGGAAGATCAATACATAAAG
AGTTGAACTTCTTTGTTGTCTTCGACATGGGTAATCCTCATGCTTGAATGGCCGGCGTCTGTTGGTAGATGCCGGAGTTT
GCTGCGATTGCTGAGGGAATCGGGTCTCCATATCTCCTTTCTCCTCTTTCTCTAGACCATGGGGTCAGTGCGTCCTGCTG
ATGTGCTCAGTATCTCTATCACTGATAGGGATGTCAATCTCTATCACTGATAGGGACTCGAGGTGAAGACGAAAGGGCCT
CGTGATACGCCTATTTTTATAGGTTAATGTCATGATAATAATGGTTTCTTAGACGTCGGAATTGGAGCTCGGACCAAAAC
GAAAAAAGGCCCCCCTTTCGGGAGGCCTCTTTTCTGGAATTTGGTACCGAGCCCGGGTTACTTGTACAGCTCGTCCATGC
CGAGAGTGATCCCGGCGGCGGTCACGAACTCCAGTAGGACCATGTGATCGCGCTTCTCGTTGGGGTCTTTGCTCAGGGCG
GACTGGGTGCTCAGGTAGTGGTTGTCGGGCAGCAGCACGGGGCCGTCGCCGATGGGGGTGTTCTGCTGGTAGTGGTCGGC
GAGCTGCACGCTGCCGTCCTCGATGTTGTGGCGGATCTTGAAGTTCACCTTGATGCCGTTCTTCTGCTTGTCGGCCATGA
TATAGACGTTGTGGCTGTTGTAGTTGTACTCCAGCTTGTGCCCCAGGATGTTGCCGTCCTCCTTGAAGTCGATGCCCTTC
AGCTCGATGCGGTTCACCAGGGTGTCGCCCTCGAACTTCACCTCGGCGCGGGTCTTGTAGTTGCCGTCGTCCTTGAAGAA
GATGGTGCGCTCCTGGACGTAGCCTTCGGGCATGGCGGACTTGAAGAAGTCGTGCTGCTTCATGTGGTCGGGGTAGCGGC
TGAAGCACTGCACGCCGTAGGTCAGGGTGGTCACGAGGGTGGGCCAGGGCACGGGCAGCTTGCCGGTGGTGCAGATGAAC
TTCAGGGTCAGCTTGCCGTAGGTGGCATCGCCCTCGCCCTCGCCGGACACGCTGAACTTGTGGCCGTTTACGTCGCCGTC
CAGCTCGACCAGGATGGGCACCACCCCGGTGAACAGCTCCTCGCCCTTGCTCACCATATCTGATTTCTCCTCTTTCTCTT
AACGCCGGCGTTATTTTTTATCCACGCTAAAGGCTTCGCCATAGTTAGCGCAAATGATCTGGGCGGCATTGTATAAAGTG
GTATCCGGGGTCAGAGATAAAGTGTGTTTCTTCTGCTTTTTCGGTGCGGCACGCAGACCTTGGCCCCAAAACTCATCGTA
GAAATCGGTGCGAACTTGGGTCAGATGGTAGTTGGCGCAGTTCAGAATTTTATACTGACGAACGCTGCGGGCGTAACGTT
TCGGCTCCGGATACACGTATAAACCTTTGTCTAAATTCACCACGGCATCGAAGTGCACAATCTGCGGTTCTTGATTGTCC
ACCCAGATGCTTTCGCTGTCAATATAGTAATTCACGTTTTTCACTAAACGGATATAACCGCTGCGCACATCGGTCGGAGG
GGCCAGTTTCACATCGTTCTGTTTGGCTTTCTGAATCTGGGCGCTGCATGCGGTCAGTAAACCTAAAGACAGAGTTAAAA
TAATTTTCTTCATATCTCCTTTCTCCTCTTTCGAATTCGGCGCTATCATGCCATACCGCGAAAGGTTTTGCACCATTCGA
GTATTAGGGCCCTCCGGAAATACAGGAACGCACGCTGGATGGCCCTTCGCTGGGATGGTGAAACCATGAAAAATGGCAGC
TTCAGTGGATTAAGTGGGGGTAATGTGGCCTGTACCCTCTGGTTGCATAGGTATTCATACGGTTAAAATTTATCAGGCGC
GATCGCGGCAGTTTTTCGGGTGGTTTGTTGCCATTTTTACCTGTCTGCTGCCGTGATCGCGCTGAACGCGTTTTAGCGGT
GCGTACAATTAAGGGATTATGGTAAATCCACTTACTGTCTGCCCTCGTAGCCATCGATAGACAAACAAAAGAATGGAATC
AAAGTTAATCTTCATTATGCTCCAATGTATTAACGAGGTATGTACTGAAGGCACTTTGACTACACAAATAATTAACTAAC
TTGTCGGTTTGCGCGCGGTGCTTTCCCCGCCTCGCCCGCCCGCTTTGCGGGGCGGTTTTAATGCAGTTGCACTGACACGC
TCAGACCGCGCCGGGAATGGCGCGGTTTGCAGAAAATGAGGCAGGAAAACGCATCCTAGGCGGCCTCCTGTGTGAAATTG
TTATCCGCTTTAATTAAAGGCATCAAATAAAACGAAAGGCTCAGTCGAAAGACTGGGCCTTTCGTTTTATCTGTTGTTTG
TCGGTGAACGCTCTCCTGAGTAGGACAAATCCGCCGCCCTAGACAGCTGGGCGCGCCTCAGTGGAACGAAAACTCACGTT
AAGGGATTTTGGTCATGAGATTATCAAAAAGGATCTTCACCTAGATCCTTTTACGCGCCCTGTAGCGGCGCATTAAGCGC
GGCGGGTGTGGTGGTTACGCGCAGCGTGACCGCTACACTTGCCAGCGCCCTAGCGCCCGCTCCTTTCGCTTTCTTCCCTT
CCTTTCTCGCCACGTTCGCTTTCCCCGTCAAGCTCTAAATCGGGGGCTCCCTTTAGGGTTCCGATTTAGTGCTTTACGGC
```

```
ACCTCGACCCCAAAAAACTTGATTTGGGTGATGGTTCACGTAGTGGGCCATCGCCCTGATAGACGGTTTTTCGCCCTTTG
ACGTTGGAGTCCACGTTCTTTAATAGTGGACTCTTGTTCCAAACTGGAACAACACTCAACCCTATCTCGGGCTATTCTTT
TGATTTATAAGGGATTTTGCCGATTTCGGCCTATTGGTTAAAAAATGAGCTGATTTAACAAAAATTTAACGCGAATTTTA
ACAAAATATTAACGTTTACAATTTAAATATTTGCTTATACAATCTTCCTGTTTTTGGGGCTTTTCTGATTATCAACCGGG
GTAAATCAATCTCGCGCCGGATATATTCCGCTTCCTCGCTCACTGACTCGCTACGCTCGGTCGTTCGACTGCGGCGAGCG
GAAATGGCTTACGAACGGGGCGGAGATTTCCTGGAAGATGCCAGGAAGATACTTAACAGGGAAGTGAGAGGGCCGCGGCA
AAGCCGTTTTTCCATAGGCTCCGCCCCCCTGACAAGCATCACGAAATCTGACGCTCAAATCAGTGGTGGCGAAACCCGAC
AGGACTATAAAGATACCAGGCGTTTCCCCCTGGCGGCTCCCTCGTGCGCTCTCCTGTTCCTGCCTTTCGGTTTACCGGTG
TCATTCCGCTGTTATGGCCGCGTTTGTCTCATTCCACGCCTGACACTCAGTTCCGGGTAGGCAGTTCGCTCCAAGCTGGA
CTGTATGCACGAACCCCCCGTTCAGTCCGACCGCTGCGCCTTATCCGGTAACTATCGTCTTGAGTCCAACCCGGAAAGAC
ATGCAAAAGCACCACTGGCAGCAGCCACTGGTAATTGATTTAGAGGAGTTAGTCTTGAAGTCATGCGCCGGTTAAGGCTA
AACTGAAAGGACAAGTTTTGGTGACTGCGCTCCTCCAAGCCAGTTACCTCGGTTCAAAGAGTTGGTAGCTCAGAGAACCT
TCGAAAAACCGCCCTGCAAGGCGGTTTTTTCGTTTTCAGAGCAAGAGATTACGCGCAGACCAAAACGATCTCAAGAAGAT
CATCTTATTAATCAGATAAAATATTTCTAGGGCCGGCCTACGGCCAGCCTCGCAGAGCAGGATTCCCGTTGAGCACCGCC
AGGTGCGAATAAGGGACAGTGAAGAAGGAACACCCGCTCGCGGGTGGGCCTACTTCACCTATCCTGCCCGGCTGACGCCG
TTGGATACACCAAGGAAAGTCTACACGAACCCTTTGGCAAAATCCTGTATATCGTGCGAAAAAGGATGGATATACCGAAA
AAATCGCTATAATGACCCCGAAGCAGGGTTATGCAGCGGAAAAGGACAACGCGCGGACCGCGGTCCAATTAATTATTAGA
AAAATTCATCCAGCATCAGATGAAATTGCAGTTTGTTCATATCCGGATTATCAATGCCATATTTCTGAAACAGACGTTTT
TGCAGGCTCGGGCTAAATTCGCCCAGGCAGTTCCACAGAATGGCCAGATCCTGATAACGATCCGCAATGCCCACACGGCC
CACATCAATGCAGCCAATCAGTTTGCCTTCATCGAAAATCAGGTTATCCAGGCTAAAATCGCCGTGGGTCACCACGCTAT
CCGGGCTAAACGGCAGCAGTTTATGCATTTCTTTCCACACCTGTTCCACCGGCCAGCCGTTACGTTCATCATCAAAATCG
CTCGCATCCACCAGGCCGTTGTTCATACGGCTCTGCGCCTGGGCCAGACGAAACACACGATCGCTGTTAAACGGGCAGTT
GCACACCGGAATGCTATGCAGACGACGCAGAAACACGGCCAGCGCATCCACAATGTTTTCGCCGCTATCCGGATATTCTT
CCAGCACCTGAAACGCGGTTTTGCCCGGAATCGCGGTGGTCAGCAGCCACGCATCATCCGGGGTGCGAATAAAATGTTTA
ATGGTCGGCAGCGGCATAAATTCGGTCAGCCAGTTCAGACGCACCATTTCATCGGTCACATCGTTCGCCACGCTGCCTTT
GCCATGTTTCAGAAACAGTTCCGGCGCATCCGGTTTGCCATACAGACGATAAATGGTCGCGCCGCTCTGACCCACGTTAT
CACGCGCCCATTTATAGCCATACAGATCCGCATCCATGTTGCTGTTCAGACGCGGACGGCTACAGCTCGTTTCACGCTGA
ATATGGCTCATAACACCCCTTGTATTACTGTTTATGTAAGCAGACAGTTTTATTGTTCATGATGATATATTTTTATCTTG
TGCAATGTAACATCAGAGATTTTGAGACACAAATTTAAATCGTAATTATTGGGGACCCCTGGATTCTCACCAATAAAAAA
CGCCCGGCGGCAACCGAGCGTTCTGAACAAATCCAGATGGAGTTCTGAGGTCATTACTGGATCTATCAACAGGAGTCCAA
GACTAGTCGCCAGGGTTTTCCCAGTCACGACGCGGCCGCAAGCTTGCATGCCTGCAGCGGCGGCTGCTAGCATTAACTTT
GATTCCATTCTTTTGTTTGTCTCAAAAAACCCCTCAAGACCCGTTTAGAGGCCCCAAGGGGTTATGCTAGTTATCTGCAC
TGCTCATTAATATACTTCTGGGTTCCTTCCAGTTGTTTTTGCATAGTGATCAGCCTCTCTCTGAGGGTGAAATAATCCCG
TTCAGCGGTGTCTGCCAGTCGGGGGGAGGCTGCATTATCCACGCCGGAGGCGGTGGTGGCTTCACGCACTGACTGACAGA
CTGCTTTGATGTGCAACCGACGACGACCAGCGGCAACATCATCACGCAGAGCATCATTTTCAGCTTTAGCATCAGCTAAC
TCCTTCGTGTATTTTGCATCGAGCGCAGCAACATCACGCTGACGCATCTGCATGTCAGTAATTGCCGCGTTCGCCAGCTT
CAGTTCTCTGGCATTTTTGTCGCGCTGGGCTTTGTAGGTAATGGCGTTATCACGGTAATGATTAACAGCCCATGACAGGC
AGACGATGATGCAGATAACCAGAGCGGAGATAATCGCGGTGACTCTGCTCATACATCAATCTCTCTGACCGTTCCGCCCG
CTTCTTTGAATTTTGCAATCAGGCTGTCAGCCTTATGCTCGAACTGACCATAACCAGCGCCCGGCAGTGAAGCCCAGATA
TTGCTGCAACGGTCGATTGCCTGACGGATATCACCACGATCAATCATAGGTAAAGCGCCACGCTCCTTAATCTGCTGCAA
TGCCACAGCGTCCTGACTTTTCGGAGAGAAGTCTTTCAGGCCAAGCTGCT
```

**Supplementary Table S21. Full plasmid sequence from the source map for pP4-∆lys (pAJ1057; 7,405 bp).**
Source file: pAJ1057_pP4_noLys_Assembly_warwick-caja4-f6.dna. Feature key: PE antigen; L3S2P21 terminator; T7 terminator; lambda t0 terminator; kanamycin-resistance promoter; kanamycin resistance gene; P4 replication origin; P4 crr replication region; P4 alpha terminator; P4 alpha; P4 epsilon; P4 cI; P4 pLE promoter; P4 gp11/vis; P4 pLL promoter; P4 *sid* promoter

```
CAGAAGAAACACACTTTATCTCTGACCCCGGATACCACTTTATACAATGCCGCCCAGATCATTTGCGCTAACTATGGCGA
AGCCTTTAGCGTGGATAAAAAATAATAACCCGGGCTCGGTACCAAATTCCAGAAAAGAGGCCTCCCGAAAGGGGGGCCTT
TTTTCGTTTTGGTCCCTAGCATAACCCCTTGGGGCCTCTAAACGGGTCTTGAGGGGTTTTTTGAGACAAACAAAAGAATG
GAATCAAAGTTAATGCTAGCAGCCGCCGCTGCAGGCATGCAAGCTTGCGGCCGCGTCGTGACTGGGAAAACCCTGGCGAC
TAGTCTTGGACTCCTGTTGATAGATCCAGTAATGACCTCAGAACTCCATCTGGATTTGTTCAGAACGCTCGGTTGCCGCC
GGGCGTTTTTTATTGGTGAGAATCCAGGGGTCCCCAATAATTACGATTTAAATTTGTGTCTCAAAATCTCTGATGTTACA
TTGCACAAGATAAAAATATATCATCATGAACAATAAAACTGTCTGCTTACATAAACAGTAATACAAGGGGTGTTATGAGC
CATATTCAGCGTGAAACGAGCTGTAGCCGTCCGCGTCTGAACAGCAACATGGATGCGGATCTGTATGGCTATAAATGGGC
GCGTGATAACGTGGGTCAGAGCGGCGCGACCATTTATCGTCTGTATGGCAAACCGGATGCGCCGGAACTGTTTCTGAAAC
ATGGCAAAGGCAGCGTGGCGAACGATGTGACCGATGAAATGGTGCGTCTGAACTGGCTGACCGAATTTATGCCGCTGCCG
ACCATTAAACATTTTATTCGCACCCCGGATGATGCGTGGCTGCTGACCACCGCGATTCCGGGCAAAACCGCGTTTCAGGT
GCTGGAAGAATATCCGGATAGCGGCGAAAACATTGTGGATGCGCTGGCCGTGTTTCTGCGTCGTCTGCATAGCATTCCGG
TGTGCAACTGCCCGTTTAACAGCGATCGTGTGTTTCGTCTGGCCCAGGCGCAGAGCCGTATGAACAACGGCCTGGTGGAT
GCGAGCGATTTTGATGATGAACGTAACGGCTGGCCGGTGGAACAGGTGTGGAAAGAAATGCATAAACTGCTGCCGTTTAG
CCCGGATAGCGTGGTGACCCACGGCGATTTTAGCCTGGATAACCTGATTTTCGATGAAGGCAAACTGATTGGCTGCATTG
ATGTGGGCCGTGTGGGCATTGCGGATCGTTATCAGGATCTGGCCATTCTGTGGAACTGCCTGGGCGAATTTAGCCCGAGC
CTGCAAAAACGTCTGTTTCAGAAATATGGCATTGATAATCCGGATATGAACAAACTGCAATTTCATCTGATGCTGGATGA
ATTTTTCTAACTTGGAAGTAAGAATGGTGCCGAAGGCCGGACTCAAACATCAAAATAAGTTAATGATAAAAAACAAATAA
TAAAACACAACAATGAAATATGCCCCCTTTTGTGCCCCCACTGTTTTTCTGACCAATCTATTTTCAGCCCATCAATAAAT
CGGAAAGTTAAATCATTTTTAATCAGTAAGTTTGGATCCGTAGCTCGGATCCAAACCAGTGCATCTTTTATCCACATAAA
AAATTTTTTTTCGAAAGAACTGTTCACACTGTTCACCTTTCTGTTTTCTCCTTTTATTTCAGAGTGATAGGTGGTGAATA
ATGGGTGAAGGGTGAACATTCGATTCTTCACCTCCGGCATTCTGCCGATGTGACTCATACCGGTGATTAATCCTCCGCAC
TGAAATCACTCAGGAAGAAAAAAGTTTTTTTTGATTTGATTGTTCACACTGTTCACCTTTCGTTTTTCTCTTTTAATTTC
AGTGTGATAACGGGTGAATATACGGTGAAGGGTGAACAGTGGATTGTTCACCTTCGGGGGATATCGGGATAAAAAAAGAC
CGGCAGATGCCGGTCAGGTGGGTCAGGCTGTTGTAGGGTCGTCACATTTTGGCAGCCAGTCGCCGTAGCTTTCCTCTTTC
AGCGTCAGGTTGGTCTGTATCCCCTGTTTGGTATGGCGTTTCTCGTAATTCAGTCCGTATTCCTTCAGCATCACCGGCAG
CCCCAGCCCGAACATTTTCAGACTGAGTACATTCCGGTAGCCGTTTGCCTCCATGTAGGCCAGATAGGCGTGATAGAGGT
ATTTACGGTAATTGCGCGGGATGATACTGGCGTTCCCCATATACATGCCGCTGGTCTGCGGCAGGGTTTCCAGATAGCCG
ATAAAATCAAACGTCGGGTCGGCATCCCGTTTGATGTTCAGTGCCTCGTCTGAGTTCTGCTGGGACTGAAGCAGTGACCG
GGCGAGCATCGGGTCGCTGAACTTCTGCATCAGGTGACGCACGATGACCGCCAGCTCGCGGGTGATTTTGTCCTTAAGCT
GCGGGTCGCGCTCCTGCGGGGCTATCTGTTCCGGGAAGTGAATAATCACCCGTCGGCGTGACACGCCGCCGCTGCGGTCG
GTGAAGCGCATCGGGTTATTGTTCACGGCCAGAATCACCGCCGGGATGTGCGTGGAGTACGCATCCCGGTATTTCGGGTC
AACGGACACCGCATCGCCGCCGGTGATGGCCTTGAGTCCGGCACCGTCGCCGCTCCATTTTTCCTGGTCCGGCAGGCGTA
```

```
TCAGTGAGAAGCCAGTTAACGCGGCACGTTCACGCGGGGATTCCAGCGTCTCGATGGTGGCCGACGTGGCGTTATCCTCC
CCGGCCAGCAGGGTGGCTATTTCGGCCATGATACTTTTGCCGCTGCCGCCGGGACCGGTCACCTCCAGAAAGAGCTGCCA
GTCGTAGCGGTTTGCCAGCACCATAAACAGTGCAGCCAGAATCACGTCGCGTTTTTCCGCACGGCCACCGGCGGCACGGT
CAAGCCAGCGCCAGAAGGCGGGGGCGTGGGTTTCCAGCGTTTCACCGTCCACCGGCGGGGTGAAATCCACATCGCACAGG
GTGCGCATCCAGTGTGACGGACTGTGCGGGTGGAACGTGCCGTTCTGCGTGTCGAGCACGCCGTTACGAAAGCCAATCAG
GCGGCGGGAGGGGGCTTCCTGCTGCGGAATAATCAGCTTCAGGGTGTCCACCACGGAGGCCACCTTCCCGGAGGAGAACG
GCGCACGCAGACGCTGAAACAGCCCGGCCACATCCCGGGCAAAGTCCTGTGGCGGCAGCACCTTCCAGACACCATTTTCA
TAGCGGGACAGAAGCTGGCCGTTGGCATCGACCGCGAGCGCCTCGCCGTAATGCTCATAGATACGCATGGCCTTTTCGCT
GGTACTCATGGCGGAAAACTCCGCTTCGCTCATGGTGTCGAACGGGCTTTCAGCCGGTGGCCGGATGGCATCGTAAATGG
CCTTACGGGTGGCCTCCCCGCCGTACTGCGTGAAGGCATCATTCCAGTCACCGAAGACCGGCGGCAGGGCAACAACACCT
TCACACGCATCTGCGGCTGCGGCGGCTTTTTTCTGGCCGTCACCACTGAGGTCACGGTCAGCGGCAAGGACAATCTGACA
GGCGGGGTGCTTCTGCCGGGCAAGGCTGGCCAGAGAAAGGAGGTTCACGGAAGAAAGCGCCACCATCACCGTTTCACCGG
TCAGGTGATGTACGGTAAGTGCGGTCGCGTATCCCTCCGCTATCCACAGACGTTTTCCGGCCTGATTCTGTCCTTCAAGG
GTGTGACAGGTGCCCCTGACCTGTCCGCCTTTCAGGGTGCGCTTACGGCCGTCAGCACTGATTAACTGAAGGTTAACCAG
TTCGCCGCTGTCGTCATACAGTGGCACCACAAGGTCACCGGCGCGCCAGCTCACGCCACCGGCTCTGTGTGTGCCGGTCA
GCATCCGGCATTCCCGGCCGGGAAAGCCCTTGCGGGTCAGGTAGGCGTTACCGGTTCCGGTACGGGTTTTCGCCATCAGG
GTTTGTGCCAGTGCGGCGGCGTTCTTCCGGGCAGCGTCTGTTTCATCAACGGCGGCGGTCGTCACTGCCGGGTCAGCCGG
TGGCAGGCTGCCGGTCACGGCAGCCACCTTTGCGGCCGCGTCGGACGGGGAAACACCAAAAACCTTTTCAACCAGTTTCA
GGCCGTCACCGGCACCACACTGATTGCAGTACCAGGTGCCGCGCCCCTCCCTGTCATCAAAACGGAAGCGGTCACTCCCG
CCACAGACCGGACAGGGCTGATGACGGTTCTTCAGCACCTGAATCCCCAGCGCCGGGAGAATACGCGGCCAGTGGCCGAG
CGCATGGCTGACGGTGGCGGTTACGTTCATTTTCATGGTGTTGTTCTCCTTCAGTGCAGTACCGGCGCTTTTATGTGACG
GGCACAGAGTTCATCCATCACAACCAGCCCGAGAAAGGACAGCGACGGCGCGGCCTTCAGGGGGCCGGATTCCATTAAAT
CTTCCAGCAGGGCACAGGCTATCTGACGCCCTTTTTCCTCACCGTGCTGGCGCAGATAAAAGCCTTCCAGCTCAGCGGCG
ATGGCCGCCTCCAGTGACTCAAGGGTGAGATGCGGGTAGCGGTGCTGACGTTCGCACACGGTCAGCCAGGCACAGGCGAC
AGCGCGACGGTAAAGGGCAGCGCGTAAGACGGGCGGTAAGGGTGTTTTCATTTGCTTTTCTCCCTGTGACAGATGACTGC
ATTCCGTGCCGGTTGCATTAACTGATAAGGCATATCTGCGTCTCCTGAAGACGTGCGTATCCCTGCGCGAATACGCACAT
TTAATTTTTCGGGGGTCGTTTTTTAATTACAGATAATTGCGGTAACTGTTATCCGGGGTGGTTTCCGGGTCAGGCTCCGT
GCGGGGAATTTCCCGCCATTCCCGCGCCACCGGTGCTGCCCGGCTGACCGGAACAGTGTCCTGCGGGTAAATATCCAGAT
ATTTTTCCCGCCATTTCTGTAATTCCGGGTCTCCGGCCATTTCTTTCAGTACCGCATGCCGGTTTACGGGGCTGCGTTTA
AACAGGTCAGGACGGTCACAGGTAAATTCCCGCAGAAAACGCCCCAGCGGGATGTCTGTGGTGCGTCCGTCAGCGAGGAT
ACGCACAAGGATACTGAATTTACGGCGGTACGGGTTCCAGACAATGTCCGGGCAGCGGTACGGCATTTCCCACGGAATAC
CGTCTTCCAGAATGCCGACCACGGCCACATCGGGAAAACCGGCAGAACGGTAAATCTCACCGGGCTGGGGAAAATCAAAC
ATGCGTCCTGTCTCCCCGGTCTTTCTGCTGGGCGAGAAAATCGCGGCACAGGCCTTTGGCTTTCAGCTCATTCAGCACAA
AATCAATATCTTCATTCAGGTAGCTGAAAATATGTGGAATGTAGAGCTGATGCAGGCCGGAGAGTTCACGGTGAATCAAA
TCACCCCCAACAAACCGGGATACGGCGCTGGCGCGGTTGAGCTTATGGTAAGCCTCAATGCTGAGGTGTTCACGGGCGTC
ATGACGCGCTGAGACGGTCTGAGGGGCTTTTTTATTACGCACGGGACACCTCCACCACCGGCAGACGGGCAGCAAGGGAG
AGCACATAGTCACGGACAAGGGAACGGCGGGCACTGCGTTCATCACCGGCGACGGTGCGAAGCATACAGATACGGGGATG
ACGGTCTGCGCGACGGACAGCCGCAAACACAAAGACAAATTCAGGGTGTGAGGGGGTAAGGGTTGTAGCCATGATGGCAG
CCTCCTGTGAATAGCAAATAACGCTATCGCCGGAGTTCTCACGCTCGATGGCGATAGCCCAGACGGGGGTGAGAATACCG
GCTTCACAGGATACCGGCCAGCCCGGAGGCTGCCCCGCCTGAGCTACCATTGACTCTGCGGCATAATGAGCGGACGCGGG
CAGGATGCACGGAATGCCATCTGCACGACTGACCACACACCACACCATAATCTGGCGCTCTGTGGCATTGATTGCGACAC
AAAAAAAGACGCGTGGCGCGTCATATGTCGCCTGTGAATTGCTCGGGTTCTCACGCCCGGCTGCCGATTTTGCGGCAGCG
GAAAAACTATATCCGCAAATGCCGGAAAAAGGCAAGCCAGAAAAAGGGAGTTTTTGCAGAGCGGGCATCATCATGCGTCG
TACCCCCGTTTGCGTCCGGCAATGCGTCCGGCCATCCATGCGGTGACTTCAGAGTGCAGCCAGGCCACATTTTTACCGCC
AAGACTCACCTGCGGCGGAAATTCCCCCTTACGGATGAGTTCGTAGATGGTCGAGCGTGACAGGCCGCACAGGTGCATCA
CTTCCGGCAGACGTAAAAAACGCTCCTGCGTGATGTCCGGCAGCGGCATCAGTGGCGTCACTGGGGCGGGAGACGGGGAA
GAAAAAACAGCTTGCATCGGGCTACCTCGTTAATGTCCATACAGCACCGGATAAGTCCGTCCGGCTTCGGGTAGCGCTTT
ATTTTATGAATATTTTCAGCAGACGCAACAGGGGGGATTTGTTCAGGCTGTCTTACAATGGCTGTGTGTTTTTTGTTCAT
CTCCACTTAAAGTCATTTAAAGCCACTTAAAGCAATTTGTAATTTTTATAGTGAAATACAAATCGTTTCTTCTTATTCAT
TCCCGGCGAATTAATAAAAACAAACAGTAGTAAACAGCACAAAAAGCCCATCAACGGGTGAACAGTGGTGAACAGACGGT
GAACAGTCATTACTGCGATTGTTCACCCTTTAACTTACTGTATTACTTATCTTTTTTATTAAGGTGAACAGAGGTGAACA
GTAAAATATAAAAAAACAAACAGTAAGCCGGTTTTTCCTGCGACCTTTTCCTGGCTTGCCGGTCTGAGGATGAGTCTCCT
GTGTCAGGGCTGGCACATCTGCAATGCGTCGTGTTGTTGTCCGGTGTACGTCACAATTTTCTTAACCTGAAGTGACGAGG
AGCCGGAAAATGTCTGACCACACTATCCCTGAATATCTGCAACCCGCACTGGCACAACTGGAAAAGGCCAGAGCCGCCCA
TCTTGAGAACGCCCGCCTGATGGATGAGACCGTCACGGCCATTGAACGGGCAGAGCAGGAAAAAAATGCGCTGGCGCAGG
CCGACGGAAACGACGCTGACGACTGGCGCACGGCCTTTCGTGCAGCCGGTGGTGTCCTGAGCGACGAGCTGAAACAGCGC
CACATTGAGCGCGTGGCACGCCGGGAGCTGGTACAGGAATATGACAATCTGGCCGTGGTGCTGAATTTCGAACGTGAACG
CCTGAAAGGGGCGTGTGACAGCACGGCCACCGCCTACCGGAAGGCACATCATCACCTTCTGAGTCTGTATGCAGAGCATG
AGCTGGAACACGCCCTGAATGAAACCTGTGAGGCGCTTGTCCGGGCAATGCATCTGAGCATTCTGGTACAGGAAAATCCG
CTCGCCAACACCACCGGCCATCAGGGCTACGTCGCACCGGAAAAG
```

**Supplementary Table S22. Full plasmid sequence from the source map for pP4-LacIq-p72 (pAJ1058; 9,381 bp).** Source file: pAJ1058_pP4-w:o-p72-correct-size-bound_Warwick4-f7.dna. Feature key: P4 pLE promoter; P4 cI; P4 epsilon; P4 alpha; P4 alpha terminator; P4 crr replication region; kanamycin resistance gene; kanamycin-resistance promoter; lambda t0 terminator; T7 terminator; L3S2P21 terminator; PE antigen; 6xHis tag; P4 *sid* promoter; P4 pLL promoter

```
TATAGTTTTTCCGCTGCCGCAAAATCGGCAGCCGGGCGTGAGAACCCGAGCAATTCACAGGCGACATATGACGCGCCACG
CGTCTTTTTTTGTGTCGCAATCAATGCCACAGAGCGCCAGATTATGGTGTGGTGTGTGGTCAGTCGTGCAGATGGCATTC
CGTGCATCCTGCCCGCGTCCGCTCATTATGCCGCAGAGTCAATGGTAGCTCAGGCGGGGCAGCCTCCGGGCTGGCCGGTA
TCCTGTGAAGCCGGTATTCTCACCCCCGTCTGGGCTATCGCCATCGAGCGTGAGAACTCCGGCGATAGCGTTATTTGCTA
TTCACAGGAGGCTGCCATCATGGCTACAACCCTTACCCCCTCACACCCTGAATTTGTCTTTGTGTTTGCGGCTGTCCGTC
GCGCAGACCGTCATCCCCGTATCTGTATGCTTCGCACCGTCGCCGGTGATGAACGCAGTGCCCGCCGTTCCCTTGTCCGT
GACTATGTGCTCTCCCTTGCTGCCCGTCTGCCGGTGGTGGAGGTGTCCCGTGCGTAATAAAAAAGCCCCTCAGACCGTCT
CAGCGCGTCATGACGCCCGTGAACACCTCAGCATTGAGGCTTACCATAAGCTCAACCGCGCCAGCGCCGTATCCCGGTTT
GTTGGGGGTGATTTGATTCACCGTGAACTCTCCGGCCTGCATCAGCTCTACATTCCACATATTTTCAGCTACCTGAATGA
AGATATTGATTTTGTGCTGAATGAGCTGAAAGCCAAAGGCCTGTGCCGCGATTTTCTCGCCCAGCAGAAAGACCGGGGAG
ACAGGACGCATGTTTGATTTTCCTCAGCCCGGTGAGATTTACCGTTCTGCCGGTTTTCCCGATGTGGCCGTGGTCGGCAT
TCTGGAAGACGGTATTCCGTGGGAAATGCCGTACCGCTGCCCGGACATTGTCTGGAACCCGTACCGCCGTAAATTCAGTA
TCCTTGTGCGTATCCTCGCTGACGGACGCACCACAGACATCCCGCTGGGGCGTTTTCTGCGGGAATTTACCTGTGACCGT
CCTGACCTGTTTAAACGCAGCCCCGTAAACCGGCATGCGGTACTGAAAGAATGGCCGGAGACCCGGAATTACAGAAATG
```

GCGGGAAAAATATCTGGATATTTACCCGCAGGACACTGTTCCGGTCAGCCGGGCAGCACCGGTGGCGCGGGAATGGCGGG
AAATTCCCCGCACGGAGCCTGACCCGGAAACCACCCCGGATAACAGTTACCGCAATTATCTGTAATTAAAAAACGACCCC
CGAAAAATTAAATGTGCGTATTCGCGCAGGGATACGCACGTCTTCAGGAGACGCAGATATGCCTTATCAGTTAATGCAAC
CGGCACGGAATGCAGTCATCTGTCACAGGGAGAAAAGCAAATGAAAACACCCTTACCGCCCGTCTTACGCGCTGCCCTTT
ACCGTCGCGCTGTCGCCTGTGCCTGGCTGACCGTGTGCGAACGTCAGCACCGCTACCCGCATCTCACCCTTGAGTCACTG
GAGGCGGCCATCGCCGCTGAGCTGGAAGGCTTTTATCTGCGCCAGCACGGTGAGGAAAAAGGGCGTCAGATAGCCTGTGC
CCTGCTGGAAGATTTAATGGAATCCGGCCCCCTGAAGGCCGCGCCGTCGCTGTCCTTTCTCGGGCTGGTTGTGATGGATG
AACTCTGTGCCCGTCACATAAAAGCGCCGGTACTGCACTGAAGGAGAACAACACCATGAAAATGAACGTAACCGCCACCG
TCAGCCATGCGCTCGGCCACTGGCCGCGTATTCTCCCGGCGCTGGGGATTCAGGTGCTGAAGAACCGTCATCAGCCCTGT
CCGGTCTGTGGCGGGAGTGACCGCTTCCGTTTTGATGACAGGGAGGGGCGCGGCACCTGGTACTGCAATCAGTGTGGTGC
CGGTGACGGCCTGAAACTGGTTGAAAAGGTTTTTGGTGTTTCCCCGTCCGACGCGGCCGCAAAGGTGGCTGCCGTGACCG
GCAGCCTGCCACCGGCTGACCCGGCAGTGACGACCGCCGCCGTTGATGAAACAGACGCTGCCCGGAAGAACGCCGCCGCA
CTGGCACAAACCCTGATGGCGAAAACCCGTACCGGAACCGGTAACGCCTACCTGACCCGCAAGGGCTTTCCCGGCCGGGA
ATGCCGGATGCTGACCGGCACACACAGAGCCGGTGGCGTGAGCTGGCGCGCCGGTGACCTTGTGGTGCCACTGTATGACG
ACAGCGGCGAACTGGTTAACCTTCAGTTAATCAGTGCTGACGGCCGTAAGCGCACCCTGAAAGGCGGACAGGTCAGGGGC
ACCTGTCACACCCTTGAAGGACAGAATCAGGCCGGAAAACGTCTGTGGATAGCGGAGGGATACGCGACCGCACTTACCGT
ACATCACCTGACCGGTGAAACGGTGATGGTGGCGCTTTCTTCCGTGAACCTCCTTTCTCTGGCCAGCCTTGCCCGGCAGA
AGCACCCCGCCTGTCAGATTGTCCTTGCCGCTGACCGTGACCTCAGTGGTGACGGCCAGAAAAAAGCCGCCGCAGCCGCA
GATGCGTGTGAAGGTGTTGTTGCCCTGCCGCCGGTCTTCGGTGACTGGAATGATGCCTTCACGCAGTACGGCGGGGAGGC
CACCCGTAAGGCCATTTACGATGCCATCCGGCCACCGGCTGAAAGCCCGTTCGACACCATGAGCGAAGCGGAGTTTTCCG
CCATGAGTACCAGCGAAAAGGCCATGCGTATCTATGAGCATTACGGCGAGGCGCTCGCGGTCGATGCCAACGGCCAGCTT
CTGTCCCGCTATGAAAATGGTGTCTGGAAGGTGCTGCCGCCACAGGACTTTGCCCGGGATGTGGCCGGGCTGTTTCAGCG
TCTGCGTGCGCCGTTCTCCTCCGGGAAGGTGGCCTCCGTGGTGGACACCCTGAAGCTGATTATTCCGCAGCAGGAAGCCC
CCTCCCGCCGCCTGATTGGCTTTCGTAACGGCGTGCTCGACACGCAGAACGGCACGTTCCACCCGCACAGTCCGTCACAC
TGGATGCGCACCCTGTGCGATGTGGATTTCACCCCGCCGGTGGACGGTGAAACGCTGGAAACCCACGCCCCCGCCTTCTG
GCGCTGGCTTGACCGTGCCGCCGGTGGCCGTGCGGAAAAACGCGACGTGATTCTGGCTGCACTGTTTATGGTGCTGGCAA
ACCGCTACGACTGGCAGCTCTTTCTGGAGGTGACCGGTCCCGGCGGCAGCGGCAAAAGTATCATGGCCGAAATAGCCACC
CTGCTGGCCGGGGAGGATAACGCCACGTCGGCCACCATCGAGACGCTGGAATCCCCGCGTGAACGTGCCGCGTTAACTGG
CTTCTCACTGATACGCCTGCCGGACCAGGAAAAATGGAGCGGCGACGGTGCCGGACTCAAGGCCATCACCGGCGGCGATG
CGGTGTCCGTTGACCCGAAATACCGGGATGCGTACTCCACGCACATCCCGGCGGTGATTCTGGCCGTGAACAATAACCCG
ATGCGCTTCACCGACCGCAGCGGCGGCGTGTCACGCCGACGGGTGATTATTCACTTCCCGGAACAGATAGCCCCGCAGGA
GCGCGACCCGCAGCTTAAGGACAAAATCACCCGCGAGCTGGCGGTCATCGTGCGTCACCTGATGCAGAAGTTCAGCGACC
CGATGCTCGCCCGGTCACTGCTTCAGTCCCAGCAGAACTCAGACGAGGCACTGAACATCAAACGGGATGCCGACCCGACG
TTTGATTTTATCGGCTATCTGGAAACCCTGCCGCAGACCAGCGGCATGTATATGGGGAACGCCAGTATCATCCCGCGCAA
TTACCGTAAATACCTCTATCACGCCTATCTGGCCTACATGGAGGCAAACGGCTACCGGAATGTACTCAGTCTGAAAATGT
TCGGGCTGGGGCTGCCGGTGATGCTGAAGGAATACGGACTGAATTACGAGAAACGCCATACCAAACAGGGGATACAGACC
AACCTGACGCTGAAAGAGGAAAGCTACGGCGACTGGCTGCCAAAATGTGACGACCCTACAACAGCCTGACCCACCTGACC
GGCATCTGCCGGTCTTTTTTTATCCCGATATCCCCCGAAGGTGAACAATCCACTGTTCACCCTTCACCGTATATTCACCC
GTTATCACACTGAAATTAAAAGAGAAAAACGAAAGGTGAACAGTGTGAACAATCAAATCAAAAAAAACTTTTTTCTTCCT
GAGTGATTTCAGTGCGGAGGATTAATCACCGGTATGAGTCACATCGGCAGAATGCCGGAGGTGAAGAATCGAATGTTCAC
CCTTCACCCATTATTCACCACCTATCACTCTGAAATAAAAGGAGAAAACAGAAAGGTGAACAGTGTGAACAGTTCTTTCG
AAAAAAAATTTTTTATGTGGATAAAAGATGCACTGGTTTGGATCCGAGCTACGGATCCAAACTTACTGATTAAAAATGAT
TTAACTTTCCGATTTATTGATGGGCTGAAAATAGATTGGTCAGAAAAACAGTGGGGGCACAAAAGGGGGCATATTTCATT
GTTGTGTTTTATTATTTGTTTTTTATCATTAACTTATTTTGATGTTTGAGTCCGGCCTTCGGCACCATTCTTACTTCCAA
GTTAGAAAAATTCATCCAGCATCAGATGAAATTGCAGTTTGTTCATATCCGGATTATCAATGCCATATTTCTGAAACAGA
CGTTTTTGCAGGCTCGGGCTAAATTCGCCCAGGCAGTTCCACAGAATGGCCAGATCCTGATAACGATCCGCAATGCCCAC
ACGGCCCACATCAATGCAGCCAATCAGTTTGCCTTCATCGAAAATCAGGTTATCCAGGCTAAAATCGCCGTGGGTCACCA
CGCTATCCGGGCTAAACGGCAGCAGTTTATGCATTTCTTTCCACACCTGTTCCACCGGCCAGCCGTTACGTTCATCATCA
AAATCGCTCGCATCCACCAGGCCGTTGTTCATACGGCTCTGCGCCTGGGCCAGACGAAACACACGATCGCTGTTAAACGG
GCAGTTGCACACCGGAATGCTATGCAGACGACGCAGAAACACGGCCAGCGCATCCACAATGTTTTCGCCGCTATCCGGAT
ATTCTTCCAGCACCTGAAACGCGGTTTTGCCCGGAATCGCGGTGGTCAGCAGCCACGCATCATCCGGGGTGCGAATAAAA
TGTTTAATGGTCGGCAGCGGCATAAATTCGGTCAGCCAGTTCAGACGCACCATTTCATCGGTCACATCGTTCGCCACGCT
GCCTTTGCCATGTTTCAGAAACAGTTCCGGCGCATCCGGTTTGCCATACAGACGATAAATGGTCGCGCCGCTCTGACCCA
CGTTATCACGCGCCCATTTATAGCCATACAGATCCGCATCCATGTTGCTGTTCAGACGCGGACGGCTACAGCTCGTTTCA
CGCTGAATATGGCTCATAACACCCCTTGTATTACTGTTTATGTAAGCAGACAGTTTTATTGTTCATGATGATATATTTTT
ATCTTGTGCAATGTAACATCAGAGATTTTGAGACACAAATTTAAATCGTAATTATTGGGGACCCCTGGATTCTCACCAAT
AAAAAACGCCCGGCGGCAACCGAGCGTTCTGAACAAATCCAGATGGAGTTCTGAGGTCATTACTGGATCTATCAACAGGA
GTCCAAGACTAGTCGCCAGGGTTTTCCCAGTCACGACGCGGCCGCAAGCTTGCATGCCTGCAGCGGCGGCTGCTAGCATT
AACTTTGATTCCATTCTTTTGTTTGTCTCAAAAAACCCCTCAAGACCCGTTTAGAGGCCCCAAGGGGTTATGCTAGGGAC
CAAAACGAAAAAAGGCCCCCCTTTCGGGAGGCCTCTTTTCTGGAATTTGGTACCGAGCCCGGGTTATTATTTTTTATCCA
CGCTAAAGGCTTCGCCATAGTTAGCGCAAATGATCTGGGCGGCATTGTATAAAGTGGTATCCGGGGTCAGAGATAAAGTG
TGTTTCTTCTGCTTTTTCGGTGCGGCACGCAGACCTTGGCCCCAAAACTCATCGTAGAAATCGGTGCGAACTTGGGTCAG
ATGGTAGTTGGCGCAGTTCAGAATTTTATACTGACGAACGCTGCGGGCGTAACGTTTCGGCTCCGGATACACGTATAAAC
CTTTGTCTCGAGTTATTAATGATGGTGGTGATGATGACCACCAGGATCAGGATACTGGGCCATGGCCTCAACCAGCTTCT
CGATGCCGGCCTGATCCACTGCCTGGTTTGCTGCATGGATAATCTCCACGCGATGATCGGCATCGCGGTACCAATCATGC
ACTGTTAAGGCATCGTCTGTGCCCAGGATACGAATCTCCAGATCGTTACCCTGGCGGGCAAACCACAGCTGATCGGCGCC
TGCATTGATGCGGATTGTGTCGTGACCACCACCGCTCTCATAAATGGTGTCGTGGCCATAACCCACGCCAAACAGGTAGG
TGTCATCACCCTGGCCGCCGAATAAATCGTCATTGCCGGCGTCACCGCTCAGTAAGTCACTGCCCTCATCACCCAGCAGA
ACATCATCACCTGCACCACCACTCAGCACGTCGTTGCCGGCCAGGCCGTTCAGAACATTTGCACCGGCATCACCGATCAG
CACGTCATCGCGGGCACTACCCACGGCGTTCTCAATGTTGCGTAACACGTTGCTGGTTTCCGGTTCGTCCATTAAGCTGG
CAAAGCCTTTGCCCAGGCTCAGAAACACACCCTTTGCGCCGGCATCCAGGCCACGACCAGGACCGCTAAAATCCACTGTA
TCACGACCGTCGCCACCGTCTAACAGATTACCGGCATTGGCTGCGTCCTGAAAGAACCAATCATCGCCTGCTTCGCCATA
CAGGCGGTCGCGACCTGCATCACCACTCAGCTGATCATCGCCGTCACCGCCCAGCAGAACGTCATCACCCTCACCACCGG
CTAACACATCGGCACCACCTGCGCCGCGCAGAACATTGGCTTGGGCATCGCCTGTGATACGGTCTGCCAGCTCTGTGCCC
ACCACGTTTTCGATGCCGCTCACTGTGTCATAGGCACTACTGCCGGCCTCACCCAGTTTATCCACACGACCGGCACCCAG
GTCTGCCAGAATACCCAGACCAATGCGACCGGCTGCGATACCTGCATGGGCGCCTGTCTGACTGTAGTCAACGGTATCCA
CGCCATCGCCACCGCGCAGCACATCATGTTCACGGGCCTGGGTTTGGCCAAACCAATCGTTGCCGGCACCGCCCTCCAGG
GTATCGTCACCCAGACCGCCGTACAGGGTGTCGTTACCGGCATCGCCATACAGCATGTCGTTACCATCGCCGCCGAACAG
CAGGTCATCACCATCACCACCGCGCACTGTATCGTCACCACCCTGGCCCATCAGGGTGTTGGCCTGGGCGTCGCCAATCA
GCACGTCTTTCATGCTGGTGCCAATCACGTTTTCCACATTGCGCACGTTGTCGTAATAGTCCACACCCAGGGCACTTGCT
TTGCGCACCCATTCACGGCTCAGATCGGCTTCGATACCAAAACCATATTCGTGCGGGGCCACAATACGGCCCGGATGAAT
CATGGCGCTGTAGTCCACTGTGTCCAGACCTGCGCCACCGTCAATGTCGTCGCTAACTGTTTCGTCATCCTGCAGAAAGA
TATCATTGCCATCCTCACCGTACAGTGTATCCAGACCCAGACCACCACGCAGAATGTCGTCGCCACCACGGCCGCGAATG
GTGTCATTACCGTCATGGCCCCACAGTTCGTTATCTTGGTCATCGCCGGCAATGCGGTCGTTCAGACGGCTGCCATGCAG

```
ATTTTCAATATTTTTAACATGATCCACGCTAAACAGGTTCAGGGCCGGCCATTTTTCCACGGTGCCTTTCTGCAGATCGG
CATGAATACCGGTGTCACCCATGCGTTCCAGACGTTCCTCGCTCGGCTGATGAACGTTGTATTTCACGGTGTCCACACCT
GCACCACCGTCCAGCTGGTTGCTCCACACGCCCAGATCCTGCAGGAAAACATCGTCGCCGGCACCGCCAATAACGGTATT
TTGACCCTCGCCACCCACCAGGGTATCATTACCGGCGCCACCATCCAGACGATCATCGCCGCTGCCGCCTGCCAGAAAGT
TGTCGTGGGCGTTGCCTGTAATGCTGTCATTGCCGGCACCGCCAATAATATGCTGCACGTGTTCCAGGGTGTCCACCTCA
ACTAATTGGCCAACGCGTGCCAGTTCAACGTGGCGCTGTTTCAGCTCGTCGCTCAGGACACCACCGGCTGCACGAAAGGC
CGTGCGCCAGTCGTCAGCGTCGTTTCCGTCGGCCTGCGCCAGCGCATTTTTTTCCTGCTCTGCCCGTTCAATGGCCGTGA
CGGTCTCATCCATCAGGCGGGCGTTCTCAAGATGGGCGGCTCTGGCCTTTTCCAGTTGTGCCAGTGCGGGTTGCAGATAT
TCAGGGATAGTGTGGTCAGACATTTTCCGGCTCCTCGTCACTTCAGGTTAAGAAAATTGTGACGTACACCGGACAACAAC
ACGACGCATTGCAGATGTGCCAGCCCTGACACAGGAGACTCATCCTCAGACCGGCAAGCCAGGAAAAGGTCGCAGGAAAA
ACCGGCTTACTGTTTGTTTTTTTATATTTTACTGTTCACCTCTGTTCACCTTAATAAAAAAGATAAGTAATACAGTAAGT
TAAAGGGTGAACAATCGCAGTAATGACTGTTCACCGTCTGTTCACCACTGTTCACCCGTTGATGGGCTTTTTGTGCTGTT
TACTACTGTTTGTTTTTATTAATTCGCCGGGAATGAATAAGAAGAAACGATTTGTATTTCACTATAAAAATTACAAATTG
CTTTAAGTGGCTTTAAATGACTTTAAGTGGAGATGAACAAAAAACACACAGCCATTGTAAGACAGCCTGAACAAATCCCC
CCTGTTGCGTCTGCTGAAAATATTCATAAAATAAAGCGCTACCCGAAGCCGGACGGACTTATCCGGTGCTGTATGGACAT
TAACGAGGTAGCCCGATGCAAGCTGTTTTTTTCTTCCCCGTCTCCCGCCCCAGTGACGCCACTGATGCCGCTGCCGGACAT
CACGCAGGAGCGTTTTTTACGTCTGCCGGAAGTGATGCACCTGTGCGGCCTGTCACGCTCGACCATCTACGAACTCATCC
GTAAGGGGGAATTTCCGCCGCAGGTGAGTCTTGGCGGTAAAAATGTGGCCTGGCTGCACTCTGAAGTCACCGCATGGATG
GCCGGACGCATTGCCGGACGCAAACGGGGGTACGACGCATGATGATGCCCGCTCTGCAAAAACTCCCTTTTTCTGGCTTG
CCTTTTTCCGGCATTTGCGGA
```

**References:**


[1] D. N. P. Doan and T. Dokland, “The gpQ portal protein of bacteriophage P2 forms dodecameric connectors in crystals.,” *J. Struct. Biol.*, vol. 157, no. 2, pp. 432–6, Feb. 2007.

[2] G. E. Christie and R. Calendar, “Bacteriophage P2.,” *Bacteriophage*, vol. 6, no. 1, p. e1145782, 2016.

[3] C. R. Büttner, Y. Wu, K. L. Maxwell, and A. R. Davidson, “Baseplate assembly of phage Mu: Defining the conserved core components of contractile-tailed phages and related bacterial systems.,” *Proc. Natl. Acad. Sci. U. S. A.*, vol. 113, no. 36, pp. 10174–9, Sep. 2016.

[4] P. G. Leiman *et al.*, “Morphogenesis of the T4 tail and tail fibers,” *Virol. J.*, vol. 7, no. 1, p. 355, Dec. 2010.

[5] J. A. Lengyel, R. N. Goldstein, M. Marsh, “Structure of the Bacteriophage P2 tail,” *Virology*, vol. 62, pp. 161–174, 1974.

[6] K. L. Maxwell *et al.*, “Structural and functional studies of gpX of Escherichia coli phage P2 reveal a widespread role for LysM domains in the baseplates of contractile-tailed phages.,” *J. Bacteriol.*, vol. 195, no. 24, pp. 5461–8, Dec. 2013.

[7] J. L. Kizziah, C. M. Rodenburg, and T. Dokland, “Structure of the Capsid Size-Determining Scaffold of "Satellite" Bacteriophage P4.,” *Viruses*, vol. 12, no. 9, 2020.

[8] T. Dokland, B. H. Lindqvist, and S. D. Fuller, “Image reconstruction from cryo-electron micrographs reveals the morphopoietic mechanism in the P2-P4 bacteriophage system.,” *EMBO J.*, vol. 11, no. 3, pp. 839–46, Mar. 1992.
